# Manipulation of Bacterial ROS Production Leads to Self-escalating DNA Damage and Resistance-resistant Lethality for Intracellular Mycobacteria

**DOI:** 10.1101/2023.07.07.548098

**Authors:** Junfeng Song, Mengmeng Wang, Huanyu Tao, Anming Yang, Zhaohong Zhu, Silei Bai, Miaomiao Luo, Junpeng Xu, Xueke Liu, Yicheng Sun, Peilei Hu, Wing-Leung Wong, Feng Li, Yongheng Chen, Qingyun Cai, Hongke Liu, Sheng-You Huang, Zhi Su, Xinxin Feng

## Abstract

The high prevalence of drug resistance in mycobacteria calls for antimicrobial mechanisms that suppresses the development of resistance. As a structurally conserved multi-site bio-macromolecule, DNA is presumed to be an ideal candidate for such resistance-resistant drug target. However, survey of marketed and investigational DNA interactors indicates that they are not immune to resistance development. Here, we report our strategy to achieve real resistance-resistant DNA targeting by incurring “catastrophic” DNA damage with an organoruthenium-natural product hybrid. The dual-mode DNA damage, in the form of strong tri-valent binding and concomitant oxidative modification, is achieved by manipulating of bacteria’s native endogenous ROS production mechanism upon lethal stress (such as DNA binding). Such self-escalating DNA damage, together with precise targeting of intracellular bacteria via vacuole fusion, thus endows the hybrid’s resistance-resistant lethality against mycobacteria and *in vivo* efficacy in animal models.

## Introduction

Antibiotic resistance is an ever-deteriorating a global problem that continues to cause considerable morbidity and mortality worldwide. The seriousness of resistance issue is best represented in the case of mycobacterial infection,^1–3^ whose treatment already suffers from the extremely limited options,^1, 4^ due to the intrinsic tolerance toward drugs caused by their waxy outer membrane^5, 6^, the intracellular persisting lifestyle sheltering themselves from antibiotic attack, as well as the proneness to resistance development endowed by the intracellular persistence^6–10^. In fact, multidrug-resistant strains of tuberculosis substantially increase the rate of mortality when compared to drug-susceptible tuberculosis cases during anti-tuberculosis treatment (39% versus 6%)^11^. The high prevalence rate and lethality of multidrug-resistant tuberculosis^12, 13^ calls for the discovery of new antimycobacterial mechanisms, and more importantly, it explicitly challenges the standard medicinal chemistry paradigm of “single drug, single target, single effect” as “single” usually means higher resistance generation. The term “resistance-resistant antibiotics” thus emerges, emphasizing on multi-targeting at an array of essential physiological processes in the bacterial lifecycle^14–17^. The termed strategy is particularly important for mycobacterial infection treatment as it touches the root issue of resistance.

For the development of resistance-resistant antibiotics, DNA has been acknowledged as a promising bio-macromolecular target. DNA is naturally a structurally conserved multi-site target, as it not only provides multiple sites for non-covalent groove binders and intercalators^18–21^, but also chemical reactivities on all purines and pyrimidines that enable covalent modification by alkylators^20, 22^, reactive oxygen species (ROS)^23–29^, reactive nitrogen species^30–34^ and others (Figure 1a). In the first glimpse, such sequence-independent DNA targeting may seem to be a way to effectively suppress resistance, but takes a risk of simultaneously stressing eukaryotes by harming their DNA. The reality, however surprisingly, is the opposite in term of both resistance and toxicity. For the matter of biocompatibility, metronidazole, an approved DNA-targeting prodrug, showcases a possible mechanism of bacterial DNA specificity by being activated only in the strongly reducing conditions found in anaerobes and protozoans^30–34^. In addition to this marketed example, we and others have also demonstrated specificity toward bacterial DNA can be established through the selective introduction of compounds towards bacterial chromosome^35–37^, and the development of sequence-specific binders^38–41^.

**Figure 1.**
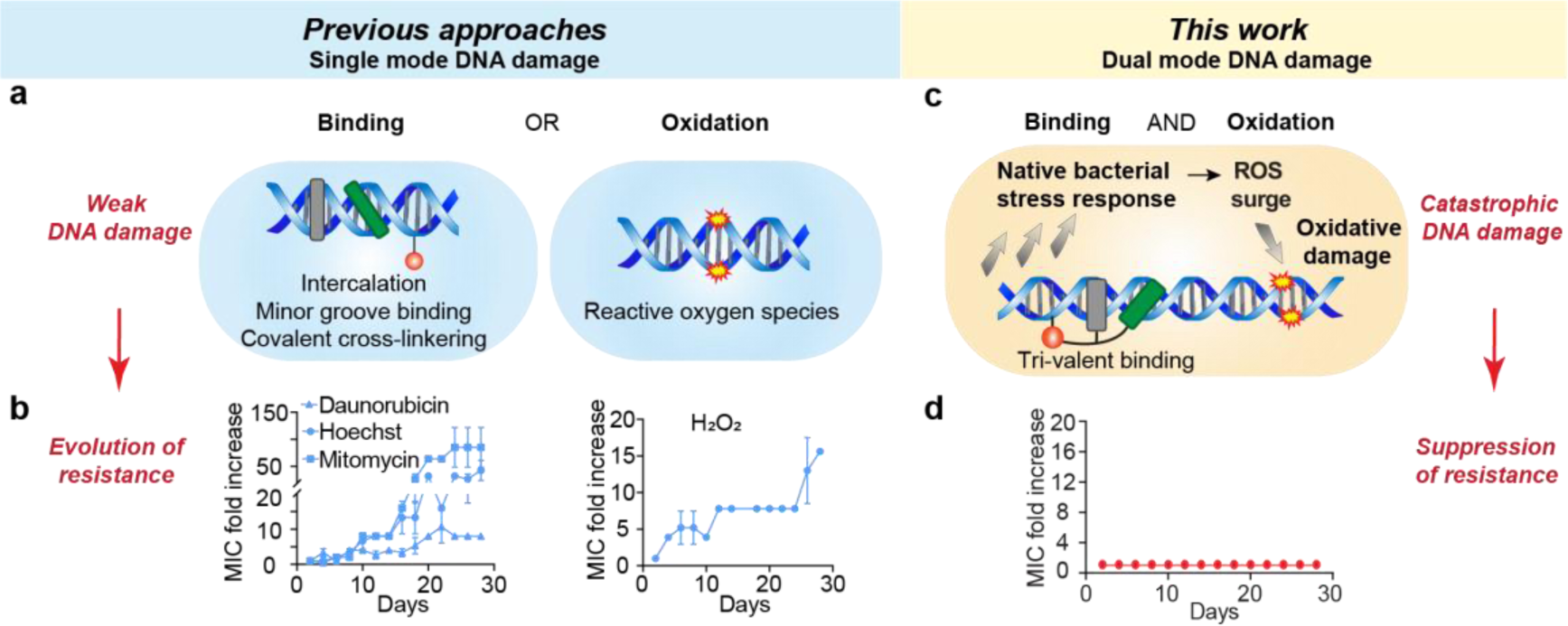
Comparison bewteen single mode DNA targeting mechanism and the proposed binding-oxidation dual mode DNA targeting. (a) Schematic representation of classic single mode DNA binding and oxidation. (b) Experimental resistance evolution profiles for Daunorubicin (an intercalator), Hoechst (a minor groove binder), Mitomycin (an cross-linker) and H2O2 (an oxidative agent). (c) Our proposed dual mode DNA damage by simultaneous multivalent DNA binding and oxidative modification as a strategy to suppress bacterial drug resistance. (d) Hypothetical resistance evolution profile for the proposed dual mode DNA damaging agent in (c).

On the resistance part, in contrast to one’s intuition, our survey of marketed and investigational DNA interactors on their resistance development revealed that DNA damage does not necessarily incur the phenotype of resistance suppression. For the abovementioned metronidazole, resistance strains have been identified^34, 42, 43^. Also, laboratory evolution has led to the emergence of resistant mutants for common DNA groove binder (Hoechst), intercalator (daunorubicin), DNA cross-linking agent (mitomycin) and ROS generating agent (H_2_O_2_) (Figure 1b). Alternation of DNA structure is apparently not a mechanism for such acquired resistance^19^. Indeed, the main modes of resistance development for those DNA-targeting compounds appear to be (1) the reduction of intracellular accumulation through either reduced influx or increased efflux, or (2) enhanced DNA-repair mechanisms^19^. Thus, the anti-intuitive resistance-prone nature of such DNA-targeting compounds may be rooted from the insufficient DNA damage they have been causing, which buys time for the bacteria to develop countermeasures in ways other than the mutation of the very drug target, i.e., DNA, leading to efficient emergence of drug resistance.

In this article, we report a strategy to incur “catastrophic” DNA damage to solve the resistance issue (Figures 1c-d). The self-escalating DNA damage, in the form of strong multivalent binding (covalent, intercalation and minor groove binding) and concomitant oxidative modification, is achieved by manipulation of the native bacterial stress-responding mechanism^44–48^ to overproduce ROS, which backfires major biomacromolecules including DNA (Figure 1c). In short, by constructing a sandwich organoruthenium-natural product hybrid, we achieved nanomolar level binding affinity to mycobacterial DNA with high degree of DNA structural alternation upon binding. Such strong perturbation of DNA function could lead to the overresponse of bacteria in the form of an ROS surge, incurring further and permanent DNA damage. This self-escalating DNA damaging through both multivalent structural alternation and oxidation leads to “catastrophic” DNA lesion, endowing the desired resistance-resistant lethality to mycobacteria. Favorably, this organoruthenium complex is also able to target and eradicate the intracellular mycobacterial persister by precisely targeting mycobacteria containing vacuoles (MCVs), allowing it to fully feature its potency and resistance-resistant nature in a mycobacteria infection animal model study.

## Results

### Designing a Ru^II^ complex with tri-valent DNA binding mode

To increase the binding affinity of a ligand to its target, multi-valence is a well-recognized strategy in both native biological system^49–51^ and synthetic binders^52–55^. We therefore aimed to build a tri-valent binding scaffold (i.e. intercalation, minor groove binding, alkylation) for DNA targeting, and we considered organometallic complex a promising starting point benefiting from its easy access to hitherto underexplored three-dimensional chemical space^56–59^. Metal complexes containing π-bonded arenes, such as Ru^II^-arene complexes of the half-sandwich “piano-stool” type, are chosen due to their known DNA intercalating and covalent binding activities, as well as other desirable characteristics, such as high aqueous solubility, clearance properties and low side effects^56, 60–62^ (Figure 2a, “Ru^II^-arene”). To complete the tri-valent design, DNA minor-groove binding ancillary group, natural products (NPs) are chosen as our candidate structures for their potential in DNA binding^38, 63, 64^ (Figure 2a, “Minor-groove binding natural product”). DNA-interacting NPs are mostly used as anti-cancer therapeutics^64–67^, but many of them enter the nucleus effectively as part of their essential working mechanism, which presents a toxicity concern for them serving as anti-infectives. Therefore, one must look for an NP with minor-groove binding activity but display low toxicity toward mammalian cells. With the above consideration, we established a focused library of NPs derived from traditional Chinese medicines. This library consisting of major classes of NPs, namely triterpenoids, alkaloids, anthraquinone, phenylpropanoids, flavonoids and polyphenols (Figure 2b and Supplementary Excel 1). To screen for such a minor-groove binding NP, a fluorophotometric competition assay with a known minor-groove binder Hoechst was employed to determine the percentage of Hoechst 33342 displacement (Figure 2c and Supplementary Fig. 1a-b). In the screening, NPs in the Rhein series, including Rhein, Emodin, and Aloe-emodin, exhibited the highest activities.

**Figure 2.**
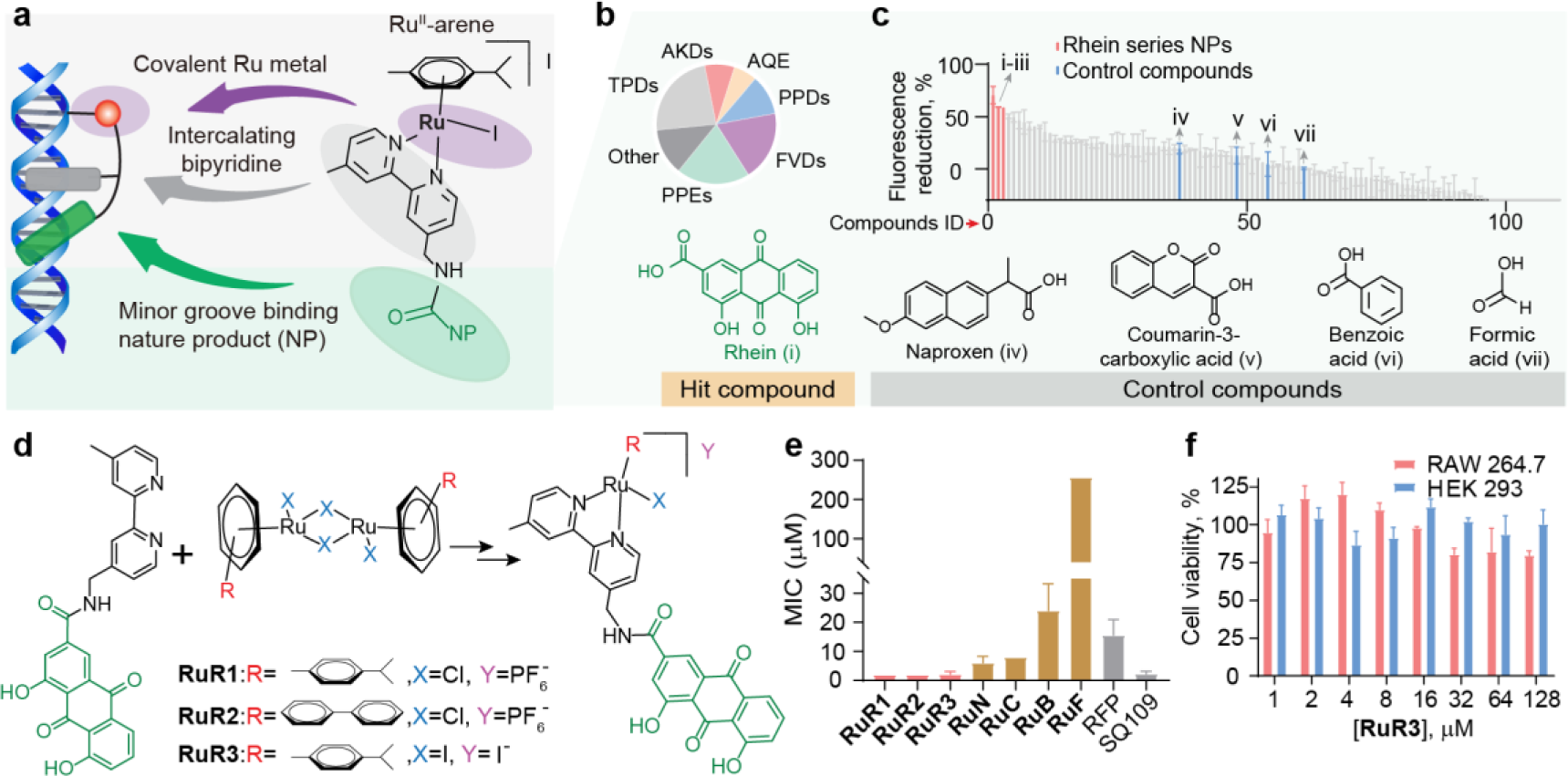
Design and synthesis of Ru^II^ arene-natural product (**Ru^II^-NP**) complexes, and these preliminary antimicrobial and cytotoxicity characterization. (a) Chemical structure of the **Ru^II^-NP** complex, and schematic representation of its multivalent DNA binding mode. (b) Composition of the natural products (NPs) library (TPDs: triterpenoids, AKDs: alkaloids, AQEs: anthraquinone, PPDs: phenylpropanoids, FVDs: Flavonoids, PPEs: Polyphenols). (c) Rank-ordered percentage of Hoechst displacement for all NPs in the library, and chemical structures of the selected minor groove binding ancillary groups. (d) Synthesis of **RuR**s and the chemical structures of **RuR1-3**. (e) MIC values of **Ru^II^-NP** complexes. (f) Cell viability assays of **RuR3** on two mammalian cell-lines (murine macrophage RAW 264.7 and HEK 293).

With the two building blocks chosen, we next synthesized a series of Ru^II^-arene-NP (**Ru^II^-NP**) complexes with the above NPs, via amide coupling reaction of Ru^II^-arene and NPs with carboxylic acid groups, and obtained **RuR**, **RuN**, **RuC**, **RuB**, **RuF** (Figure 2d and Supplementary Fig. 1c). A few other NPs with minor-groove binding activities, together with some simple carboxylic acids, were chosen as the control compounds (Supplementary Fig. 1a-b). For the complex with the best minor-groove binder hit, **RuR**, its arene group, ligand and counter ion was optimized to give **RuR1-3**. These compounds were characterized for ROS generating abilities (Extended Data Fig. 1a-b), finding that they are stable in varies physiological environments and are free of ROS production by themselves.

Preliminary antimicrobial test indicates that **Ru^II^-NP** complexes are active against a model Mycobacterial strain, *Mycobacterium smegmatis* (*M. smegmatis*), with **RuR1-3** exhibiting fastest and strongest bactericidal activities with minimum inhibitory concentrations (MICs) of 2 µM (Figure 2e and Supplementary Fig. 2a), and they also showed highest activity against pathogenic mycobacteria spp.^68–72^, including *Mycobacterium fortuitum* (*M. fortuitum*), *Mycobacterium marinum* (*M. marinum*), and *M. tuberculosis*, among all the candidates, with **RuR3** exhibiting the best overall activities (Supplementary Table 1). Toxicity and hemolytic activity evaluation showed that **RuR3** could preferentially kill bacteria over mammalian cells (Figure 2f and Supplementary Fig. 2b). Apparently, the tri-valent design was successful in term of cell culture performance, but the value of this design strategy can only be unveiled with further exploration in its mechanism of action as well as anti-resistance property.

### Evidence for multivalent DNA binding by RuR3 and its impact on DNA function

The multivalent DNA binding properties of **RuR3** and the downstream impact on DNA function was examined with a panel of biochemical experiments (Figure 3a). First, to quantitatively compare the intercalating- and groove binding-activities of **RuR3** and related compounds, fluorophotometric titration assays with Hoechst (a DNA minor groove binder) and EB (a DNA intercalator) were performed. **RuR3** displays stronger DNA binding affinity than both building blocks (Rhein, Ru^II^-arene) and classic single-valent DNA binders (Hoechst, EB), with *K_i_* (**RuR3** as groove binder) = 6.4 nM and *K_i_* (**RuR3** as intercalator) = 540 nM (Figure 3b-c and Supplementary Fig. 3). Additional evidence of DNA binding is provided by an electrophoretic mobility shift assay with plasmid DNA (Extended Data Fig. 2a) as well as an exogenous DNA interference assay (Extended Data Fig. 2b). Interaction between **RuR3** and DNA is covalent-based, as the adduct was clearly observed with ESI-MS (Figure 3d and Extended Data Fig. 2c). Note that, the strength of covalent binding was largely originated from the added minor-groove binding ancillary group (Rhein), since the yield of DNA-Ru^II^ arene adduct was significantly lower (Figure 3d and Extended Data Fig. 2c). This directly demonstrates the essentiality of proximity enabled covalent DNA modification.

**Figure 3.**
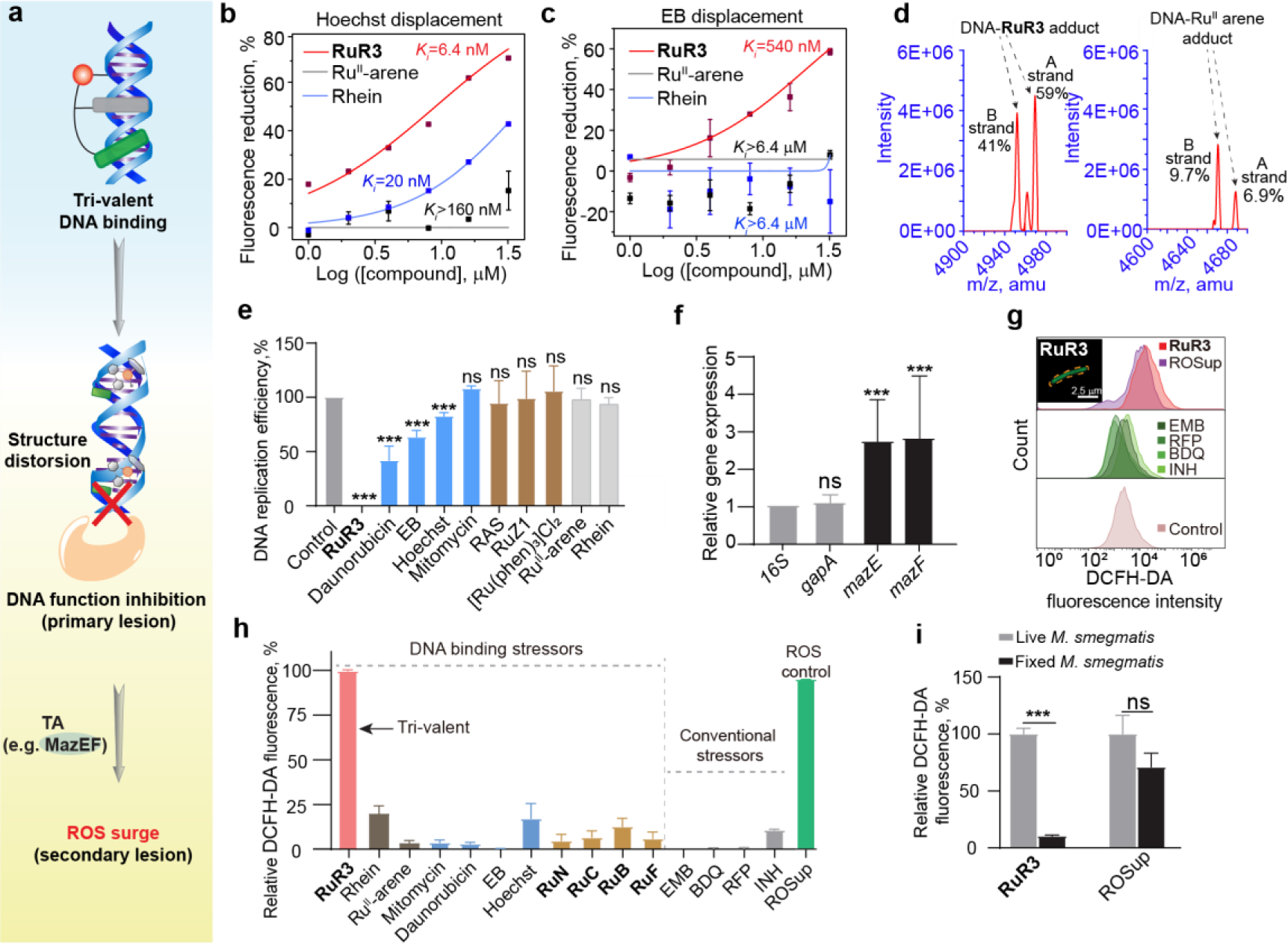
Evidence for multivalent DNA binding, DNA function interferencing and endogenous ROS production by **RuR3**. (a) Schematic illustration of tri-valent DNA binding by **RuR3** and the downstream effects on DNA function inhibition and endogenous ROS production. (b-c) Titration curves of **RuR3**, Ru^II^-arene and Rhein in Hoechst or EB displacement assays. (d) Negative mode ESI-MS spectrum of d(ATACATGGTACATA)2 treated with **RuR3** or Ru^II^-arene. (e) Effects of **RuR3** and various compounds on DNA replication. (f) Relative gene expression of *mazEF* upon **RuR3** treatments at 0.5 μM. Expression levels were normalized to that of the housekeeping gene *16S*. *gapA* is a control housekeeping gene that is not upregulated. (g) Confocal visualization and flow cytometric analysis of ROS accumulation in *M. smegmatis* upon treatment of **RuR3** and conventional stressors. ROS was probed by the green fluorescent dye DCFH-DA. (h) Comparison of ROS production levels upon compound treatments at 4×MIC. ROSup is a commercial exogenouse ROS producer. (i) Flow cytometry quantification of **RuR3**- or ROSup-induced ROS accumulation in live *M. smegmatis* and fixed *M. smegmatis*.

Overall analysis of the above DNA-binding assays revealed that, **RuR3** possesses higher affinity toward DNA when compared with classic single-valent DNA interactors (daunorubicin, EB, Hoechst, mitomycin), or other reported Ru-based metal complexes (RAS^73^, RuZ1^74^, [Ru(phen)_3_]Cl_2_). (Extended Data Fig. 2d, Supplementary Fig. 4 and Table 2). This affinity is presumably originated from the synergistic collaboration of the two building blocks, i.e., the intercalating/covalent Ru^II^-arene and the best minor groove binder from screening, Rhein. The above multivalent binding has led to strongest perturbation of DNA function, e.g. replication, as demonstrated with an *in vitro* PCR-based DNA replication assay (Figure 3e and Supplementary Fig. 5).

### DNA function perturbation induced an ROS surge in mycobacterium

Bacteria growing aerobically generate ROS via the endogenous ROS production system, as a metabolic byproduct to increase efficiency and yield of energy conversion from growth substrate^75^. It is known that a variety of lethal external stressors can stimulate an additional cascade of ROS production in bacteria via a genetic pathway of toxin-antitoxin modules (MazEF)^45^. Common stressors of ROS production include antimycobacterials that inhibit bacterial biomacromolecule synthesis^76, 77^, such as isoniazid for cell wall synthesis and rifampicin for RNA transcription. Here, we found that the tri-valent DNA binder and function perturbator, **RuR3**, exerts much stronger stress on the bacteria, as evidenced by higher level of *MazEF* activation (Figure 3f) and more abundant production of ROS upon **RuR3** treatment, as compared with reported common stressors (Figure 3g, Extended Data Fig. 3a-b and Supplementary Fig. 6). Further cross comparison of ROS production reveals that, **RuR3** treatment resulted in much higher ROS level than all control DNA binding stressors (Figure 3h and Supplementary Fig. 7), including (1) classic single valent DNA interactors (daunorubicin, EB, Hoechst, mitomycin), (2) building blocks of **RuR3** (Ru^II^-arene and Rhein), and (3) other **Ru^II^-NP**s with a weaker MGB ancillary group (**RuN**, **RuC**, **RuB**, **RuF**). The correlation between DNA-binding primary lesion and ROS generation was also examined, and as shown in Extended Data Fig. 3c, the degree of DNA binding affinity is highly correlated with the resulting ROS production, establishing the essentiality of multivalent DNA binding for effectively perturbating DNA function and resulting in ROS production. Next, we tried to examine the endogenous nature of **RuR3**-induced ROS. As we have previous shown in Extended Data Fig. 1a-b, the **RuR3** induced ROS surge was not from an exogenous source, since ROS was not detected by Electron Paramagnetic Resonance (EPR) or fluorescent probes when **RuR3** was mixed with DNA *in vitro*. The endogenous nature of **RuR3** induced ROS was further supported by the lack of intra-bacterial ROS accumulation caused by **RuR3** in fixed bacterial cells. In contrast, the ROS generation of ROSup, an exogenous ROS generator, remained unaffected in the fixed cells (Figure 3i and Extended Data Fig. 3d). This suggested that **RuR3**-induced ROS should be relying on the proper metabolic function of bacterial cells. High level of **RuR3** induced ROS products in turn triggered cellular response, in the form of up-regulation of ROS quenching enzymes such as AphC, SOD, KatG and OxyR (Extended Data Fig. 3e).

### Endogenous ROS surge results in oxidative damage of bacterial genomic DNA

There was direct evidence showing that the antimicrobial effect of **RuR3** was heavily attributed to ROS generation. For example, perturbation with reducing agent was shown to reduce the amount of endogenously generated ROS (Extended Data Fig. 3f) and partially rescue the bacterial cell death (Extended Data Fig. 3g). Also, as a result of the elevated intracellular ROS concentration, high level of DNA oxidative damages was expected in the form of base oxidation and strand breaks (Figure 4a). 8-Hydroxy-2 deoxyguanosine (8-OHdG) is the most common oxidative lesion observed in duplex DNA because guanine has a lower one-electron reduction potential than the other nucleosides in DNA^78, 79^, and it was indeed observed that the treatment with **RuR3** resulted in elevated levels of 8-OHdG on genomic DNA of *M. smegmatis* (Figure 4b). Importantly, this phenomenon was neither observed for genomic DNA from mammalian cells nor free DNA *in vitro* (Supplementary Fig. 8), indicating the inherent selectivity of this killing mechanism based on intracellular ROS generation.

**Figure 4.**
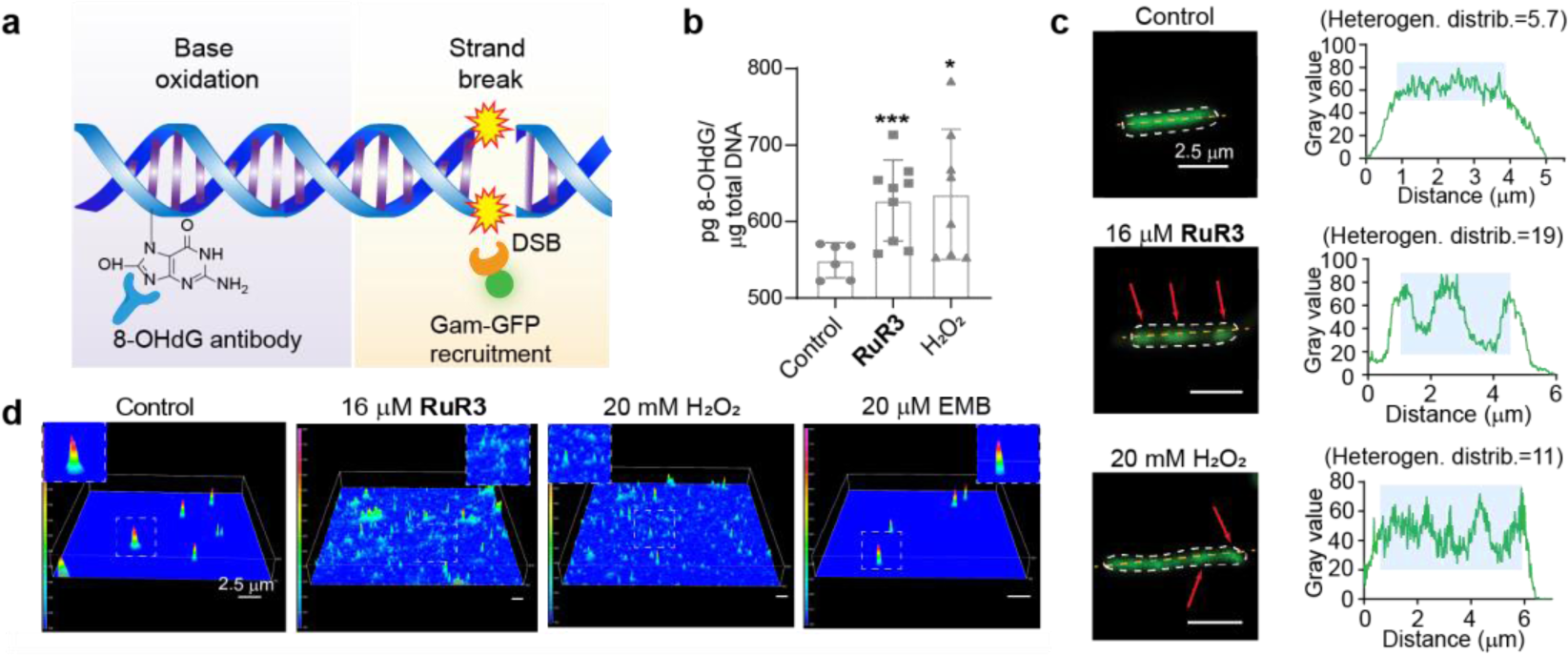
Oxidation of genomic DNA. (a) Scheme illustration of two major forms of DNA oxidation lesion: 8-OHdG formation and double strand break. (b) ELISA-based quantification of 8-OHdG level on genomic DNA of *M. smegmatis* upon treatment of **RuR3** and H2O2. (c) Results of Gam-GFP detection of DNA double strand break. Left: representative confocal images with arrows indicating Gam-GFP foci, which occur at double-strand breaks. Right: quantification of Gam-GFP distribution within *M. smegmatis*. Gam-GFP fluorescence heterogeneity is calculated as standard deviation of Gam-GFP distribution within the bacterial cell. (Heterogen. distrib.: heterogeneity distribution). (d) Results of single-cell gel electrophoresis assay to examine integrity of genomic DNA. Representative confocal images display intact genomic DNA in control and ethambutol treated samples, and DNA fragments in **RuR3** and H2O2 treated samples.

The ROS generation in bacterial cells was so prominent that we even observed the presence of double strand breaks (DSBs) in the bacterial chromosomes. The DSBs were examined by two methods: visualization using a previously reported engineered fluorescent protein-based probe (Gam-GFP)^80^., and quantification with an adapted comet assay via single-cell gel electrophoresis. The Gam-GFP probe relies on a fusion of GFP to the Gam protein from phage Mu that robustly binds to DSBs when expressed in bacteria. Without **RuR3**, Gam-GFP fusion proteins diffuse freely within bacterial cytosol (Figure 4c and Supplementary Fig. 9a, control), but quickly formed apparent foci when **RuR3** was added, suggesting prominent DSBs induced by the **RuR3**-causing endogenous ROS surge (Figure 4c and Supplementary Fig. 9a, 16 μM **RuR3**). The foci formation in the **RuR3**-treating group was even more prominent than in the positive control group, which was treated directly with H_2_O_2_ (Figure 4c and Supplementary Fig. 9a, 20 mM H_2_O_2_). Using the heterogeneity of GFP distribution as an indicator for the levels of DSBs in a single bacteria cell, statistical analysis again clearly revealed significantly higher DSB prevalence in the bacterial population treated with **RuR3** than those treated with H_2_O_2_ (Supplementary Fig. 9b). Consistently, when the lysate of **RuR3**-treated bacteria was analyzed by single-cell electrophoresis (Extended Data Fig. 4a, b-i), the results (Figure 4d and Supplementary Fig. 10a) indicated intact chromosomal DNA in the untreated and EMB-treated samples, but shattered DNA fragments for **RuR3-** or H_2_O_2_-treated samples. Statistical analysis of genomic DNA dimension showed that **RuR3** treated samples had significantly smaller overall size due to severe oxidative fragmentation (Extended Data Fig. 4b-ii and Supplementary Fig. 10b), consistent with the results obtained with the ensemble-level electrophoresis (Extended Data Fig. 4a, c).

### Dual-mode DNA damage endows the resistance-resistant nature of RuR3

It is known that extensive DNA damage in the bacterial chromosome triggers the induction of many distantly located genes, such as SOS response (Figure 5a)^81^. Therefore, we attempted to use SOS response as an indicator to quantify the level of damage by **RuR3** treatment and to analyze its correlation with other phenotypic measurements of its antimicrobial efficacy. In the SOS response, strand breaks and single-strand gaps in the DNA trigger the overexpression and activation of RecA protease, which cleaves the LexA repressor, leading to the induction of SOS genes involving in error-prone translesion DNA synthesis, such as *uvrA* and *dinB*. Transcription levels of these genes upon **RuR3** treatment were thus examined with qRT-PCR, which revealed that the expression of *recA*, *uvrA* and *dinB* was all upregulated significantly (2-105 fold, Figure 5b). Such results suggested the induction of strong SOS response upon the dual binding-oxidation mode of DNA damage from **RuR3**. In addition, *dinB* upregulation resulting from **RuR3** treatment was the highest compared to that from other DNA binding stressors (Figure 5c), indicating **RuR3** exerted the highest level of DNA damage to the bacterial chromosome. DinB overproduction is known to be cytotoxic to bacteria due to the production of 8-OHdG and double-strand DNA breaks^82^, corroborating with the observed oxidative damage of DNA from **RuR3** treatment (Figures 4b-d). Comparative analysis of the up- and down-regulated genes in whole transcriptomic profiling (Extended Data Fig. 5 and Supplementary Excel 2) revealed that over 43% of the differentially expressed genes were associated with DNA-related function and ROS stress response signaling pathway (Figure 5d).

**Figure 5.**
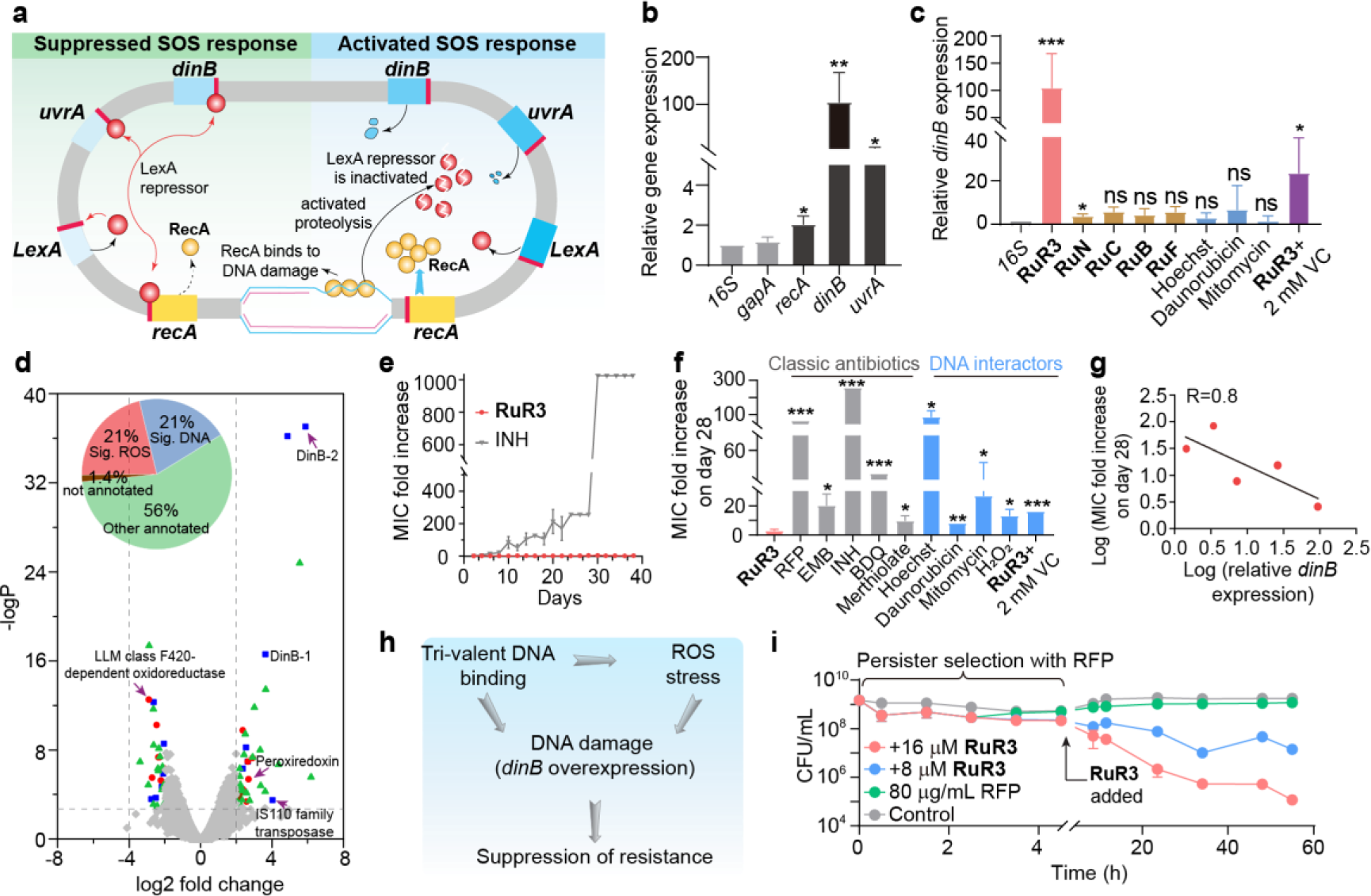
Dual-mode DNA damage endows the resistance-resistant nature and anti-persistent property of **RuR3**. (a) Schematic representation of SOS response to DNA damage in bacteria reproduced from reference. (b) The SOS response related genes were upregulated by treatment of 0.5 μM **RuR3**. (c) *dinB* upregulation in *M. smegmatis* under different treatments at 0.25×MIC. (d) Volcano plot of the transcriptome results using **RuR3**-treated *M. smegmatis*. log2 fold change: log2 (fold change of RPKM for a certain gene); −logP: −Log10 of P-value for the log2 fold change of a certain gene. (e) Resistance evolution profile of *M. smegmatis* against **RuR3** and isoniazid. Isoniazid exhibited a significant increase in MIC by ∼1000 fold after 274 generation. (f) Comparison of resistance generation of *M. smegmatis* under different drug treatments, revealing that **RuR3** has lowest level of resistance compared to conventional antimycobacterial (ethambutol, isoniazid and rifampicin), new generation of antimycobacterial approved in 2012 (bedaquiline), clinical metal-based antimicrobial (the Hg-based merthiolate, which are thought to have low resistance). (g) Correlation between resistance generation and level of DNA damage. (h) Schematic representation of generation of dual-mode DNA damage and the resulting resistance-resistent properties of **RuR3** treatment. (i) Killing of *M. smegmatis* persisters in 7H9 medium (plus 10% ADC) by **RuR3**. Persisters were generated *in vitro* with a pre-treatment of rifampicin at 10×MIC (80 μg/mL).

We next sought to see if the “catastrophic” DNA damage of **RuR3** treatment will lead to suppression of resistance emergence of mycobacteria. Resistance profiles of **RuR3** and related compounds were obtained by stepwise evolution of *M. smegmatis* under sub-inhibitory concentration of drug treatment for 40 days. **RuR3** treated sample was free from resistance development, while classic antibiotics exhibited significant increase in MIC (isoniazid in Figure 5e, and quantitative analysis of other antibiotics in Figure 5f and Extended Data Fig. 6a). In addition, **RuR3** remains highly active against the evolved antibiotic-resistant mutant of *M. smegmatis* (Extended Data Fig. 6b), as well as clinical multi-drugs resistant (cMDR) strains of *M. tuberculosis* (Extended Data Fig. 6c).

**Figure 6.**
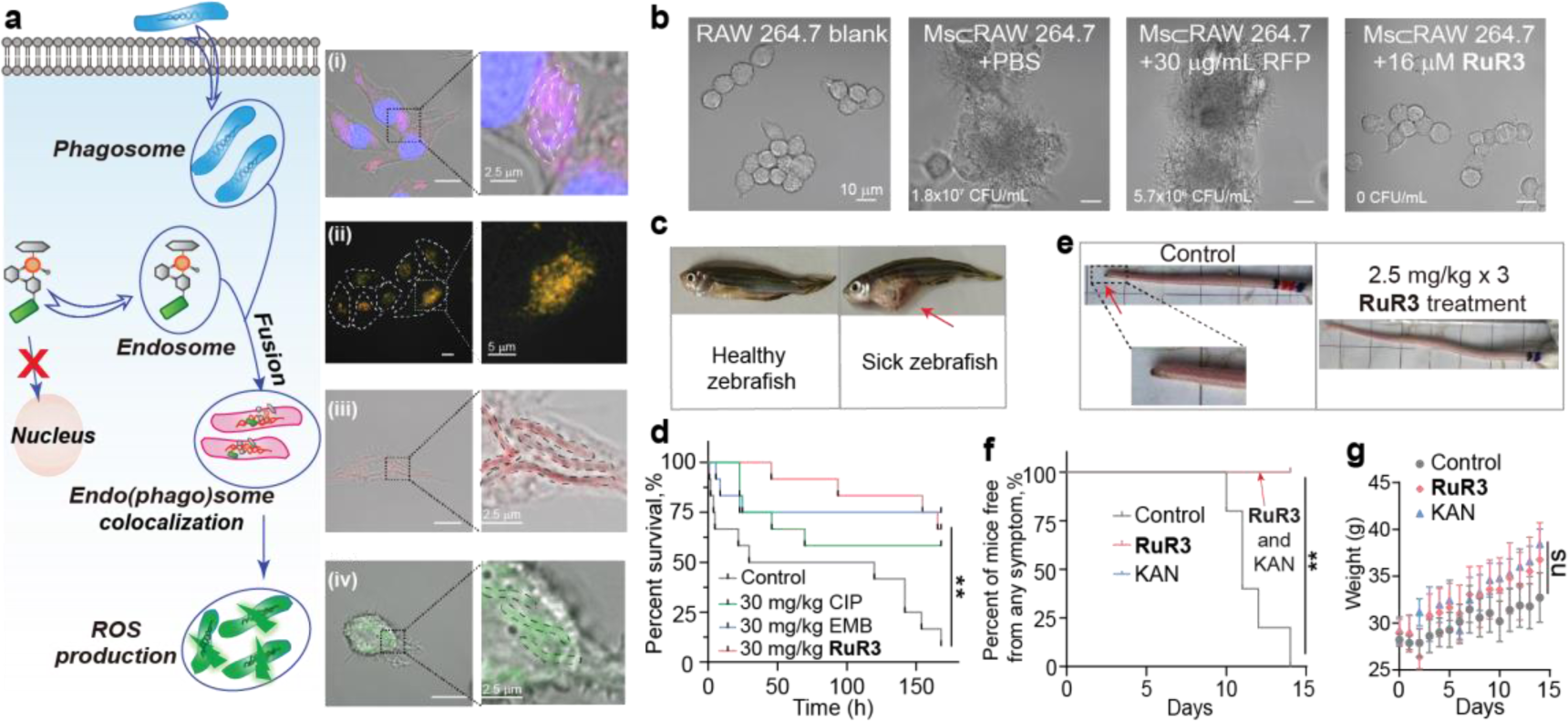
**RuR3**-induced bacterial specific ROS surge in *M. smegmatis* infected macrophages, and *in vivo* efficacy evaluation of **RuR3**. (a) Illustration of the intracellular fate of **RuR3** and phagocytosed mycobacteria. And confocal microscopic images of in *M. smegmatis* infected macrophages. (i) subcellular localization of *M. smegmatis* suggesting the residence of *M. smegmatis* in the acidic mycobacteria pathogen vacuoles (MCV, pH∼6.2) (blue: Hoechst-stained cell nucleus and *M. smegmatis*; red: lysotracker-red to stain acidic compart.); (ii) subcellular localization of **RuR3** suggesting the residence of **RuR3** in the acidic endosomes or lysosomes (red: **RuR3** fluorescence; green: lysotracker-green to stain acidic compartent); (iii) subcellular colocalization of **RuR3** and *M. smegmatis* (red: **RuR3** fluorescence); (iv) subcellular colocalization of **RuR3**-induced ROS and *M. smegmatis* (green: DCFH-DA probed ROS). (b) Brightfield images of Ms⊂RAW 264.7 cells after a 24 h treatment with 16 μM **RuR3**, 30 μg/mL RFP or PBS. (c) Healthy zebrafish or dead zebrafish with characteristic distended abdomen from *M. fortuitum* infection. (d) Increased survival rate for zebrafish treated with **RuR3** in the aquatic *M. fortuitum* infection model study. (e) Photos of mice tails taken 14 days post infection. Condition of tails in the **RuR3** treatment group was visually better than the ones in the control group. (f) Decreased symptom occurrence for mice treated with **RuR3** in the *M. marinum* infection model study. (g) Toxicity evaluation of **RuR3** during treatment infected mice. Weights of mice remained stable growth after three doses of **RuR3** injection.

To find further evidence for the causative relationship between extensive DNA damage and resistance suppression, we also tested the resistance profile of the dual mode DNA bomb **RuR3**, and compared it with other DNA modulators with various modes. Resistance evolution of *M. smegmatis* clearly show that, single-valent DNA binders (Figure 5f, Hoechst, Daunorubicin and Mitomycin) and weak ROS generation agents (Figure 5f, H_2_O_2_) uniformly have higher resistance than **RuR3**. In addition, when the **RuR3**-induced ROS level was reduced by ROS scavenger (Vitamin C), resistance accelerated noticeably. The trend of resistance emergence was correlated with overall DNA damage as measured by *dinB* overexpression (Figure 5g), providing evidence for our proposed mechanism that dual action of tight DNA binding and oxidation is the key to suppress resistance emergence (Figure 5h).

### Eradication of intracellular Mycobacteria and RuR3’s bactericidal activity against dormant and persistent bacteria

As a typical intracellular pathogen, Mycobacteria enter host cells via phagocytosis, residing in mycobacteria containing vacuoles (MCVs) and adopting a growth-dormant persistence state. Such state makes them insensitive to traditional growth-inhibiting antibiotics that target the cellular biosynthetic machinery essential for bacterial growth^83–89^. Therefore, to successfully eradicate intracellular mycobacteria, **RuR3** must be bactericidal to dormant and persistent mycobacteria, and we used bacteria maintained in nutrient-depleted medium (such as 1×PBS) as models^90–92^ for the non-replicating “dormant” intracellular mycobacteria for an evaluation. As the resistance of dormant mycobacteria is resulted from the shutdown of central metabolism and termination of replication for better survival in the host^90^, rifampicin, with a growth-inhibiting bacteriostatic mechanism of action at therapeutic concentration^4^, completely loses its activity even at high concentrations (Extended Data Fig. 7a, green). This is well expected since rifampicin’s antimicrobial target, RNA polymerase, is less abundant in such non-growth state^85^. An antimycobacterial currently in phase II clinical trial with combined enzyme and biogenesis targeting mechanism^93–95^, SQ109, displays only mediocre antimicrobial activity (blue). In sharp contrast, **RuR3** remained highly active with a minimum eradication concentration of 2 µM (Extended Data Fig. 7a, red), and a fast-bactericidal kinetics to completely sterilize the culture within 6 hours (Extended Data Fig. 7b-c and Supplementary Fig. 11). Furthermore, **RuR3** also exhibits high bactericidal ability against dormant *M. tuberculosis* (Extended Data Fig. 7d). As shown Figure 5i, persister cells were first generated in a (“persister selection period”, i.e., a 5-hour treatment with 80 μg/mL (10×MIC) of rifampicin)^90^, and without **RuR3** treatment, these bacteria are phenotypically resistant to the antibiotic (green trace, “80 μg/mL RFP”), although they are in fact genetically susceptible^88, 96^. In contrast, treatment of **RuR3** at 4× or 8×MIC concentration revealed high effectiveness against these persistent cells, both in nutrient-replete medium (Figure 5i, Blue and red traces) and nutrient-deplete medium (Extended Data Fig. 7e).

### Precise intracellular co-localization of RuR3 with MCVs

In addition to the bactericidal activity for persisters, another prerequisite for **RuR3**’s effectiveness against intracellular mycobacteria is its ability to precisely co-localize with intracellular bacteria. As we shall show in the following, **RuR3** is indeed able to achieve colocalization with intracellular bacteria through the adopted endocytic internalization pathway. After phagocytosis by the host cells, mycobacteria reside in phagosomal compartments called MCVs with a pH of 6.2 (Figure 6a-i, Extended Data Fig. 8a-b and Supplementary Fig. 12), which retains the ability to interact with early and recycling endosomes to acquire nutrients delivered by endosomal recycling pathways.^97, 98^ **RuR3** was observed to enter host cells through a micropinocytosis pathway Extended Data Fig. 8c and Supplementary Fig. 13), accumulating in the endosomal-lysosomal system (Figure 6a-ii, Extended Data Fig. 8d-e and Supplementary Fig. 14, 15) instead of cell nucleus or mitochondria (Extended Data Fig. 8f). Such behavioral feature facilitates **RuR3**’s fusion with the phagosomal MCVs and leads to specific targeting of intracellular mycobacteria, but not to any other compartments of the host cells (Figure 6a-iii and Supplementary Fig. 16). As **RuR3** shows no accumulation in the DNA-containing organelles, a.k.a nucleus and mitochondria, thus it has no interference with the DNA functions of the host cells. Meanwhile, the mycobacteria-specific localization of **RuR3** results in high localized ROS surge on the intracellular mycobacteria as well (Figure 6a-iv and Supplementary Fig. 17). As ROS generally have short life-times^99^ and travelling spans (<200 nm) ^100^, they are incapable of escaping the MCV and causing damage to the host cells once their generation sites are localized.

Intracellular bacteria clearance efficacy of **RuR3** was next evaluated (Supplementary Fig. 18a). With its unaltered antibacterial activity in the hydrolytic phagolysosome microenvironment (Extended Data Fig. 8g), **RuR3** effectively rescue the Ms⊂RAW 264.7 from mycobacteria invasion while leaving the host cell intact (Figures 6b, Extended Data Fig. 9a-b and Supplementary Fig. 18b). Thus, we have demonstrated in cellular level that, the dual-mode catastrophic DNA damaging mechanism of **RuR3** had resulted in resistance-resistant lethality specifically targeting intracellular mycobacteria. Such property was further examined in *in vivo* model systems as following.

### Efficacy evaluation using a zebra fish infection model and mouse models

Before proceeding to *in vivo* mycobacterial infection models, we first assessed the toxicity of **RuR3** on zebrafish and mice. For zebrafish, single dose treatment of 30 mg/kg **RuR3** was shown to be non-toxic (*N* = 5, Extended Data Fig. 9c). For mice model, two ways of drug administration were performed for toxicity evaluation. A preliminary test was carried out with a single dose of 100 mg/kg **RuR3** via intraperitoneal injection (*N* = 3), which showed no acute toxicity. In a second model, **RuR3** (2.5 mg/kg) was intravenously administrated for three times (on day 1, 4 and 7), and the mice was found to have maintained normal weight gain as the control group during treatment and in the follow-up observation period (Extended Data Fig. 9d). H&E staining of mice organs revealed no pathological change at day 17 after **RuR3** injection (Extended Data Fig. 9e). These results demonstrate **RuR3**’s high biocompatibility for *in vivo* applications.

To evaluate the potential of **RuR3** for *in vivo* therapeutic use, its *in vivo* performance was evaluated first in a zebrafish model. *M. fortuitum*, an aquaculture pathogen and also a causative agent for osteomyelitis, skin disease and joint infections in human^101^, was used to infect zebrafish at 1.5×10^8^ CFU. This infection resulted in the development of a characteristic distended abdomen of zebrafish in 1–3 days (Figure 6c and Supplementary Fig. 19), and eventual lethality in a few days if not treated (Figure 6d, “Control”). Intraperitoneal treatment with 30 mg/kg **RuR3** increased the survival rate of fish from 8.3% to 66.7%, outperforming ciprofloxacin and ethambutol (Figure 6d).

After this success in zebrafish, a mice mycobacterial infection model was then established with *M. marinum*, a marine counterpart of *M. tuberculosis* that causes chronic infection as an occupational disease for aquaculture and fishery workers^102^. Similar to *M. tuberculosis*, *M. marinum* is able to elicit a granulomatous disease, and is thus regarded as the best BSL2 model for tuberculosis^103–105^. With an optimal growth temperature of 32 °C, *M. marinum* causes a systemic tuberculosis-like infection and disease in ectotherms such as frogs and fish, as well as lesions in body parts with lower temperatures (e.g., skin and tail) for mammals. Thus, a mice tail infection model^106^ was established by injecting 2×10^7^ CFU of *M. marinum* intravenously to the tails of mice on day 0. Treatments with **RuR3** and kanamycin were administrated intravenously on day 1, 4 and 7. Severe symptoms, such as visible open lesions, blackened tails or broken-off tail-ends caused by tissue necrosis, were observed for mice in the PBS group starting from day 10 (Supplementary Fig. 20), consistent with literature reports, and the symptoms worsens each day. At day 14, visible symptoms were observed for all mice in the PBS group, while all mice in the **RuR3**-treated group were symptom-free (Figures 6e-f). In addition, the body weight of **RuR3** treatment group increased normally as the PBS treatment group, indicating the non-toxic nature of **RuR3** during mycobacterial infection treatment (Figure 6g).

## Discussion and Conclusion

The key findings we have in this report is the new mechanism to achieve resistance-resistant lethality toward mycobacteria by targeting bacterial DNA with a self-escalating damage mode. The anti-intuitive resistance-prone nature of DNA-targeting compounds was reversed by binding/oxidation DNA interaction, taking advantage of the endogenous ROS production mechanism. The multivalent nature of **RuR3**-DNA interaction forms that basis of downstream DNA function inhibition, as well as endogenous ROS stimulation. Molecular dynamic simulation reveals that **RuR3** can insert into DNA double helical structure via an unusual “puncturing” mode of binding as displayed in Extended Data Fig. 10a. In this **RuR3**-DNA covalent adduct, bipyridine moiety of Ru^II^-arene intercalates between two base pair planes and the Ru metal covalently binds to the G base at N7, while the Rhein tail extends into the minor groove of the opposite site. The ability of the Rhein group to cross through the base pair plane is presumably due to its additional ability to intercalate (Figure 3c), which offers initial binding with DNA. After the initial binding, since the groove binding ability of Rhein is stronger than its intercalating ability, also because of the tendency of bipyridine toward intercalation ability (Figure 3b-c), the Rhein group eventually “slides off” the intercalating site and binds into the groove binding site. Binding of Rhein offers a binding free energy calculated by the Molecular Mechanics/Generalized Born Surface Area (MM/GBSA) method of −33.4 kcal/mol between the unbonded **RuR3** and DNA, which is significantly more stable than that of the unbonded Ru^II^-arene and DNA (−25.8 kcal/mol, Extended Data Fig. 10b). This thus forms the molecular basis of proximity-enabled covalent binding between **RuR3** and DNA induced by the addition of Rhein group. Such “puncturing” mode of binding of **RuR3** has resulted in a higher degree of elongation of DNA helix (as quantified by helix rise per base pair) than the reported DNA intercalators and the non-covalent Ru^II^ arene-DNA complex (Extended Data Fig. 10c and Supplementary Fig. 21). This distorted DNA backbone leads to mis-alignment of important DNA-polymerase interactions (Extended Data Fig. 10d), resulting in a destabilization effect as large as 63 kcal/mol for the DNA-protein interaction, much more pronounced than that of the non-covalent Ru^II^ arene-DNA complex (Extended Data Fig. 10e). Therefore, the simulation provided strong structural evidence for the inhibition of *in vitro* DNA replication shown in Figure 3e. Thus overall, the multivalent nature as well as the unique **RuR3**-DNA interaction mode work together to endow high degree of DNA function perturbation, which leads to activation of the stress-responding endogenous ROS production.

The above endogenous mechanism of ROS generation of **RuR3** also endows the desired bacterial specificity of the destructive oxidative stress, displaying distinct selectivity patterns when compared with exogenous ROS generators such as H_2_O_2_ or ROSup. As shown in Extended Data Fig. 10f and Supplementary Fig. 22, the non-selective exogenous ROS generators exert oxidative damaging indiscriminately in all four conditions, namely, in PBS, in *M. smegmatis* cells, in host cells (RAW 264.7), and in *M. smegmatis* infected cells (Ms⊂RAW 264.7), leading to a high toxicity profile. In sharp comparison, **RuR3** only generates ROS in *M. smegmatis* existing conditions, since the origin of its ROS production was from endogenous bacterial metabolic alternation upon DNA binding. Therefore, the oxidative power of **RuR3** only selectively impacts in the bacteria-presenting environments (Ms and Ms⊂RAW 264.7), leaving the healthy host cells (RAW 264.7) unaffected and featuring a non-toxic profile (Figure 2f). With this attractive biocompatibility profile, together with its high antimicrobial activity against clinical *M. tuberculosis* multi-drug resistant strains, **RuR3** displays better therapeutic indexes compared with rifampicin against cMDR strains of *M. tuberculosis* (Extended Data Fig. 10g), representing a promising candidate for development of new generation of antimycobacterial. As this report provides the first detailed mechanistic investigation of DNA-targeting based resistance-resistant antimicrobial mechanism, and the excellent performance it has unveiled, we believe that the work can contribute to the development of next-generation resistance-resistant antibiotics, with the working mechanism readily applicable to other molecular scaffold designs.

## Acknowledgments

The authors thank Dr. Jian Sun (South China Agricultural University) for providing the library of the traditional Chinese medicine derived natural products, Dr. Jing Huang (Hunan University) for providing Pcold-ctxm-15, Dr. Yugang Bai (Hunan University), Dr. Liangdong Lyu (Fudan University) and Dr. Guodong Rao (University of California, Davis) for their helpful discussion.

## Funding

The funding support from the National Natural Science Foundation of China (Grants 22177031 to X.F., Grants 21977052, 22277056 to Z. S., Grants 62072199, 32161133002 to S. H., Grants 22077066 to H. L.)

## Author contributions

J. S., Z. S. and X. F. designed research; J. S., M. W., Y. T., A. Y., Z. Z., S. B., M. L., J. X., X. L., Y. S., P. H., W. W., F. L., Y. C., Q. C., H. L., S. H., Z. S. and X. F. performed research; all authors contributed to data analysis; J. S., M. W., Y. T., Z. S. and X. F. wrote the paper.

## Competing interests

H. L. and Z. S. are inventors on a patent (CN108530343B) for the **RuR1-3** reported in this manuscript. The authors declare that they have no other competing interests.

## Extended Figures

**Extended Data Fig. 1.**
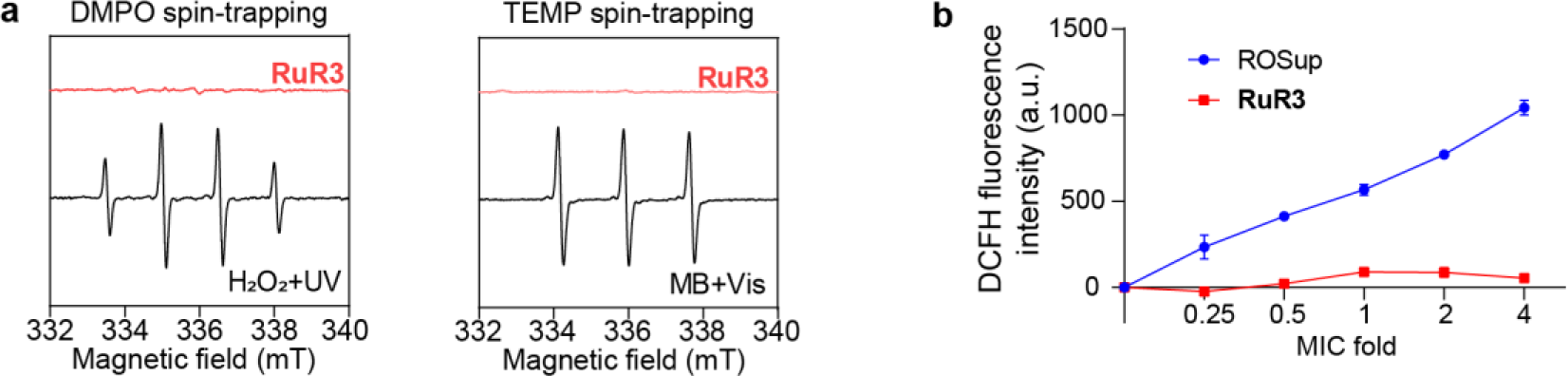
(a) DMPO spin-trapping EPR spectra of **RuR3** and H_2_O_2_ +UV in PBS solution. TEMP spin-trapping EPR spectra of **RuR3** and methylene blue (MB)+Vis in PBS solution. (b) DCFH fluorescence intensity (fluorescent probe allowing for detection of a vast range of ROS, λ_ex_ = 488 nm, λ_em_ = 525 nm) upon treatment with **RuR3** and ROSup at 0.25×/0.5×/1×/2×/4× MIC. MICs of **RuR3** and ROSup are 2 μM and 0.5 mg/mL, respectively.

**Extended Data Fig. 2.**
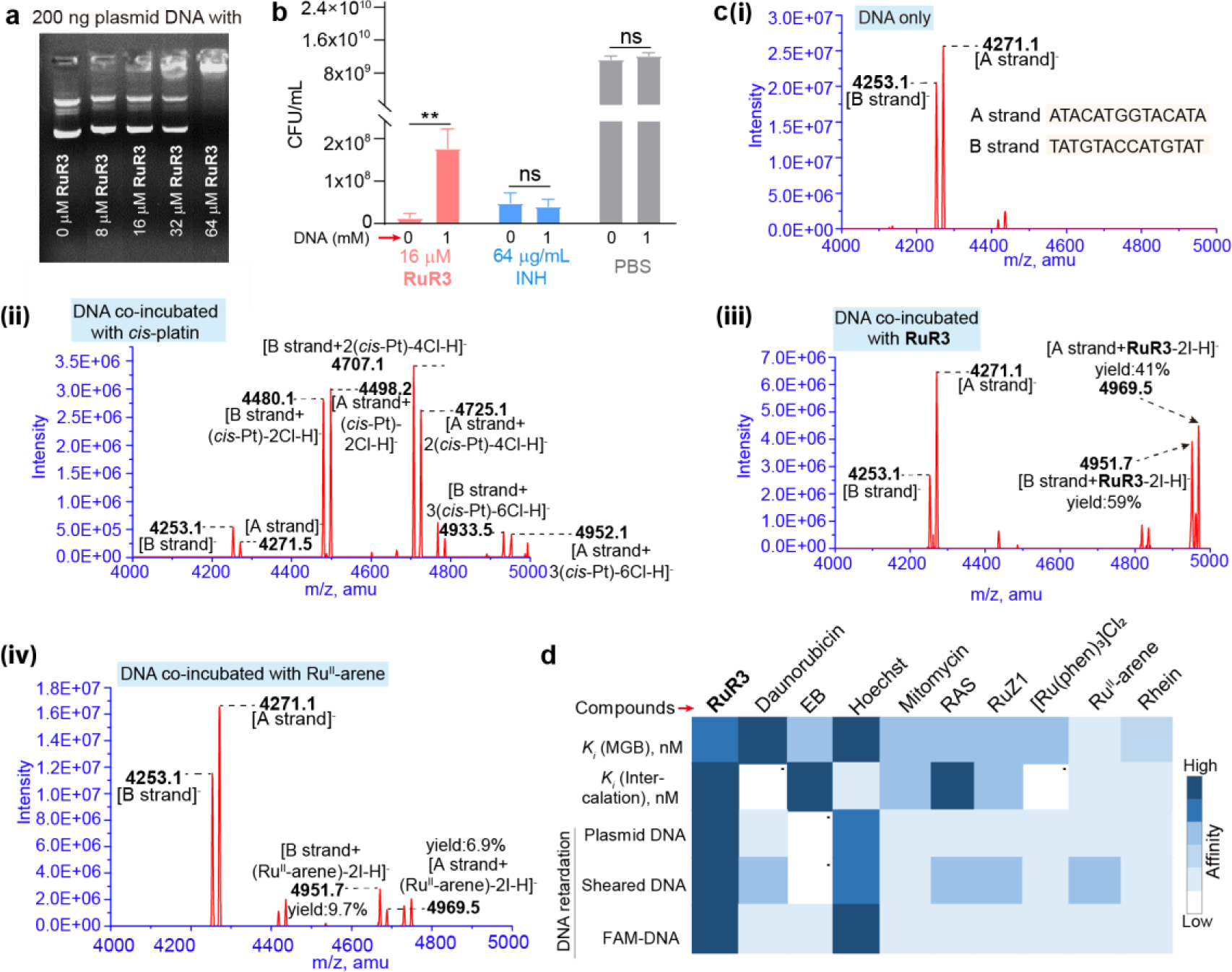
Additional evidence for multi-valent DNA binding by **RuR3**. (a) Electrophoretic mobility shift assay for visualization of **RuR3**-DNA interaction. Plasmid DNA (pCold-ctxm-15) were used in this assay. (b) Interference of 1 mM DNA on antimicrobial activities of **RuR3** and isoniazid. (c) Negative mode ESI-MS spectrum of d(ATACATGGTACATA)_2_: (i) untreated control; (ii) treated with *cis*-Pt, complexation of DNA and *cis*-Pt observed; (iii) treated with **RuR3**, complexation of DNA and **RuR3** observed; (iv) treated with Ru^II^-arene, complexation of DNA and Ru^II^-arene observed. (d) Heatmap of DNA binding affinity for **RuR3**, classic single mode DNA interactors, Ru-based metal complexes and two single components composing **RuR3**.

**Extended Data Fig. 3.**
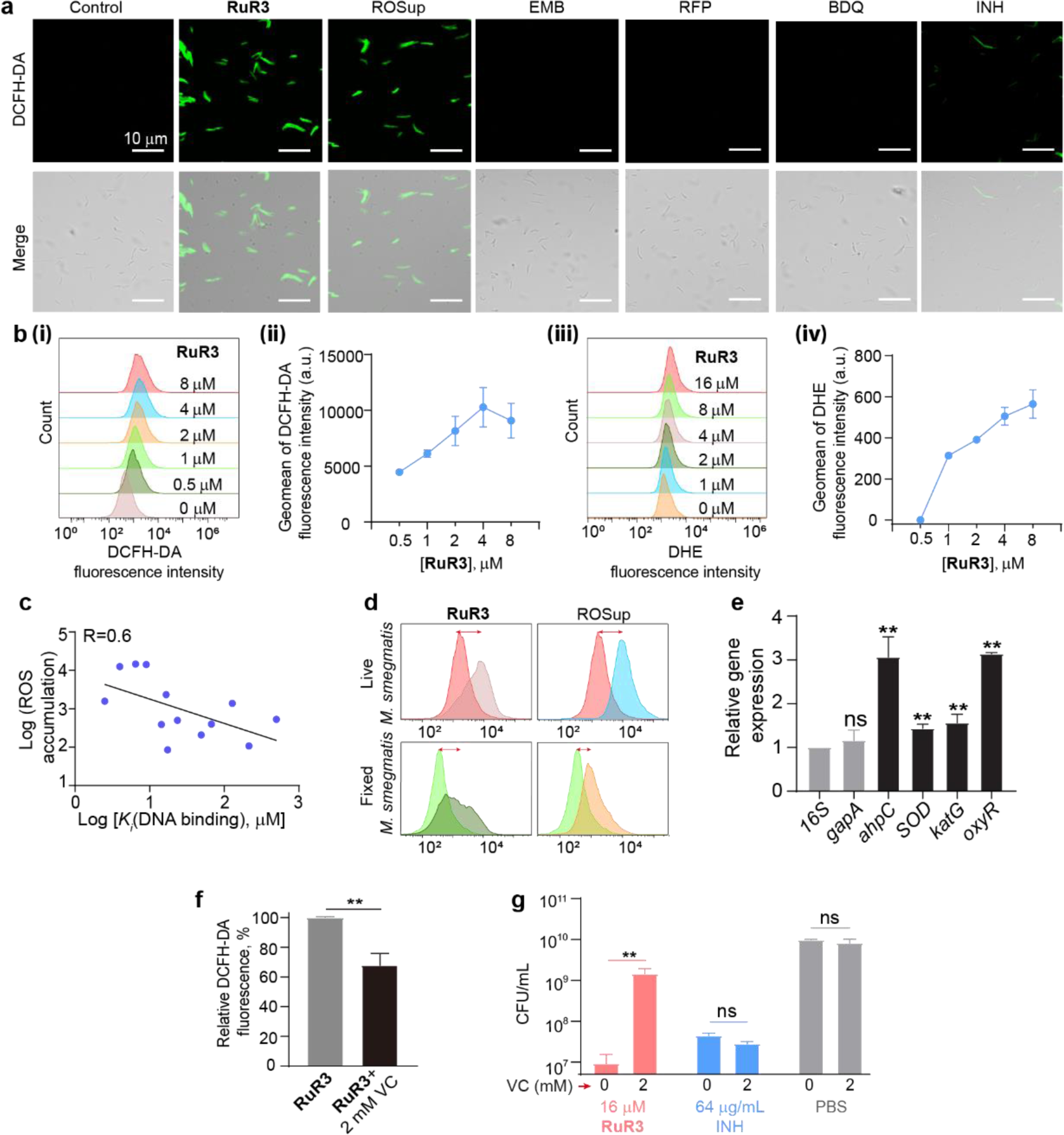
Endogenous ROS production of **RuR3**. (a) Confocal visualization of ROS accumulation in *M. smegmatis* upon treatment of **RuR3** and conventional stressors (EMB, RFP, BDQ and INH). ROS was probed by the green fluorescent dye DCFH-DA. (b) Flow cytometric analysis of ROS accumulation of **RuR3** under different concentrations, probed by DCFH-DA (fluorescent probe allowing for detection of a vast range of ROS, i-ii) or DHE (for O_2_^•-^, iii-iv). (c) The correlation between the ROS accumulation and DNA binding ability for compounds, fitted with a multiple linear regression model. (d) Flow cytometry quantification of **RuR3**- or ROSup-induced ROS accumulation in live *M. smegmatis* and fixed *M. smegmatis*. (e) Upregulation of ROS quenching enzyme in response to **RuR3** treatment at 0.5 μM. (f) Effect of Vitamin C (VC, a ROS scavenger) on reducing **RuR3** induced ROS accumulation. (g) Effect of Vitamin C on bactericidal activities of **RuR3** and isoniazid.

**Extended Data Fig. 4.**
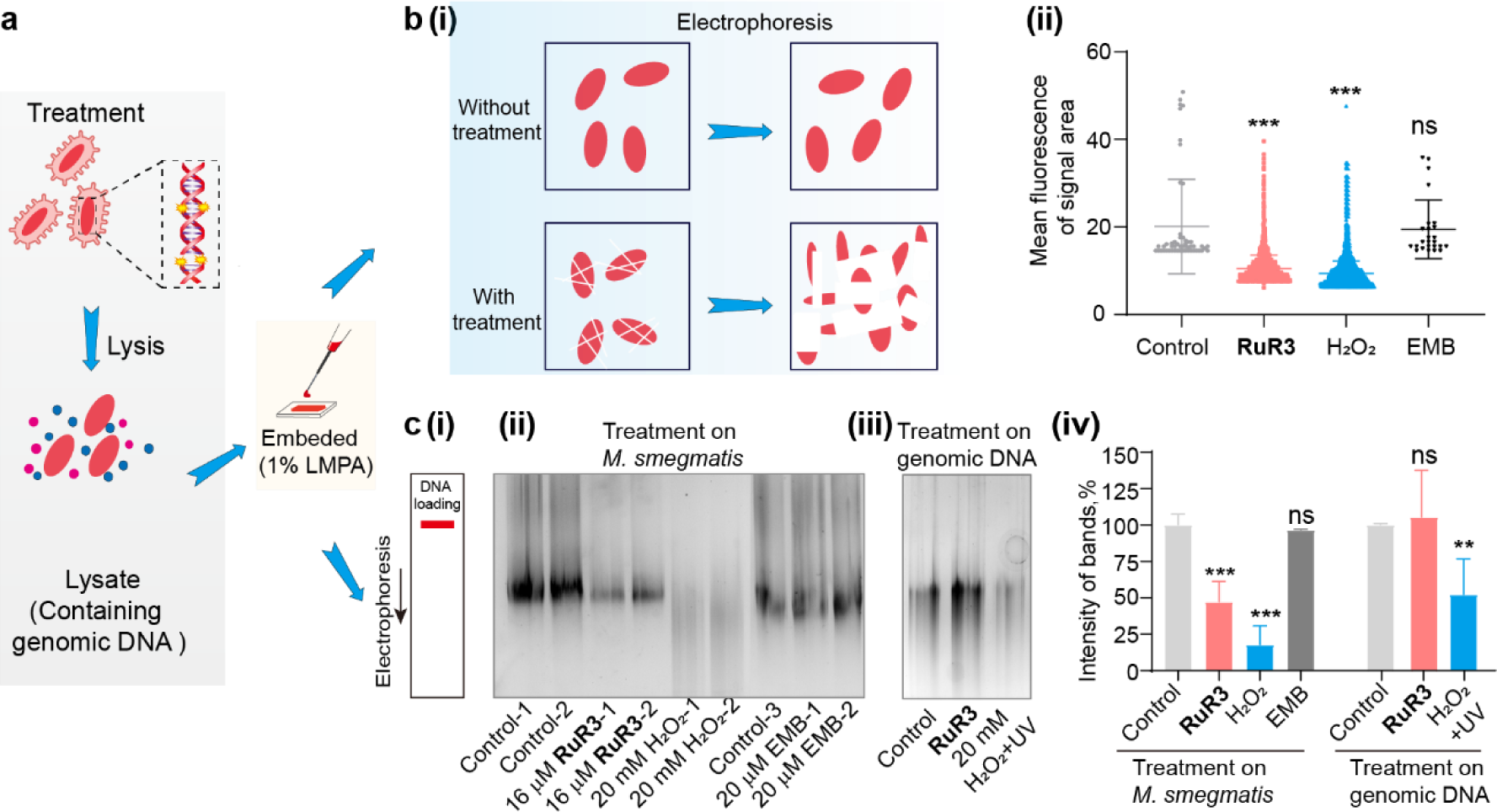
Additional evidence for oxidative fragmentation of genomic DNA. (a) Schematic illustration of sample preparation to evaluate genomic DNA integrity. (b) (i) Schematic illustration of single-cell gel electrophoresis assay. (ii) Statistical analysis of DNA fragmentation upon different treatments, calculating fluorescent intensities for pixels displaying fluorescent signals in Figure 4d. (c) (i) Schematic illustration of ensemble-level electrophoresis. (ii) Compound treatment on *M. smegmatis* cells, indicating oxidative fragmentation of DNA upon **RuR3** treatment. High concentration of H_2_O_2_ treatment serve as positive control of genomic DNA degradation. (iii) Compound treatment on free genomic DNA, indicating no fragmentation of DNA upon **RuR3** treatment. This is another evidence that **RuR3** resulting ROS is originated from bacterial endogenous metabolism. (iv) Quantitative analysis of c-ii and c-iii.

**Extended Data Fig. 5.**
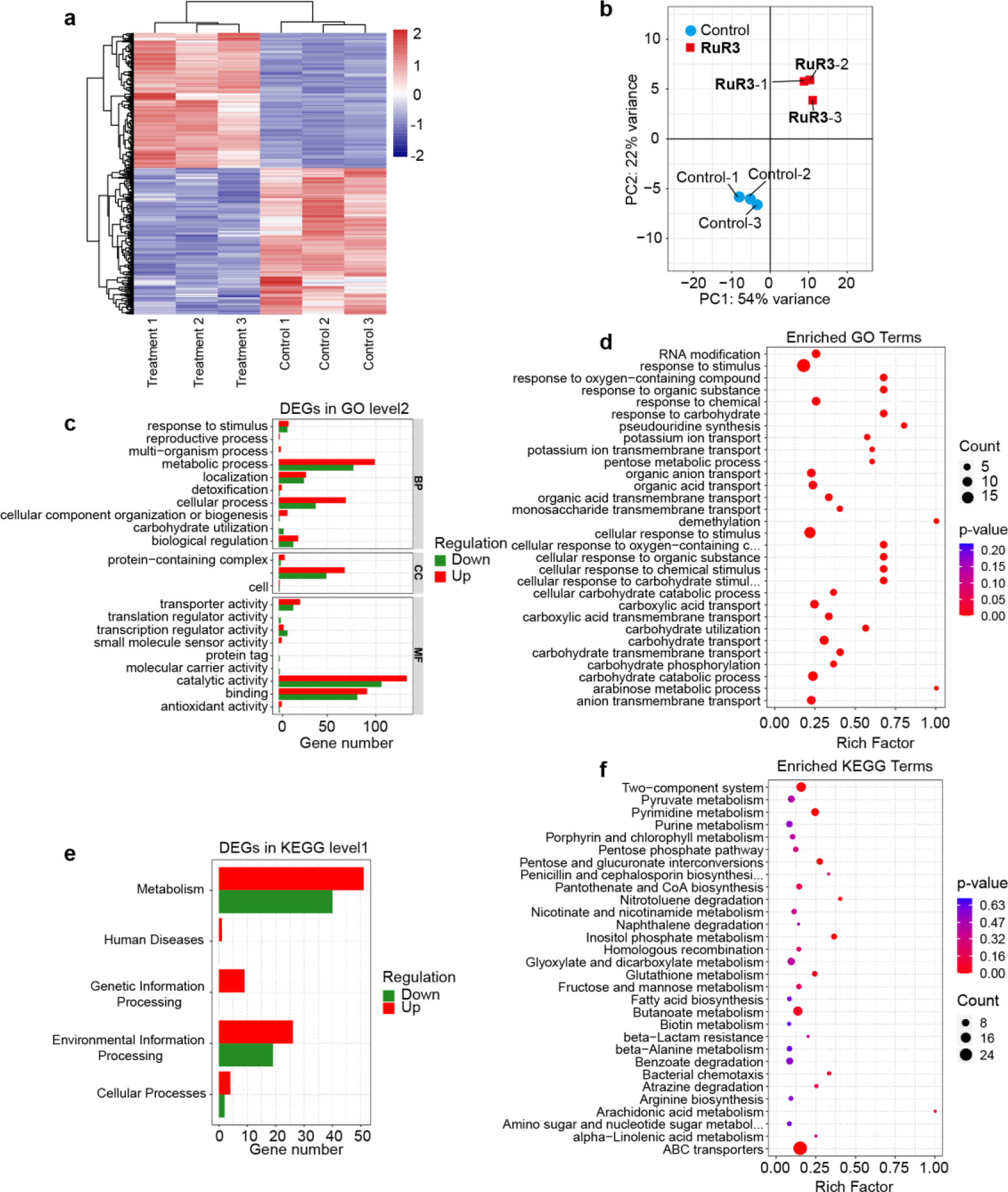
Transcriptomics analysis of **RuR3** treated *M. smegmatis*. (a) Heatmap of differential expression analysis showing gene regulation changes in *M. smegmatis* treated with 1 μM **RuR3**. (b) Principal component analysis (PCA) of *M. smegmatis* RNA-seq results. (c) GO level 1 and 2 analysis of differential expression genes (DEGs) of *M. smegmatis* in response to **RuR3** exposure. (d) Enrichment analysis of top 30 most significant GO terms in Biological process (BP) of *M. smegmatis* in response to **RuR3** exposure. The size of the dot indicates the number of DEGs involved in the pathway. The color scale indicates the significance level. The rich factor is the ratio between the number of DEGs and all genes enriched in the pathway. (e) KEGG level 1 pathway analysis of DEGs of *M. smegmatis* in response to **RuR3** exposure. (f) Enrichment analysis of top 30 most significant KEGG terms of *M. smegmatis* in response to **RuR3** exposure. The meanings of symbols are the same as in D. The fragment per kilobase per million mapped reads (FPKMs) of each gene was obtained by comparing **RuR3**-treated and untreated bacteria. The heatmap result indicates differential expression patterns between control and **RuR3** treatment groups (Extended Data Fig. 5a), supporting the hypothesis that the dual-mode DNA destructor **RuR3** affects genome-wide gene expression, which leads to misregulation of essential genes and subsequent cell death (Extended Data Fig. 5). Significantly Up- or down-regulated gene was defined with a cutoff value of |log_2_ (fold change) | ≥ 2 and P < 0.0015 (Supplementary Excel 2). Comparative analysis of the up- and down-regulated genes were carried out, which revealed that over 43% of them were associated with DNA-related function and ROS stress response signaling pathway including DinB family protein associated with DNA repair synthesis, IS110 family transposase (WP_011729446.1) with transposase activity via DNA binding, peroxiredoxin associated with ROS scavenge (WP_011730169.1), LLM class F420-dependent oxidoreductase (WP_011728963.1) (Figure 5d). Consistent with the qRT-PCR result showing *dinB* upregulation, a gene annotated as DinB family protein (WP_011727620.1), a poorly processive, error-prone DNA polymerase involved in untargeted mutagenesis, was most up-regulated in this transcriptome study by 59-fold.

**Extended Data Fig. 6.**
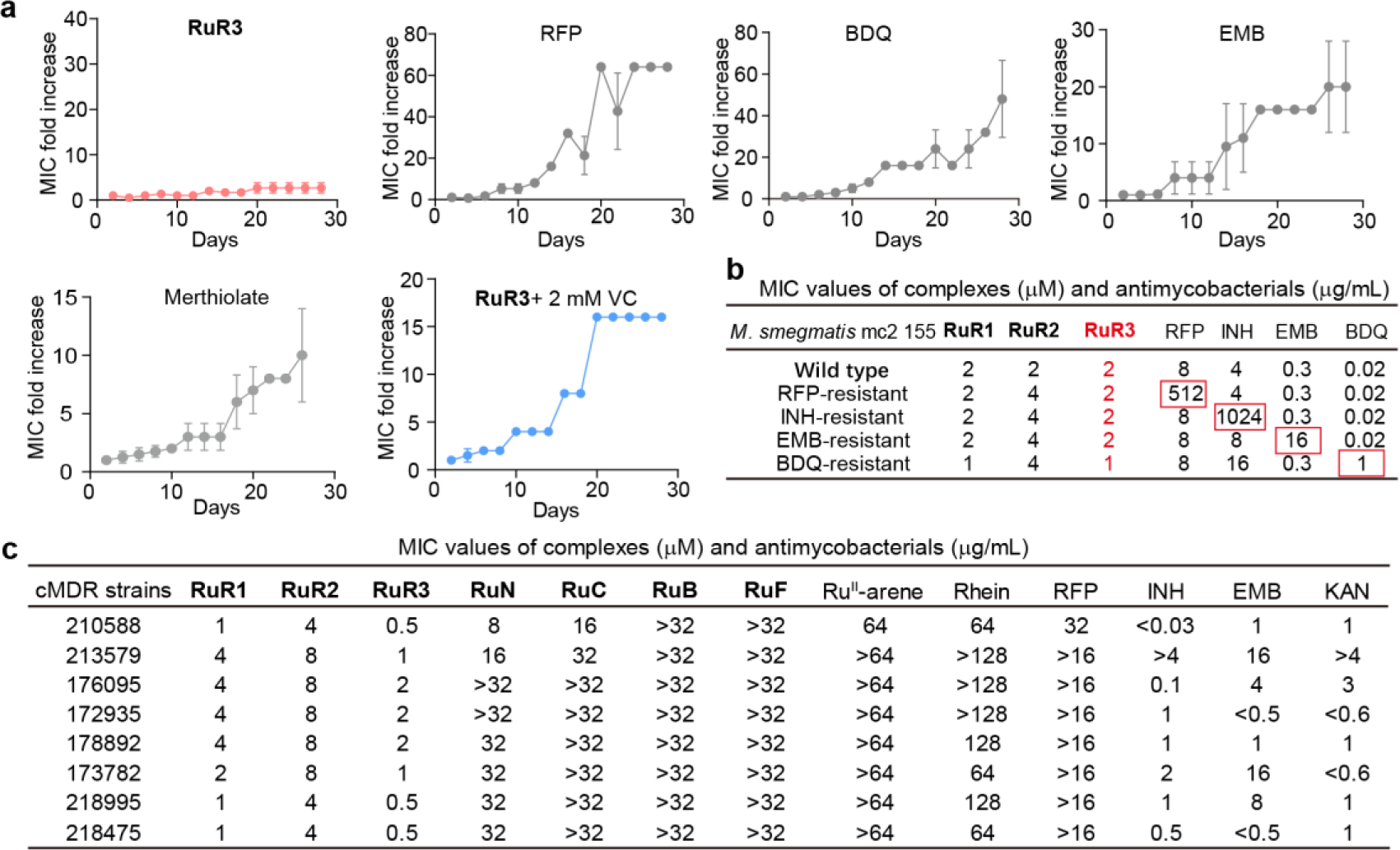
Resistance evolution of *M. smegmatis* under treatments of **RuR3** and other compounds. (a) Resistance generation profiles of **RuR3**, rifampin (RFP), bedaquiline (BDQ), ethambutol (EMB), merthiolate and **RuR3**+2 mM VC. (b) Cross-resistance examination of wild type *M. smegmatis* and mutants with various resistance, showing free of cross-resistance of **RuR**s. (c) MIC values of compounds against clinically isolated multi-drugs resistance (cMDR) *M. tuberculosis*.

**Extended Data Fig. 7.**
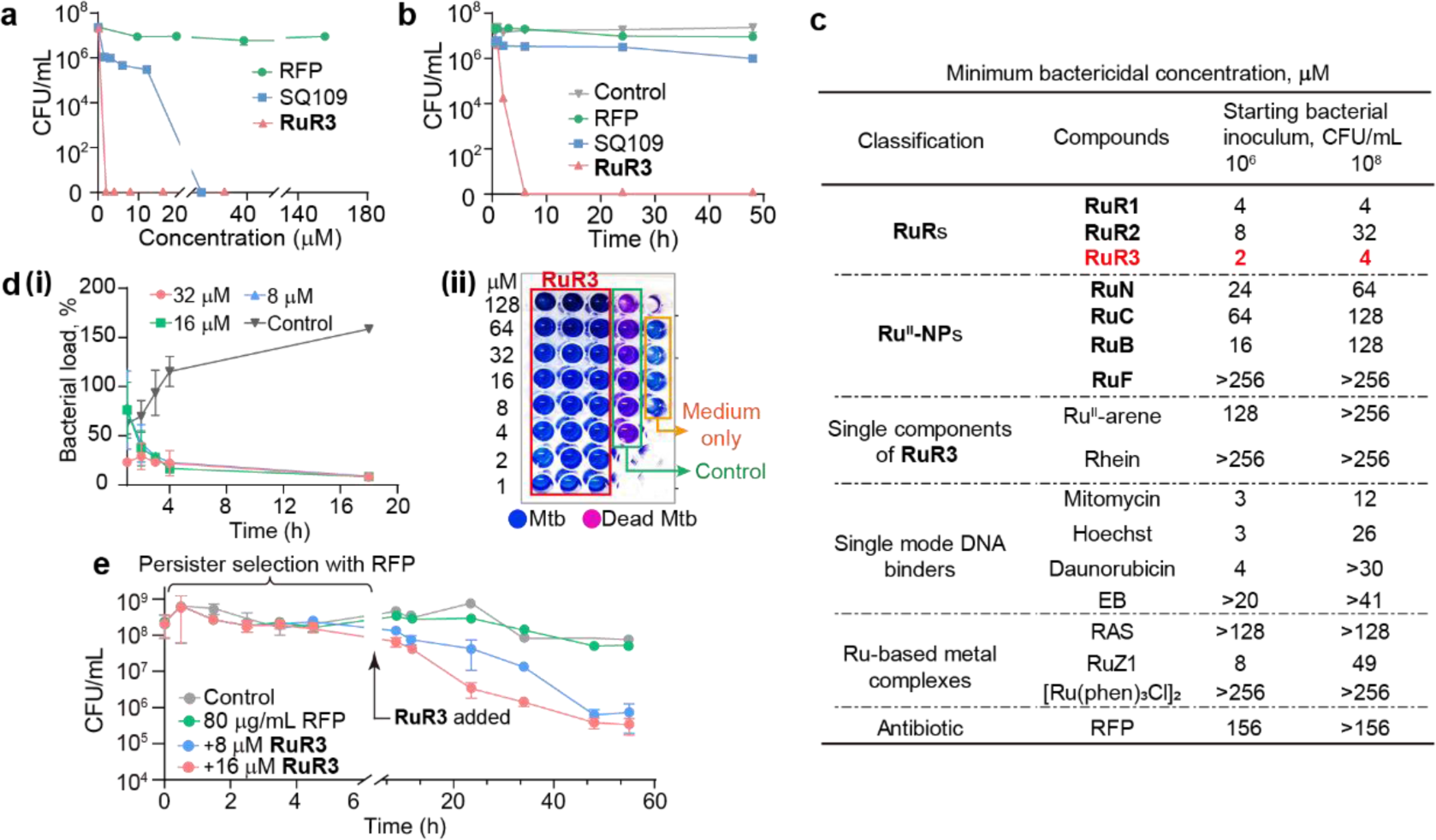
Antimicrobial activities against dormant and persistent *M. smegmatis*. (a) Dose-dependent antibacterial activities of **RuR3**, SQ109 and rifampicin (RFP) against dormant *M. smegmatis*. (b) Killing kinetics of **RuR3**, SQ109 and RFP against dormant *M. smegmatis* at 2×MIC. (c) MBC values of **RuR**s, other **Ru^II^-NP**s, single components of **RuR3**, single mode DNA binders, Ru-based metal complexes and RFP, suggesting best overall activity of **RuR3**. The initial cell density is ∼1×10^6^ or ∼1×10^8^ CFU/mL. (d) (i) Killing kinetics of dormant *M. tuberculosis* H37Rv in the presence of varying concentrations of **RuR3**. (ii) Visualization of **RuR3** killing of high-density *M. tuberculosis* H37Rv (initial OD_600_∼0.5), with resazurin as an indicator. (e) Killing of *M. smegmatis* persisters (induced with 80 μg/mL of RFP) by **RuR3** and RFP.

**Extended Data Fig. 8.**
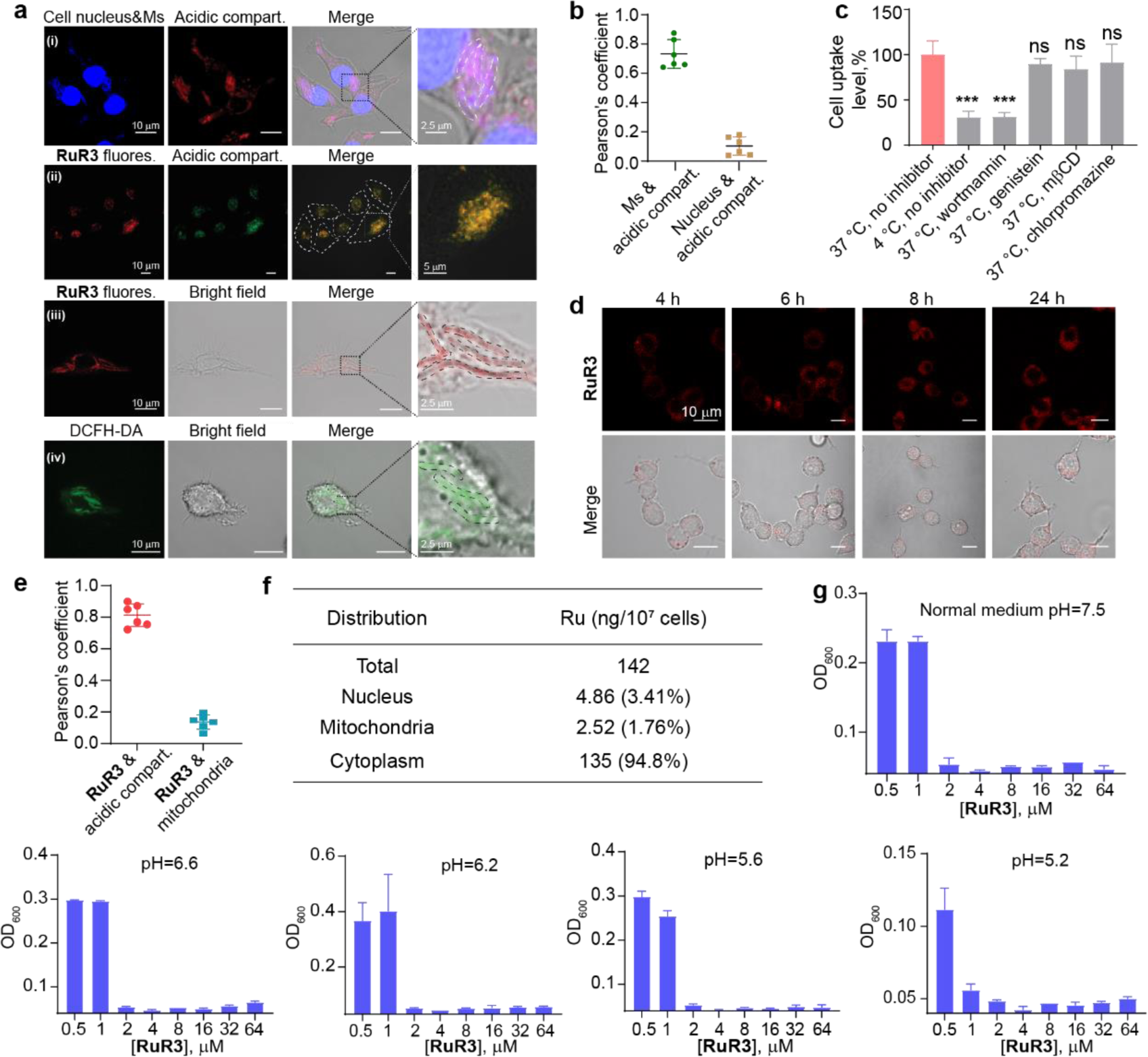
Intracellular localization of **RuR3** and **RuR3**-induced ROS. (a) Confocal microscopic images of in *M. smegmatis* infected macrophages. (i) subcellular localization of *M. smegmatis* suggesting the residence of *M. smegmatis* in the acidic mycobacteria pathogen vacuoles (MCV, pH∼6.2) (blue: Hoechst-stained cell nucleus and *M. smegmatis*; red: lysotracker-red to stain acidic compart.); (ii) subcellular localization of **RuR3** suggesting the residence of **RuR3** in the acidic endosomes or lysosomes (red: **RuR3** fluorescence; green: lysotracker-green to stain acidic compartent); (iii) subcellular colocalization of **RuR3** and *M. smegmatis* (red: **RuR3** fluorescence); (iv) subcellular colocalization of **RuR3**-induced ROS and *M. smegmatis* (green: DCFH-DA probed ROS). (b) Quantitative colocalization analysis of **RuR3**, acidic compartment (probed by Lysotracker) and cell nucleus within macrophage. (c) Mechanism of internalization of **RuR3**. As a standard method to investigate mode of entry, 4 °C condition, wortmannin, genistein, mβCD and chlorpromazine, respectively, inhibit energy-dependent endocytosis, macropinocytosis, clathrin-mediated endocytosis and caveolae-mediated endocytosis, in general. (d) Intracellular localization of **RuR3** (red fluorescence) at different time points, indicating its exclusion from the nucleus. (e) Quantification of **RuR3**/acidic compartment/mitochondria colocalization. (f) Intracellular distribution of Ru (ng/10^6^ cells) in RAW 264.7 after incubation with **RuR3** (10 μM) for 24 h at 37 °C. (g) Antimicrobial activities of **RuR3** under lower pH mimicking the hydrolytic phagolysosome microenvironment.

**Extended Data Fig. 9.**
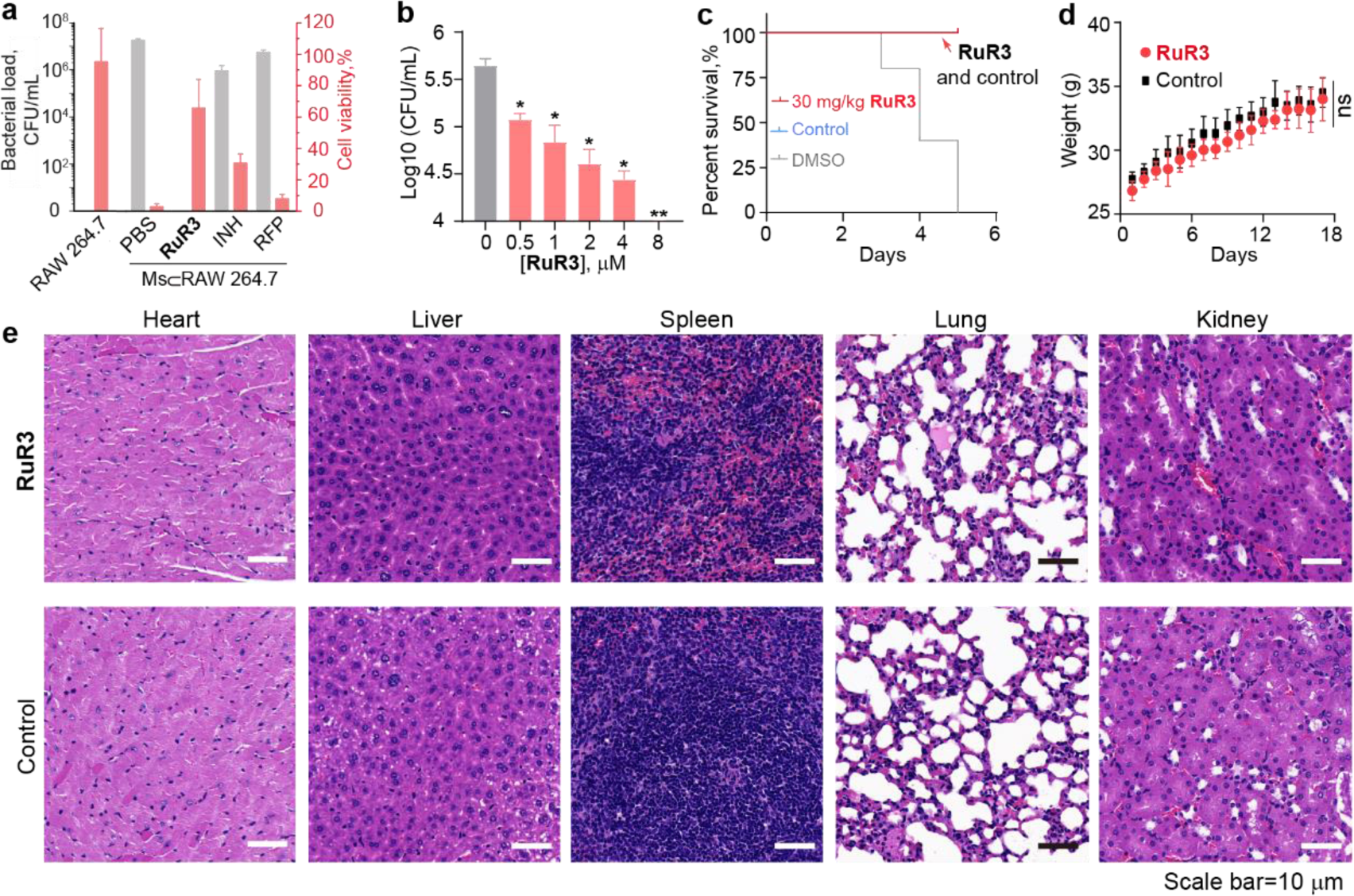
Eradication of intracellular mycobacteria and biocompatibility evaluation of **RuR3**. (a) Intracellular bacterial load (black columns) and mammalian cell viability (pink columns) of Ms⊂RAW 264.7 treated with 12 μM **RuR3**, 30 μg/mL RFP or 30 μg/mL INH. Uninfected RAW 264.7 was used as control. (b) Intracellular bacterial load of *M. fortuitous* infected RAW 264.7 treated with **RuR3** at different concentrations. (c) Toxicity evaluation of **RuR3** on healthy zebrafish. (d) Toxicity evaluation of **RuR3** on healthy mice. Weights of mice remained stable after three doses of **RuR3** intravenous (i.v.) injection (2.5 mg/kg). (e) Tissue immunohistology section images of mice treated with three i.v. injections of **RuR3** (2.5 mg/kg).

**Extended Data Fig. 10.**
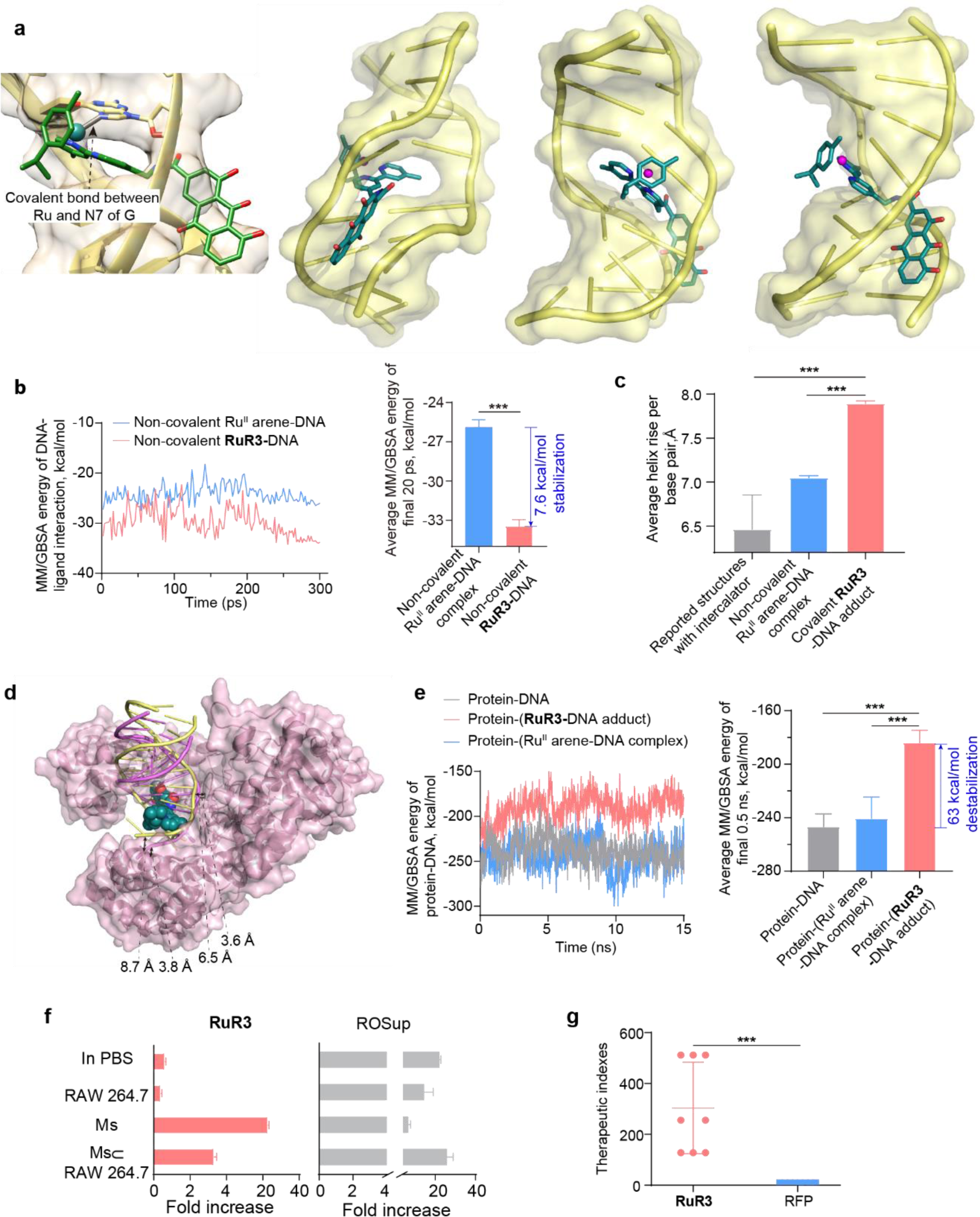
Molecular dynamics of multi-valent DNA binding by **RuR3** and its impact on DNA-protein interaction. (a) Molecular dynamic simulation of multi-valent **RuR3**-DNA binding, displaying an unusual “puncturing” mode of binding. (b) MM/GBSA energy trajectories for the DNA-ligand interaction during molecular dynamic simulation, and average MM/GBSA energy for the last 20 ps. (c) Comparison of average helix rise per base pair for reported structure with intercalators, non-covalent Ru^II^ arene-DNA complex and **RuR3**-DNA adduct. (d) Destabilization of DNA-polymerase interaction upon **RuR3** binding. Pick surface: *Bacillus* DNA Polymerase I (PDB ID code 1L3U); magenta DNA structure: original DNA bound to Polymerase I; yellow DNA structure: **RuR3**-liganded DNA; blue chemical structure: **RuR3**. (e) MM/GBSA energy trajectories for DNA-polymerase interaction during molecular dynamic simulation, and average MM/GBSA energy for the last 0.5 ns of molecular dynamic simulation. (f) Comparison of ROS level of **RuR3** and exogenous ROS generators in four different environments. (g) Therapeutic indices of **RuR3** and RFP for clinically isolated multi-drug resistant *M. tuberculosis* (cMDR *M. tuberculosis*). Therapeutic indexes are calculated as (IC_50_ for HEK293)/ (MIC for cMDR *M. tuberculosis*).

## Notes

### Competing Interest Statement

The authors have declared no competing interest.

