## Supporting Information for "Manipulation of Bacterial ROS Production Leads to Self-escalating DNA Damage and Resistance-resistant Lethality for Intracellular Mycobacteria"

### *Chemicals, bacterial strains and cell lines, instrumentations*

**Chemicals**

All solvents and reagents were purchased from Sigma-Aldrich, Aladdin, Selleck, TCI America, Adamas Reagent, Macklin Inc., J&K China, Beyotime Biotech., Sangon Biotech., TsingKe Biotech., and Vazyme Biotech., and were used without further purification unless otherwise specified.

4,4'-Dimethyl-2,2'-bipyridyl (L0), Rhein, Benzoic acid, Naproxen, Coumarin-3-carboxylic acid, Formic acid, EDCI (N-(3-Dimethylaminopropyl)-3-ethylcarbodiimide hydrochloride), HOBt (1-Hydroxybenzotriazole), $NH_4PF_6$ were obtained from HEOWNS; The metal arene dimer $[(\eta^6\text{-p-cym})RuCl_2]_2$, $[(\eta^6\text{-p-bip})RuCl_2]_2$, $[(\eta^6\text{-p-cym})RuI_2]_2$, DMSO (dimethyl sulfoxide) were purchased from Sigma Aldrich. Deuterated solvents for NMR purposes were obtained from Merck and Cambridge Isotopes. The library of the traditional Chinese medicine derived natural products was provided by School of Veterinary Medicine, South China Agricultural University.

**Biological reagents**

Calf thymus DNA (CT-DNA, NO. A002105) was purchased from Sangon Biotech. Sheared Herring Sperm DNA (sheared DNA, ~45 bp, D3159) was purchased from Sigma-Aldrich. Plasmid DNA pCold-ctxm-15 was a kind gift from Prof. Jing Huang.

Reactive Oxygen Species Assay Kit (S0033S) and Genomic DNA Mini-Extraction Kit (Universal, Spin Column, D0063) were purchased from Beyotime Biotech. 8-OHdG (8-hydroxydeoxyguanosine) ELISA Kit (E-EL-0028c) was purchased from Elabscience. HiPure Bacterial RNA Kit was purchased from Magen, Shanghai (R418102). HiScript II One Step qRT-PCR SYBR Green Kit was purchased from Vazyme (Q221-01). Gold mix (TSE101) was purchased from TsingKe Biotech.

Dulbecco's modified Eagle's medium (DMEM, SH30022.01) and fetal bovine serum (FBS, 13011-8611) were purchased from HYCLONE® and TIANHANG Biotech. Water was generated using a Milli-Q water purifier.

**Bacterial strains and cell lines**

*Mycobacterium smegmatis* (*M. smegmatis*) MC2 155 (ATCC 700084) was obtained from the American Type Culture Collection (ATCC, Manassas, VA). *Mycobacterium fortuitum* (*M. fortuitum*), *Mycobacterium marinum* (*M. marinum*), *Mycobacterium tuberculosis* (*M. tuberculosis*) H37Rv and clinically isolated multi-drugs resistance *M. tuberculosis* (cMDR) were provided by Hunan Institute for Tuberculosis Control. The bacterial strains were confirmed by 16S rDNA sequencing. Murine macrophage cell line RAW 264.7 (ATCC® TIB71™), Homo sapiens embryonic kidney cell line HEK293 (ATCC CRL-1573) were purchased from ATCC.

**Instrumentations**

The $^1H$ and $^{13}C$ NMR spectra were recorded on a Bruker AVANCE 400 spectrometer at ambient temperature. Electrospray ionization mass spectra (ESI-MS) were obtained using an LCQ spectrometer (Thermo Scientific). Cell imaging experiments were carried out on a confocal microscope (A1, Nikon, Japan). Fluorescence data were recorded on a fluorimeter (F-7000 spectrofluorometer, Hitachi). Flow cytometric quantification were carried out on flow cytometry (BD Accuri C6 Plus). Agarose gel and polyacrylamide gel imaging were recorded by using ChemiScope 6100 and Fusion FX spectra (Vilber) respectively. The Electron Paramagnetic Resonance (EPR) spectra were recorded on an X-band EPR spectrometer (JES-FA200, JEOL, Tokyo, Japan). qRT-PCR was performed by using QuantStudio 7 Flex (Thermo Fisher Scientific, USA). DNA annealing and *in vitro* DNA replication assay were carried out by

PCR instrument (Bio-rad, T100 Thermal Cycler).

**Special notes**

All experiments involving the use of *M. tuberculosis* must be performed in a laboratory that meets the standards of Biosafety Level 3 or above. All experiments involving the use of other pathogenic bacteria must performed in a laboratory that meets the standards of Biosafety Level 2 or above.

Animal Studies: All the animal studies were performed in accordance with the national and provincial regulations on animal studies. The certificate for the use of animals for research was SYXK (Hunan) 2022-0007, licensed by the Department of Science and Technology of Hunan Province on April 20, 2022, valid for 5 years. The protocols were approved by the Laboratory Animal Welfare and Ethics Review Committee, Hunan University.

### *Synthesis and characterization for Ru$^{II}$-complexes*

The synthetic schemes and supporting figures for characterizations of the compounds were given in Supplementary Figure 23-51.

**General procedures for the reaction between NP-COOH and Ligands (A, L1-L5):**

Ligand A, 4-Aminomethyl-4'-methyl-2,2-bipyridyl, was synthesized by following the reported procedures[1]. To a suspension of NP-COOH (2.5 mmol, 1 equiv), EDCI (2.5 mmol, 1 equiv), HOBt (2 mmol, 0.8 equiv) in DMF, Ligand A (2.5 mmol, 1 equiv) and 2 mL TEA were added in a 50 mL Schlenk tube. The reaction mixture was stirred at ambient temperature for 24 h, then treated with ice-cold water. The precipitate was filtered and re-dissolved in $CH_2Cl_2$, and then purified by column chromatography on silica gel eluted with $CH_2Cl_2/CH_3OH$ (v/v, 100: 1 ~ 40:1).

Synthesis of ligand L1: The solid was obtained as yellow powder. Yield: 0.578 g (50%). $^1$H NMR (400 MHz, $d_6$-DMSO) δ (ppm): 11.93 (*d*, *J* = 12.0 Hz, 2H), 9.68 (*t*, *J* = 5.9 Hz, 1H), 8.62 (*d*, *J* = 5.0 Hz, 1H), 8.51 (d, J = 5.0 Hz, 1H), 8.37 (s, 1H), 8.27-8.20 (m, 2H), 7.85 (dd, J = 10.1, 4.9 Hz, 2H), 7.77 (*d*, *J* = 7.4 Hz, 1H), 7.41 (*dd*, *J* = 10.4, 5.4 Hz, 2H), 7.28 (*d*, *J* = 4.3 Hz, 1H), 4.63 (*d*, *J* = 5.8 Hz, 2H), 2.41 (s, 3H).

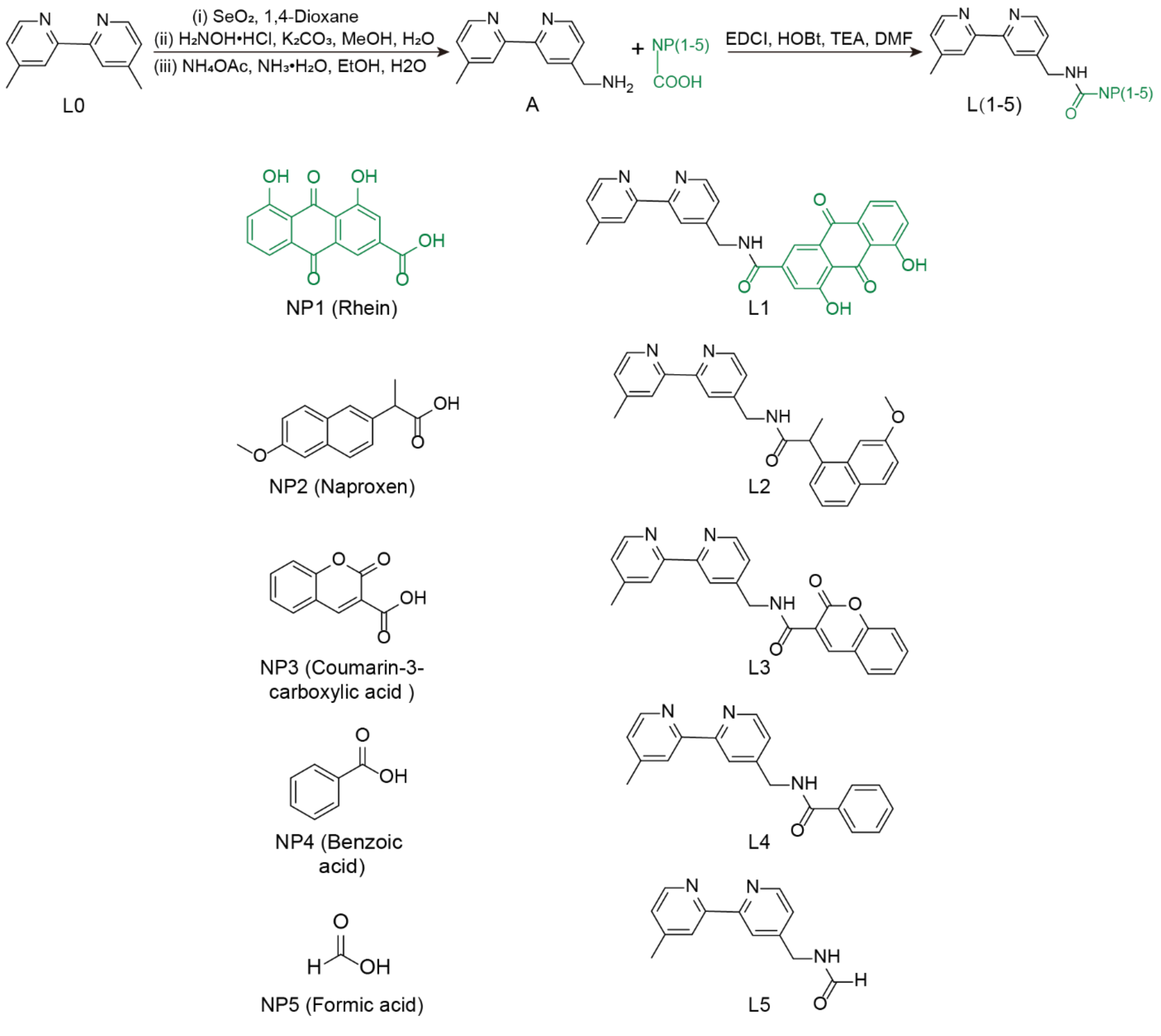


Synthesis of ligand L2: The solid was obtained as white powder. Yield: 0.495 g (48%). $^1$H NMR (400 MHz, $d_6$-DMSO) δ (ppm): 8.77 (*t*, *J* = 6.0 Hz, 1H), 8.53 (*dd*, *J* = 5.0, 0.7 Hz, 1H), 8.46 (*dd*, *J* = 4.8, 0.7 Hz, 1H),

8.32 (*s*, 1H), 8.21 (*s*, 1H), 7.77 (*s*, 1H), 7.77-7.74 (*m*, 2H), 7.51 (*dd*, *J* = 8.6, 1.7 Hz, 1H), 7.27 (*d*, *J* = 2.6 Hz, 1H), 7.24 (*dd*, *J* = 5.0, 1.8 Hz, 1H), 7.21 (*dd*, *J* = 5.0, 1.7 Hz, 1H), 7.13 (*dd*, *J* = 8.9, 2.5 Hz, 1H), 4.40 (*dd*, *J* = 5.9, 2.0 Hz, 2H), 3.90 (*t*, *J* = 7.0 Hz, 1H), 3.86 (*s*, 3H), 2.38 (*s*, 3H), 1.49 (*d*, *J* = 7.0 Hz, 3H).

Synthesis of ligand L3: The solid was obtained as white powder. Yield: 0.398 g (43%). $^1$H NMR (400 MHz, $d_6$-DMSO) δ (ppm): 9.35 (*t*, *J* = 6.2 Hz, 1H), 8.91 (*s*, 1H), 8.61 (*dd*, *J* = 5.0, 0.8 Hz, 1H), 8.52 (*d*, *J* = 4.9 Hz, 1H), 8.40-8.32 (*m*, 1H), 8.27-8.20 (*m*, 1H), 8.00 (*dd*, *J* = 7.9, 1.6 Hz, 1H), 7.77 (*ddd*, *J* = 8.7, 7.4, 1.6 Hz, 1H), 7.53 (*d*, *J* = 8.4 Hz, 1H), 7.45 (*td*, *J* = 7.6, 1.0 Hz, 1H), 7.42-7.36 (*m*, 1H), 7.28 (*ddd*, *J* = 4.9, 1.8, 0.9 Hz, 1H), 4.66 (*d*, *J* = 6.1 Hz, 2H), 2.41 (*s*, 3H).

Synthesis of ligand L4: The solid was obtained as white powder. Yield: 0.386 g (51%). $^1$H NMR (400 MHz, $d_4$-Methanol) δ (ppm): 8.61 (*d*, *J* = 5.1 Hz, 1H), 8.49 (*d*, *J* = 5.1 Hz, 1H), 8.27 (*s*, 1H), 8.14 (*s*, 1H), 7.93 (*d*, *J* = 7.5 Hz, 2H), 7.59 (*t*, *J* = 7.3 Hz, 1H), 7.51 (*t*, *J* = 7.5 Hz, 2H), 7.44 (*d*, *J* = 5.0 Hz, 1H), 7.30 (*d*, *J* = 5.0 Hz, 1H), 4.72 (*s*, 2H), 2.48 (*s*, 3H).

Synthesis of ligand L5: The solid was obtained as white powder. Yield: 0.312 g (55%). $^1$H NMR (400 MHz, *d*-Chloroform) δ (ppm): 8.64 (*d*, *J* = 5.0 Hz, 1H), 8.54 (*d*, *J* = 4.7 Hz, 1H), 8.38 (*s*, 1H), 8.31 (*s*, 1H), 8.24 (*s*, 1H), 7.25 (*d*, *J* = 3.1 Hz, 1H), 7.17 (*d*, *J* = 5.9 Hz, 1H), 6.20 (*s*, 1H), 4.61 (*d*, *J* = 6.2 Hz, 2H), 2.46 (*s*, 3H).

**General procedures for synthesis of Ru$^{II}$-complexes:**

A suspension of Ru$^{II}$-arene dimer (0.05 mmol, 1 equiv) and ligand (0.1 mmol, 2 equiv) in a $CH_3OH$ and $CH_2Cl_2$ mixture was refluxed overnight under argon atmosphere. The solution was cooled to room temperature and then was evaporated to dryness under reduced pressure.

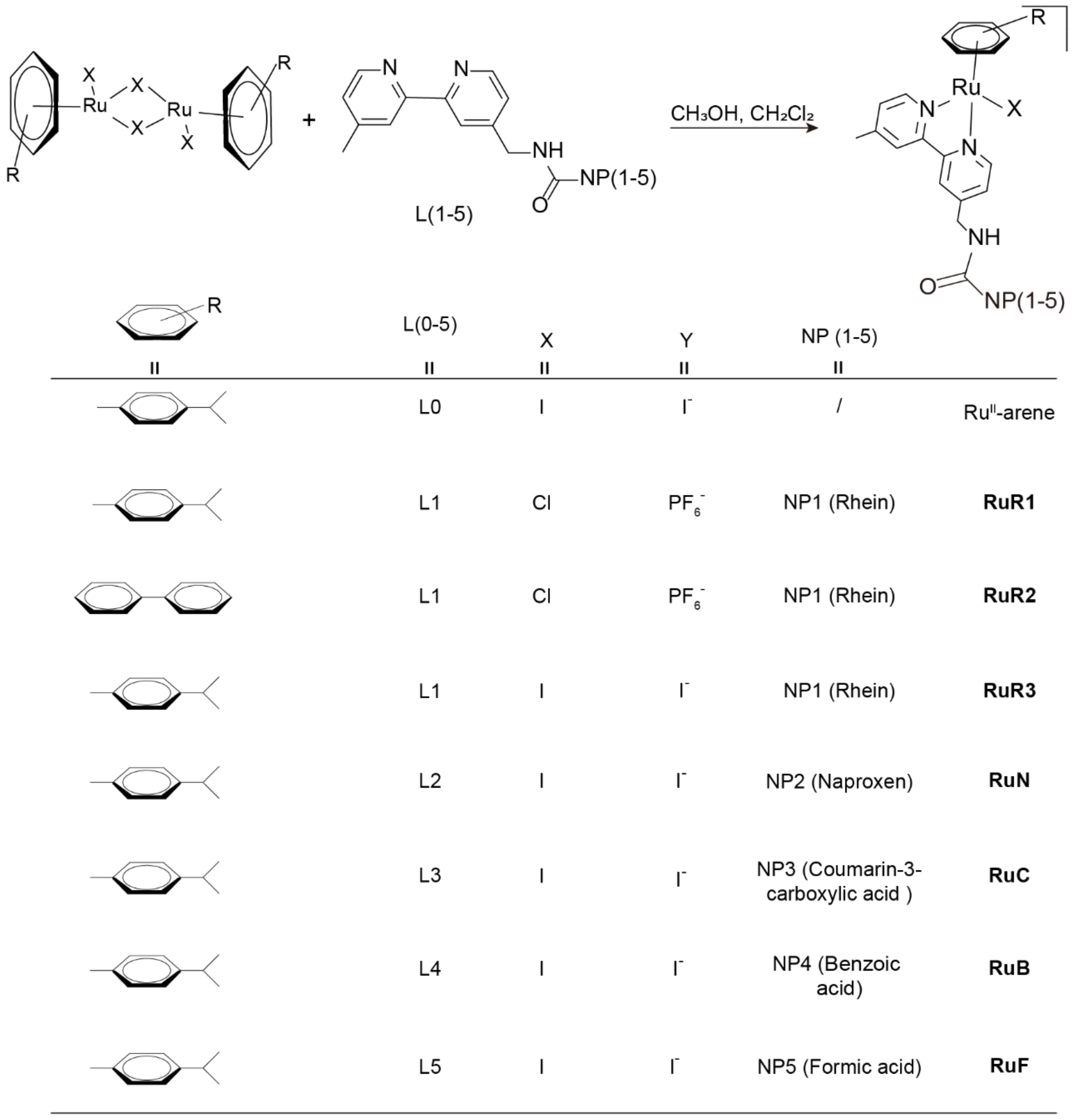


For the synthesis of **RuR1** and **RuR2**, the crude product was isolated by anion exchange with $NH_4PF_6$. The pure product of the complexes was obtained by recrystallizing from $CH_2Cl_2/Et_2O$.

For complex $Ru^{II}$-arene, containing no Rhein, was synthesized as a reference and the ligand used was 4,4'-Dimethyl-2,2'-bipyridyl (L0).

[($\eta^6$-p-cym)Ru(L0)I]I ($Ru^{II}$-arene). Complex $Ru^{II}$-arene was obtained as yellow powder. Yield: 43.0 mg (64%). $^1$H NMR (400 MHz, $d_6$-DMSO) δ (ppm): 9.29 (*d*, *J* = 5.8 Hz, 2H), 8.51 (*s*, 2H), 7.59 (*d*, *J* = 5.9 Hz, 2H), 6.11 (*d*, *J* = 6.1 Hz, 2H), 5.98 (*d*, *J* = 6.1 Hz, 2H), 2.67 (*s*, 1H), 2.57 (*s*, 6H), 2.40 (*s*, 3H), 0.96 (*d*, *J* = 6.8 Hz, 6H). $^{13}$C NMR (101 MHz, $d_6$-DMSO) δ (ppm): 156.43, 154.28, 152.24, 128.53, 124.73, 106.46, 101.86, 86.15, 85.34, 31.36, 22.00, 21.04, 20.49. ESI-MS ($CH_3OH$): calcd for $[Ru^{II}\text{-arene-}I^-]^+$ m/z = 546.43, found m/z = 547.02.

[($\eta^6$-p-cym)Ru(L1)Cl]$PF_6$ (**RuR1**). Complex **RuR1** was obtained as yellow crystalline powder. Yield: 62.0 mg (70%). $^1$H NMR (400 MHz, $d_6$-DMSO) δ (ppm): 9.75 (*t*, *J* = 5.8 Hz, 1H), 9.46 (*d*, *J* = 5.9 Hz, 1H), 9.38 (*d*, *J* = 5.8 Hz, 1H), 8.61 (*s*, 1H), 8.53 (*s*, 1H), 8.21 (*s*, 1H), 7.90 (*s*, 1H), 7.85 (*t*, *J* = 7.9 Hz, 1H), 7.77 (*d*, *J* = 7.4 Hz, 1H), 7.67 (*dd*, *J* = 14.3, 5.9 Hz, 2H), 7.44 (*d*, *J* = 8.3 Hz, 1H), 6.20 (*dd*, *J* = 9.7, 6.2 Hz, 2H), 5.96 (*t*, *J* = 7.0 Hz, 2H), 4.77 (*s*, 2H), 2.58 (*s*, 3H), 2.17 (*s*, 3H), 1.23 (*s*, 1H), 0.95 (*dd*, *J* = 6.9, 4.5 Hz, 6H). $^{13}$C

NMR (101 MHz, $d_6$-DMSO) δ (ppm): 191.92, 181.62, 165.18, 161.88, 161.63, 155.95, 155.46, 154.68, 154.26, 153.15, 152.49, 141.35, 138.10, 134.20, 133.78, 128.79, 125.89, 125.11, 124.83, 123.22, 122.48, 119.97, 118.40, 118.11, 116.64, 103.90, 103.67, 86.95, 86.80, 84.33, 84.12, 42.64, 30.86, 22.18, 22.12, 21.14, 18.77. ESI-MS ($CH_3OH$): calcd for [**RuR1**-$PF_6^-$]$^+$ m/z = 736.21, found m/z = 736.25.

[($\eta^6$-p-bip)Ru(L1)Cl]$PF_6$ (**RuR2**). Complex **RuR2** was obtained as olive yellow crystalline powder. Yield: 67.0 mg (74%). $^1$H NMR (400 MHz, $d_6$-DMSO) δ (ppm): 9.57 (*s*, 1H), 9.30 (*d*, *J* = 6.0 Hz, 1H), 9.20 (*d*, *J* = 5.9 Hz, 1H), 8.56 (*s*, 1H), 8.48 (*s*, 1H), 8.00 (*s*, 1H), 7.72 (*dd*, *J* = 13.9, 6.0 Hz, 2H), 7.68-7.60 (*m*, 3H), 7.55 (*dd*, *J* = 17.3, 5.9 Hz, 2H), 7.42 (*dt*, *J* = 9.0, 6.8 Hz, 3H), 7.31 (*d*, *J* = 8.3 Hz, 1H), 6.63 (*t*, *J* = 4.2 Hz, 2H), 6.44 (*q*, *J* = 6.2 Hz, 2H), 6.19 (*t*, *J* = 5.7 Hz, 1H), 4.69 (*s*, 2H), 2.53 (*s*, 3H). $^{13}$C NMR (101 MHz, $d_6$-DMSO) δ (ppm): 191.60, 181.64, 165.28, 161.90, 161.81, 155.65, 155.13, 154.61, 154.13, 153.20, 152.60, 141.11, 138.05, 134.13, 133.63, 133.35, 130.58, 129.38, 128.75, 128.62, 125.75, 125.19, 124.72, 123.32, 122.40, 119.93, 118.29, 117.90, 116.49, 100.93, 88.96, 88.89, 85.02, 83.99, 42.59, 21.05. ESI-MS ($CH_3OH$): calcd for [**RuR2**-$PF_6^-$]$^+$ m/z = 756.20, found m/z = 756.17.

[($\eta^6$-p-cym)Ru(L1)I]I (**RuR3**). Complex **RuR3** was obtained as brown crystalline powder. Yield: 75.0 mg (79%). $^1$H NMR (500 MHz, $d_6$-DMSO) δ (ppm): δ 11.95 (*s*, 2H), 9.68 (*t*, *J* = 5.7 Hz, 1H), 9.40 (*d*, *J* = 5.9 Hz, 1H), 9.32 (*d*, *J* = 5.8 Hz, 1H), 8.61 (*s*, 1H), 8.54 (*s*, 1H), 8.25 (*s*, 1H), 7.93-7.83 (*m*, 2H), 7.79 (*d*, *J* = 7.5 Hz, 1H), 7.64 (*dd*, *J* = 15.7, 5.9 Hz, 2H), 7.45 (*d*, *J* = 8.2 Hz, 1H), 6.13 (*t*, *J* = 5.6 Hz, 2H), 6.05-5.94 (*m*, 2H), 4.79 (*t*, *J* = 5.4 Hz, 2H), 2.72 (*p*, *J* = 7.0 Hz, 1H), 2.60 (*s*, 3H), 2.39 (*s*, 3H), 0.99 (*dd*, *J* = 12.7, 6.9 Hz, 6H). $^{13}$C NMR (101 MHz, $d_6$-DMSO) δ (ppm): 191.88, 181.65, 165.19, 161.89, 161.67, 157.17, 156.67, 154.56, 154.17, 152.91, 152.22, 141.31, 138.09, 134.20, 133.79, 128.68, 125.69, 125.05, 123.25, 122.57, 119.95, 118.42, 118.09, 106.74, 101.63, 86.19, 86.07, 85.87, 85.68, 42.53, 31.41, 22.15, 22.11, 21.11, 20.32. ESI-MS ($CH_3OH$): calcd for [**RuR3**-$I^-$]$^+$ m/z = 827.65, found m/z = 828.05.

[($\eta^6$-p-cym)Ru(L2)I]I (**Ru-Naproxen**, **RuN**). Complex **RuN** was obtained as yellow powder. Yield: 59.0 mg (66%). $^1$H NMR (400 MHz, $d_6$-DMSO) δ (ppm): 9.36-9.32 (*m*, 1H), 9.32-9.28 (*m*, 1H), 8.78 (*t*, *J* = 6.0 Hz, 1H), 8.38 (*d*, *J* = 13.5 Hz, 1H), 8.25 (*d*, *J* = 23.8 Hz, 1H), 7.81-7.72 (*m*, 3H), 7.60 (*d*, *J* = 6.0 Hz, 1H), 7.52-7.47 (*m*, 1H), 7.42 (*dd*, *J* = 12.7, 5.8 Hz, 1H), 7.29 (*d*, *J* = 2.5 Hz, 1H), 7.16-7.08 (*m*, 1H), 6.11 (*q*, *J* = 6.5 Hz, 2H), 6.04-5.92 (*m*, 2H), 4.52 (*q*, *J* = 6.6, 6.1 Hz, 2H), 3.91 (*d*, *J* = 6.6 Hz, 1H), 3.86 (*s*, 4H), 3.17 (*d*, *J* = 5.2 Hz, 3H), 2.70-2.62 (*m*, 1H), 2.37 (*d*, *J* = 1.6 Hz, 3H), 1.48 (*dd*, *J* = 7.0, 2.3 Hz, 3H), 0.99-0.89 (*m*, 6H). $^{13}$C NMR (101 MHz, $d_6$-DMSO) δ (ppm): 174.91, 157.47, 156.82, 156.48, 154.30, 153.92, 153.45, 152.19, 137.36, 133.60, 129.58, 128.77, 128.63, 127.32, 126.72, 125.79, 125.42, 124.44, 121.90, 119.11, 106.69, 106.13, 101.85, 86.15, 86.08, 85.59, 85.47, 55.64, 45.58, 41.64, 31.37, 21.99, 21.02, 20.38, 18.94, 18.88. ESI-MS ($CH_3OH$): calcd for [**RuN**-$I^-$]$^+$ m/z = 773.61, found m/z = 774.11.

[($\eta^6$-p-cym)Ru(L3)I]I (**Ru-Coumarin-3-carboxylic acid**, **RuC**). Complex **RuC** was obtained as yellow powder. Yield: 49.0 mg (57%). $^1$H NMR (400 MHz, $d_6$-DMSO) δ (ppm): 9.44 (*t*, *J* = 6.1 Hz, 1H), 9.38 (*d*, *J* = 6.0 Hz, 1H), 9.32 (*d*, *J* = 5.9 Hz, 1H), 8.90 (*d*, *J* = 0.6 Hz, 1H), 8.55 (*d*, *J* = 1.8 Hz, 1H), 8.50 (*d*, *J* = 1.7 Hz, 1H), 8.01 (*dd*, *J* = 7.8, 1.6 Hz, 1H), 7.79 (*ddd*, *J* = 8.8, 7.4, 1.6 Hz, 1H), 7.68-7.60 (*m*, 2H), 7.57 (*d*, *J* = 8.4 Hz, 1H), 7.47 (*td*, *J* = 7.6, 1.1 Hz, 1H), 6.23-6.11 (*m*, 2H), 6.07-5.96 (*m*, 2H), 4.81 (*d*, *J* = 6.1 Hz, 2H), 2.70 (*p*, *J* = 6.8 Hz, 1H), 2.58 (*s*, 3H), 2.38 (*s*, 3H), 0.98 (*dd*, *J* = 9.6, 6.8 Hz, 6H). $^{13}$C NMR (101 MHz, $d_6$-DMSO) δ (ppm): 162.77, 160.66, 156.90, 156.50, 154.46, 154.33, 154.10, 153.14, 152.31, 148.11, 134.88, 130.73, 128.69, 125.78, 125.65, 124.83, 122.12, 119.38, 118.73, 116.65, 106.67, 101.84, 86.22, 86.12, 85.63, 85.48, 42.38, 31.40, 22.04, 22.01, 21.07, 20.38. ESI-MS ($CH_3OH$): calcd for [**RuC**-$I^-$]$^+$ m/z = 733.5, found m/z = 734.04.

[($\eta^6$-p-cym)Ru(L4)I]I (**Ru-Benzoic acid**, **RuB**). Complex **RuB** was obtained as brown powder. Yield: 61.0 mg (77%). $^1$H NMR (400 MHz, *d*$_6$-DMSO) δ (ppm): 9.37 (*d*, *J* = 5.9 Hz, 1H), 9.28 (*dd*, J = 12.5, 6.6 Hz, 1H), 8.59 (*s*, 1H), 8.52 (*s*, 1H), 7.95 (*d*, *J* = 7.5 Hz, 2H), 7.70-7.44 (*m*, 5H), 6.12 (*t*, *J* = 5.5 Hz, 2H), 5.98 (*d*, *J* = 6.1 Hz, 2H), 4.73 (*t*, *J* = 6.5 Hz, 2H), 2.70 (*p*, *J* = 6.6 Hz, 1H), 2.58 (*s*, 3H), 2.37 (*s*, 3H), 0.97 (*dd*, *J* = 10.3, 6.8 Hz, 6H). $^{13}$C NMR (101 MHz, *d*$_6$-DMSO) δ (ppm): 167.59, 157.03, 156.51, 154.43, 154.07, 153.43, 153.32, 133.92, 132.31, 129.03, 128.70, 127.79, 125.53, 124.84, 122.42, 106.69, 101.78, 86.19, 86.08, 85.68, 85.51, 42.30, 31.39, 22.05, 21.08, 20.36. ESI-MS ($CH_3OH$): calcd for [**RuB**-$I^-$]$^+$ m/z = 665.47, found m/z = 666.05.

[($\eta^6$-p-cym)Ru(L5)I]I (**Ru-Formic acid**, **RuF**). Complex **RuF** was obtained as rust red powder. Yield: 55.0 mg (77%). $^1$H NMR (400 MHz, *d*$_6$-DMSO) δ (ppm): 9.38 (*d*, *J* = 5.9 Hz, 1H), 9.30 (*d*, *J* = 5.8 Hz, 1H), 8.72 (*t*, *J* = 6.1 Hz, 1H), 8.52 (*d*, *J* = 1.8 Hz, 1H), 8.46 (*d*, *J* = 1.8 Hz, 1H), 8.28 (*d*, *J* = 1.4 Hz, 1H), 7.61 (*dd*, *J* = 6.0, 1.7 Hz, 1H), 7.59-7.51 (*m*, 1H), 6.19-6.08 (*m*, 2H), 6.06-5.90 (*m*, 2H), 4.58 (*d*, *J* = 6.1 Hz, 2H), 2.68 (*p*, *J* = 6.9 Hz, 1H), 2.59 (*s*, 3H), 2.39 (*s*, 3H), 0.96 (*dd*, *J* = 6.9, 5.3 Hz, 6H). $^{13}$C NMR (101 MHz, *d*$_6$-DMSO) δ (ppm): 162.73, 156.89, 156.51, 154.55, 154.05, 152.86, 152.34, 128.71, 125.59, 124.87, 121.91, 106.66, 101.92, 86.25, 86.15, 85.55, 85.45, 31.38, 22.02, 21.04, 20.44. ESI-MS ($CH_3OH$): calcd for [**RuF**-$I^-$]$^+$ m/z = 589.46, found m/z = 590.02.

### *Antimicrobial assays*

#### Minimum inhibitory concentration (MIC) measurement

MIC for *M. smegmatis* or *M. fortuitum*

The MIC test was conducted using a standard microdilution broth method.[2] *M. smegmatis* or *M. fortuitum* were grown to log-phase and the culture was diluted using a fresh 7H9-tween 80 (plus 10% ADC) medium to generate the working solution containing $5\times10^5$ CFU/mL bacteria. 100 μL of this working suspension was added into each well of a 96-well plate. The solution of compounds of interest was added into the wells of the first row and a 2-fold serial dilution was performed in each column. The wells containing only 7H9-tween 80 (plus 10% ADC) were serving as the negative controls. The wells containing only the working suspension were serving as the positive controls. The 96-well plate was incubated at 37 °C with 220 rpm shaking for 48 h. $OD_{600}$ values were read on a microplate reader.

MIC for *M. marinum*

Log-phase *M. marinum* was diluted using 7H9-tween 80 (plus 10% ADC) medium to give the working suspension ($5\times10^5$ CFU/mL). 100 μL of this working solution was transferred into every well of a flat-bottom 96-well plate, except for the first column (which was reserved for 7H9-tween 80 (plus 10% ADC) as negative control) and the first row (reserved for initial compound solutions). Compounds were added at specific starting concentrations with a total volume of 200 μL into the first row. The compounds were then sequentially diluted two-fold across 8 wells with working solution. The 96-well plate was incubated without shaking at 32 °C for 48 h. Then, resazurin (0.2%, 20 μL) was added into each well and further incubated for 2 h. The color of resazurin was changed from blue to pink, indicating the growth of bacteria[3]. The columns with no color change (blue resazurin remained unchanged) were scored as above the MIC value. Data were obtained in three or more independent experiments.

MIC for *M. tuberculosis*

Note: Experiments involving the use of *M. tuberculosis* must be performed in a laboratory that shall at least meet the standard of Biosafety Level 3 (BL3).

*M. tuberculosis* were grown in a modified L-G base medium for 2 weeks and the bacteria colonies were scraped off from the agar and transferred to a centrifuge tube with 2 mL of normal saline. Glass beads were added to homogenize the colonies on a vortex. The resulting homogenized bacterial solution was diluted in fresh Middlebrook 7H9 media (plus 10% ADC supplement, Sigma: M0553-1VL; 0.5% glycerol; 0.05% Tween 80) to generate a working solution with turbidity of approximately 0.1. Compounds were diluted by serial dilution and transferred into the wells of a flat-bottom 96-well plate. Then, the bacterial working solution was added in each well to a total volume of 110 μL (10 μL of compound solution and 100 μL of working solution). The 96-well plates were incubated at 37 °C without shaking for 7 days. Resazurin[3] (0.2%, 20 μL) was then added to each well and incubated at 37 °C for 24 h. The color change of resazurin from blue to pink indicated the growth of bacteria. The MIC value was determined visually as the minimum concentration of compound that prevented color changes.

#### Time-kill assay

Log-phage *M. smegmatis* was diluted to ~$1\times10^6$ CFU/mL in 7H9-tween 80 (plus 10% ADC). And the working suspension was treated with compounds of different concentrations. At indicated time point, aliquots of the mixtures were taken out and diluted 1000-fold using PBS, and 5 μL of each was dropped onto an agar plate. After 48 h cultivation at 37 °C, colonies were count, and CFU/mL was calculated.

**Time–kill studies of *M. tuberculosis***

*M. tuberculosis* solution was diluted in sterile normal saline to prepare a working solution with $OD_{600}$ of approximately 0.5. This working solution was treated with **RuR3** at different concentrations and then incubated without shaking. At an indicated time point, 5 μL culture was taken and diluted with sterile normal saline by a 10-fold serial dilution. Then, these samples were placed onto 7H10 (plus 10% OADC) agar plates. After incubation for 1 week at 37 °C, CFU was counted and CFU/mL was calculated.

**Dormant bacteria killing assay**

$5\times10^8$ CFU/mL log-phage *M. smegmatis* were washed and resuspended in nutrient-deplete conditions (PBS). After treatment with the compound of interest for the required duration, *M. smegmatis* was diluted by 10-fold serial dilution in PBS and then plated on a LB agar plate to quantify bacterial viability.

**Bacterial persister killing assay**[4]

Persistent *M. smegmatis* were generated by treating the bacterial suspensions with 10×MIC rifampicin[5]. In brief, $1\times10^8$ CFU/mL *M. smegmatis* were treated with 80 μg/mL rifampicin for 5 h. Then, **RuR3** at different concentrations was added. At each designated time point, an aliquot (10 μL) of the suspension was diluted by 10-fold serial dilution and then was plated onto LB agar plates. After 48 h cultivation at 37 °C, colonies were counted. The CFU/mL was calculated.

**The evaluation for multistep resistance generation**

The method for multistep resistance evolution was adapted from reported procedures[6]. Briefly, the broth microdilution method for MIC determination against *M. smegmatis* was repeated for 20 passages over a period of 40 days. The initial inoculum was $5\times10^5$ CFU/mL in 7H9-tween 80 (plus 10% ADC) medium. For each subsequent passage, the inoculum for MIC determination was adjusted to a final density of approximately $5\times10^5$ CFU/mL using the content of a well containing the compound of interest at a sub-inhibitory concentration (at which bacterial growth was observed from the previous passage). To measure the MIC of each passage, bacteria were transferred to a new 96-well microtiter plate. The compound was added in triplicate to wells in the first row of the microtiter plate and then was diluted by serial dilution. The plate was incubated at 37 °C for a minimum of 48 h before the MIC was determined by reading $OD_{600}$ values. Resistance was induced when a 4-fold or greater increase compared to the initial MIC.

The resistance generation of $H_2O_2$ was reported in reference[7-8], displaying consistent results as this report (Figure1a).

**Cross-resistance assay**

The bacterial working solution for MIC determination was papered by using drug resistant strains, which were obtained from the multistep resistance generation evaluation assays described above.

**DNA or ROS scavenger interference study**

Calf thymus DNA (CT-DNA) and ROS scavenger of interest (Vitamin C, VC) purchased from Sangon Biotech were used for this study. An overnight cultured *M. smegmatis* was centrifuged and washed by PBS and then it was diluted with 7H9 medium (within 10% ADC) to $2\times10^9$ CFU/mL and was used as the working solution.150 μL of this working solution was added into each well of a 96-well plate. **RuR3** or INH with 1 mM CT-DNA or 2 mM VC were added into the designated wells. The mixtures were incubated at 37 °C with shaking (220 rpm). At specific times, 5 μL samples were taken and diluted with PBS by serial dilution at $10^4$-fold and spot-plated onto LB agar plates. The plates were incubated for 48 h at 37 °C. The colonies were counted to enumerate the number of cells.

### *Toxicity assays*

#### Cytotoxicity evaluation of RuR3 to RAW 264.7 and HEK293

Cells were seeded into 96-well plates (BIOFIL, catalog number TCP-011-096) at $1\times10^4$ cells per well and incubated for 12 h at 37 °C in 5% $CO_2$ to facilitate attachment. Cells were treated with different concentrations of **RuR3** in DMEM with 10% FBS and then incubated for 24 h. Cells with no compound added were served as the control. After incubation, the old media were removed, and cells were washed with PBS once before cell media were replaced with 120 μL of fresh media with MTT (0.5 mg/mL). Cells were incubated for another 1.5 h at 37 °C in 5% $CO_2$. Then, the medium was replaced with 100 μL of DMSO. The cell viability was determined by measuring the absorbance at 600 nm. Cell viability values were expressed as percentages and calculated as follows: Viability% = [$Abs_{600}$ of treated sample) / [$Abs_{600}$ of control] × 100%.

#### Hemolysis assay

Fresh sheep blood was subjected to a 25-fold dilution with PBS buffer to reach a concentration of 4% blood. 150 μL of PBS solution containing **RuR3** at various concentrations was placed in 96 well plate, followed by addition of equal volume of red blood cell suspension (150 μL). The mixture was incubated at 37 °C for 1 h to allow hemolysis to take place. At the end of the incubation period, non-hemolyzed red blood cells were separated by centrifugation at 300 g for 10 min. Aliquots (100 μL) of the supernatant were transferred to a 96-well plate, and hemoglobin release was measured by absorbance at 576 nm using a microplate reader (TECAN, Switzerland). Two controls were used: an untreated red blood cell suspension in PBS as the negative control; a solution containing red blood cells lysed with 0.2% Triton-X100 as the positive control. Percent hemolysis was calculated by using the following formula: Hemolysis (%) = [($OD_{576}$ of the treated sample − $OD_{576}$ of the negative control) / ($OD_{576}$ of positive control − $OD_{576}$ of negative control)] ×100%.

### *DNA-interacting related assays*

#### Hoechst/EB displacement study

Fluorophotometric competition screening and investigations of compounds' binding mode were carried out using Hoechst 33342 (a groove-binding molecule) or ethidium bromide (EB, an intercalation molecule). Binding was performed in PBS buffer (10 mM, pH 7.4) at 25 °C. 10 μM CT-DNA was pre-incubated with 4 mM Hoechst 33342 or 4 μM EB, and compounds of interest at indicated concentrations were added. For natural product library screening, 50 μM of each NP was used. After a 30-min incubation to reach equilibrium, the fluorescence was recorded by fluorescence spectroscopy. The percentage of Hoechst/EB displacement was calculated as indicated in Supplementary Figure 1a. The $IC_{50}$ values for the compounds of interest were obtained by using dose-response curves with the percentage displacement data fitted into a rectangular hyperbolic dose–response function in OriginPro 8.5 (OriginLab, Northampton, MA). The $K_i$ (binding affinity) values of compound-DNA interaction were calculated as follows:

$$K_i = \frac{IC_{50}}{1 + \frac{[4\mu M]}{K_d}}$$

$K_d$ (dissociation constant) was $2.5\times10^{-9}$ M for Hoechst 33342[9] and $1\times10^{-7}$ M for EB[10], respectively. The experiments were performed at least in triplicate.

#### Electrophoretic mobility shift assay

DNA was incubated with the compound of interest (128 μM) for 30 min at 37°C in PBS buffer (10 mM, pH 7.4). For plasmid DNA pCold-ctx-m-15 (~6500 bp) or sheared DNA (~45 bp), the mixture was run on a 1% agarose gel with Gel-Red (2 μg/mL) for 35 min. The DNA band shift was visualized and imaged under ultraviolet light using ChemiScope 6100 (Clinx). The $K_d$ value for Gel-Red is $5.6\times10^{-8}$ M[11]. For carboxyfluorescein (FAM)-labeled DNA (5'-TCACGGTACATCCTCACAGAA-FAM-3'), the mixture was run on 12% polyacrylamide gel in 1×TBE buffer solution (10 mM $Mg^{2+}$) at 120 V for 1 h. The DNA band shift was visualized and imaged with excitation wavelength of 480 nm using Fusion FX spectra (Vilber). Grayscale analysis was carried out by ImageJ software.

#### ESI-MS analysis for DNA-compound complex

Single strand DNA (ssDNA) purified by HPLC were purchased from Sangon Biotech. The solutions for ESI-MS measurement were prepared as follows: ssDNA stock solutions (100 μM) were annealed in 1x annealing buffer for DNA Oligos (Solarbio) by PCR instrument (Bio-rad, T100 Thermal Cycler) to form double-stranded DNA (dsDNA, 50 μM). Then, the dsDNA (25 μL) were mixed with 500 μM **RuR3**, $Ru^{II}$-arene, or *cis*-platin, at 37 °C for 24 h to prepare the DNA-compound complex. The resulting mixture was diluted with spray solvent (50/50 v/v MeOH/annealing buffer) to 100 μL, and was subjected to ESI-MS spectrometric analysis with negative ion mode. The DNA sequence used in this experiment is 5'-ATACATGGTACATA-3'.

#### *In vitro* DNA replication assay

Plasmid pCold-ctxm-15 (20 μg/mL) was co-incubated with the compound of interest at 128 μM for 2 h at 37 °C. The PCR amplification procedure was carried out by using gold mix (TSE101, tsingke), with the treated plasmid as a template. The *ctxm-15* gene was cloned with a PCR procedure as following: 98 °C DNA denaturation, 55 °C primers annealing and 72 °C fragment extension. Finally, the amplified products were analyzed by agarose gel electrophoresis (1% w/v in 1×TAE buffer) loading Gel-Red (2 μg/mL). The data were recorded using ChemiScope 6100 (Clinx). Grayscale analysis was carried out by ImageJ software.

Forward primer: TCGGTACCCTCGAGGGATCCGAATTCATGATGTTCGCGGCGGCGGC.
Reverse primer： CTATCTAGACTGCAGGTCGACAAGCTTTTACAGCCCTTCGGCGATGATTC TCGC-3’.

### *ROS generation related assays*

#### *In vitro* detection of exogenous ROS with EPR

20 µl **RuR3** at 250 µM in PBS buffer was prepared. The ability of **RuR3** to generate $^1O_2$ was detected by 2,2,6,6-tetramethyl-4-piperidinone hydrochloride (TEMP), using 2 µL 0.1% methylene blue (MB, 10 mM) with 15 minutes irradiation at visible light as positive control[12]. The ability of **RuR3** to generate •OH was detected by 5,5-dimethyl-1-pyrroline N-oxide (DMPO), using 100 mM $H_2O_2$ with 10 minutes UV irradiation at 365 nm as positive control[13].

Samples supplied with 100 mM TEMP or DMPO were immediately transferred to a capillary plugged with sealing putty (TERUMO CORPORATION, Tokyo Japan). Then, the capillary was put into EPR tube (WILMAD QUARTZ (CFQ), DIAM. 5MM) before it was placed in the TE mode cavity. The EPR spectra were recorded on an X-band EPR spectrometer (JES-FA200, JEOL, Tokyo, Japan). The EPR experiments were performed at room temperature with following parameters: microwave frequency: 9200 MHz; microwave power: 0.998 mW; center magnetic field: 336 mT; field sweep width: ±5 mT; time constant: 0.03 s; field modulation width: 0.05 mT. The EPR data acquisition was controlled by the WIN-RAD EPR Data Analyzer System. The spectra were simulated by JEOL IsoSimu/Fa Version 2.2.0 isotropic simulation program.

#### *In vitro* detection of exogenous ROS with DCFH

The ability of **RuR3** or ROSup (from Reactive Oxygen Species Assay Kit) in generating ROS under *in vitro* conditions was investigated. ROS generation assays were carried out by using 2,7-dichlorodihydrofluorescein (DCFH) as the probe. Briefly, **RuR3** or ROSup at various concentrations was mixed with 10 µM DCFH in 200 µL PBS, and the resulting mixture was incubated at 37 °C for 30 min. The fluorescence emission spectra of the mixture were subsequently recorded ($\lambda_{ex}/\lambda_{em}$ = 488 nm/500–560 nm, slit-widths of 5.0 nm for both excitation and emission wavelengths) with a fluorimeter (F-7000 spectrofluorometer, Hitachi). Control experiment was conducted with DCFH in PBS without adding compounds. The results reported in the present study are the average of two independent trials.

#### *In cellulo* ROS detection with live *M. smegmatis* and RAW 267.4 cells

*In cellulo* ROS generation upon compound treatment was detected for *M. smegmatis* and RAW 267.4 cells, with confocal visualization and flow cytometric quantification.

For *M. smegmatis*, overnight culture with $OD_{600}$ = 0.5 were washed with PBS for three times and resuspended in PBS. The suspension was diluted to $OD_{600}$ = 0.1 and then was mixed with 2',7'-dichlorodihydrofluorescein diacetate (DCFH-DA, 10 µM) or dihydroethidium (DHE, 5 µM), and the resulting solution was used as the working solution. Bacteria were placed into a 96-well plate (200 µL per well) and treated with the compound of interest at different concentrations for 1 h at 37°C in the dark. Confocal visualization of ROS accumulation imaged using a Nikon A1R MP confocal microscope with a 60× objective lens (DCFH-DA: $E_x$ = 488 nm, $E_m$ = 500-550 nm). Flow cytometric analysis of ROS was performed by using flow cytometry (BD Accuri C6 Plus; DCFH-DA: $E_x$ = 488 nm ± 30 nm, $E_m$ = 533 ± 30 nm; DHE: $E_x$ = 488 nm ± 30 nm, $E_m$ = 585 ± 40 nm). For each sample, 10,000 cells were counted. The geometric means were analyzed by FlowJo 10 software (Tree Star, OR, USA).

For RAW 267.4 cells, $5\times10^5$ cells were seeded into each well of a 6-well plate. The plate was incubated at 37 °C with 5% $CO_2$ for 12 h before use. The cells were treated with the compound of interest at different concentrations for 1 h at 37°C in the dark. DCFH-DA (10 µM) was added into each well, and the samples were incubated for 30 min. The cells were washed with PBS for imaging by using a Nikon A1R MP confocal microscope with a 60× objective lens. Cells were detached, centrifuged, washed, re-suspended with DMEM, and analyzed with a BD C6 Plus flow cytometer. Cells treated with PBS were used as control. For each sample, 10,000 cells were counted. The geometric means were analyzed by FlowJo 10 software (Tree Star, OR, USA).

#### *In cellulo* ROS detection with fixed *M. smegmatis*

Overnight cultured *M. smegmatis* ($OD_{600}$ = 0.5) was washed three times with PBS and resuspended in 4% paraformaldehyde for a 6-h fixation. Then, the samples were collected and resuspended in PBS for ROS detection as described above.

#### RT-PCR study

For each experiment, PBS or the compound of interest was added to different groups of *M. smegmatis* ($N$ = 3). The samples were incubated with the compound at 0.25×MIC. The RNA of *M. smegmatis* were extracted with the HiPure Bacterial RNA Kit (Magen, Shanghai), following the manufacturer's instructions. The extracted RNA was converted to cDNA. The qPCR was conducted in a real-time PCR system (QuantStudio 7 Flex, Thermo Fisher Scientific, USA) with the HiScript II One Step qRT-PCR SYBR Green Kit (Vazyme, Nanjing). Gene expression was normalized to the expression of the housekeeping gene (16S rRNA and *gapA*). The primer sequences for each gene were listed as following:

| Gene name | Sequence (5'-3') | |
|---|---|---|
| | Forward | Reverse |
| *16S* | ACAAACGCGACAAGACCGG | CAGTAGTGAGCCGCTCGT |
| *gapA* | AAGATCAAGGCCCTCGAGG | GGCGTTGGAGATGATGTTCT |
| *ahpc* | CGTCACCAGCAAGGATTACG | CTTGAGGTCCTCGTGCTGT |
| *katG* | ATCTGCTGGTGTTCACCGG | CATCGTGGTCGCACCGTA |
| *SOD* | AGAACTCAAGACCGCCGAC | GCCGGACACCTGGAAGTG |
| *mazF* | ATATCTACACCGCGGCGGC | GAGCTGACCGTGGAGAGTCC |
| *mazE* | CCGAGTACGCCGACATCGC | GCGTCGTCCCAGTCGACG |
| *oxyR* | GTACTTCGTCGCCGTGGC | CTCTTGGCCCACACCACGA |
| *recA* | AAAGCTGGGTGTGGACACC | GTGGTCCCCGAGTTGTTCAA |
| *dinB* | ATGCGCAACCATTCACACTT | GCAGATCATCCTCGCTCAAC |
| *uvrA* | CTGCTCAAGGTGATCAACGG | TGATCCAGTCGGAGGTCTTG |

### *Detection of oxidative DNA damage*

#### 8-OHdG quantitation assay for *M. smegmatis*, RAW 264.7 and CT-DNA

Sample preparation

For *M. smegmatis*, the overnight culture was diluted 1:100 into 20 mL 7H9-tween 80 (plus 10% ADC) medium in 250 mL baffled flasks and grown to an $OD_{600}$ of 0.4. Cells were then treated with 16 μM **RuR3**, or 20 mM $H_2O_2$, for 5 h. After that, the cells were collected and spun down at 10000 g for 2 min in a benchtop swinging bucket centrifuge. Genomic DNA was extracted using the Genomic DNA Mini-Extraction Kit (Universal, Spin Column).

For RAW 264.7, $1\times10^6$ cells were exposed to 16 μM **RuR3** and 1 mM $H_2O_2$. After incubating for 1 h, the samples were centrifugated at 300 g for 5 min. The supernatants were collected and resuspended in PBS. The cells were lysed by ultrasonic immersion probe sonication (1 min, 20% power) for the determination of DNA oxidation.

For CT-DNA, DNA were dissolved in sterile $H_2O$ at a concentration of 500 μM and were treated with compound of interest or UV irradiation at TL-D 18W (Philips).

8-OHdG quantitation

8-OHdG levels were quantified with an 8-OHdG (8-hydroxydeoxyguanosine) ELISA Kit (E-EL-0028c, Elabscience). Samples were assayed in three replicates. Sensitivities of the ELISA kits are 0.94 ng/mL.

#### Detection of DNA double strand break with Gam-GFP fusion protein[14]

Gam-GFP plasmid construction

Gam-GFP plasmid was generated by homologous recombination with standard protocols. Briefly, the target gene were amplified with forward and backward primers, purified, and ligated into the corresponding vectors with recombinase.

Vectors: pMV261-Gam-Linker-EGFP
Restriction sites: BamH1, Hind3
Forward primers: CAATGGCCAAGACAATTGCCGGATCCATGGCCAAACCAGCAAAACG
Backward primer: TCCTCGCCCTTGCTCACCATGGTGGCGACCGGTGGATCCGGACTG
CCACCACCACCAC

Gam-GFP Assay

Gam-GFP plasmid was electric-transferred into *M. smegmatis*. *M. smegmatis* containing the Gam-GFP plasmid were grown to log-phase in the presence of 50 μg/mL kanamycin. The cells were treated with 16 μM **RuR3** or 20 mM $H_2O_2$, respectively, and incubated at 37 °C with shaking at 220 rpm for 5 h. After that, the bacteria were collected and re-suspended in 5 μL/mL of DAPI/PBS and incubated at room temperature in the dark for 15 min. The bacteria were centrifuged again and then re-suspended in 100 μL PBS. To image cells, 2 μL of the cell culture was placed on a slide and sealed with a coverslip. Imaging was performed on Nikon Laser Confocal Microscope with a 60× objective lens. Quantitative analysis of Gam-GFP foci was performed using ImageJ software.

**Gel electrophoresis assay to evaluate genomic DNA integrity**

Evaluation of genomic DNA integrity was carried out on both single-cell level[15] and ensemble level. For compound treatment, log-phase *M. smegmatis* were treated with 16 μM **RuR3**, or 20 mM $H_2O_2$ (positive control) or 20 μM ethambutol for 5 h. Then, the cells were collected and lysed with lysozyme and proteinase K. Lysate containing genomic DNA was mixed with 2% molten low-melting point agarose ("genomic DNA mixed agarose").

For single-cell level electrophoresis, the genomic DNA mixed agarose was spread on slides precoated with 1% normal-melting agarose. After solidification at 4 °C, the slides were immersed in an alkaline lysis buffer at 4 °C for further lysing. Then, the slides were rinsed with PBS and transferred into running buffer (1×TBE). Electrophoresis was carried out at 12 V and 40 mA for 30 min. Slides were washed with PBS and then stained by using Gel-Red (2 μg/mL). Samples were imaged with Nikon Laser Confocal Microscope with a 60× objective lens.

For ensemble-level electrophoresis, the genomic DNA mixed agarose was loaded into a conventional agarose gel, and electrophoresed in 0.5×TBE at 75 V for 8.5 h. The gel was then stained by using a Gel-Red (2 μg/mL) solution for 30 min on a shaker (60 rpm). The gel was imaged using ChemiScope 6100 (Clinx). As a control, extracted genomic DNA of *M. smegmatis* was examined with the above protocol, to investigate the effect of live cell metabolism on DNA oxidation.

### *Transcriptome analysis*

1. Sample preparation

An overnight starter culture in 7H9-tween 80 (plus 10% ADC) of *M. smegmatis* MC2 155 (ATCC 700084) was diluted 50-fold in a fresh medium. The control group and treatment group with 1 μM (0.5×MIC) **RuR3** were grown for 19 h and 34 h, respectively, to reach log phase. The cells were harvested by centrifuging at 3200 g for 10 min. Samples were then frozen in liquid nitrogen. The weights of the control group were found to be 160.4 mg, 148.5 mg, 162.4 mg, and 111.9 mg, 109.4 mg, respectively. 152.4 mg was obtained for **RuR3** treatment group.

2. RNA quantification and qualification

Total RNAs obtained from replicate samples of *M. smegmatis* were extracted by using commercial kits, according to the manufacturer's instructions. RNA degradation and contamination were monitored on 1% agarose gels. RNA concentrations were measured using Qubit 2.0 (Thermo Fisher Scientific, MA, USA) and Nanodrop One (Thermo Fisher Scientific, MA, USA), at the same time. The RNA integrity was determined by using an Agilent 2100 system (Agilent Technologies, Waldbron, Germany).

3. Library preparation.

Whole mRNAseq libraries were generated using NEB Next® UltraTM Directional RNA Library Prep Kit for Illumina® (New England Biolabs, MA, USA), following the manufacturer's recommendations. Briefly, bacterial and archaeal 16S and 23S rRNA transcripts in total RNA samples were reduced by using a Ribo-zero rRNA Removal Kit. Fragmentation was carried out using NEB Next First Strand Synthesis Reaction Buffer. The first strand cDNA was synthesized using random hexamer primer and M-MuLV Reverse Transcriptase (RNase H). During the synthesis of the second strand of cDNA, a chain-specific library was constructed by replacing dTTP with dUTP, to improve accuracy. Remaining overhangs were converted into blunt ends via exonuclease/polymerase reactions. After adenylation of the 3' ends of DNA fragments, NEB Next Adaptor with a hairpin loop structure were ligated to be ready for hybridization. In order to select cDNA fragments of about 150-200 bp length, fragments were selected with AMPure XP beads (Beckman Coulter, Beverly, USA). Then, PCR was performed with Phusion High-Fidelity DNA polymerase, Universal PCR primers and Index (X) Primer. Finally, PCR products were purified with AMPure XP beads and library insert size assessed by using an Agilent 2100 system (Agilent Technologies, Waldbron, Germany)

4. Transcriptome sequencing

The clustering of the index-coded samples was performed on a cBot Cluster Generation System. After cluster generation, the library was sequenced on an Illumina Hiseq Xten platform and 150 bp paired-end reads were generated.

5. Data analysis

Data analysis was performed in the following steps:

a) Quality control. Raw data in fastq format were processed by Trimmomatic (v.0.36,ing to NCBI Rfam databases, to remove the rRNA sequence, by using Bowtie2 (v2.33, https://github.com/BenLangmead/bowtie2).

b) Reads mapping to the reference genome. Reference genome and gene model annotation files were downloaded from the NCBI genome website directly. The remaining mRNA sequences were mapped to the *M. smegmatis* reference genome by using Hisat2 (version 2.1.0, https://github.com/infphilo/hisat2).

c) Transcript quantification and sample relationship analysis. HTSeq-count (v0.9.1, http://htseq.readthedocs.io/en/release_ 0.9.1/) was used to obtain the read count and function information of each gene, based on the mapping results. In order to make the expression levels of genes be comparable among different genes and different experiments, the RPKM of each gene was calculated. RPKM, Reads Per Kilobase of transcript, per Million mapped reads, is a normalized unit of transcript expression and considers the effects of sequencing depth and gene length for the read count, and is currently the most commonly used method for estimating gene expression levels. PCA (principal component analysis), correlation coefficient heat maps and expression heat maps were then used to reveal transcription relationships between all samples.

d) Differential expression analysis. Read count of each gene obtained from HTSeq-count was used for differential expression analysis. Differential expression analysis of gene expression data was performed using edgeR (v3.16.5, http://www bioconductor.org/packages/release/bioc/html/edgeR.html), which takes the length and number of genes into account. The resulting P-values were adjusted by using the Benjamini and Hochberg approach for controlling the false discovery rate (FDR). Genes with FDR $\leq$ 0.05 and $|\log2(\text{fold change})| \geq 1$ were taken as differentially expressed genes, and these were used for heatmap construction.

e) Bioinformatic analysis. GO (Gene Ontology, http://www.geneontology.org) annotation analysis of differentially expressed genes were implemented by using clusterProfiler (v3.4.4, http://www.bioconductor.org/packages/release/bioc/html/ clusterProfiler.html). Searching of sequence homologs, secondary structure prediction and protein 3D structure prediction of gene products were performed with the Protein Basic Local Alignment Search Tool (BLAST, see *J. Mol. Biol.* **1990**, *215*, 403- 410), TMHMM (see *J. Mol. Biol.* **2001**, *305*, 567-580) and Phyre2 (*Nat. Protoc.* **2015**, *10*, 845-858) servers, respectively.

### *Intracellular localization studies*

**Localization of *M. smegmatis* in macrophages**

RAW 264.7 cells were cultured in DMEM supplemented with 10% FBS at 37 °C in an incubator with a 5% $CO_2$ atmosphere. RAW 264.7 cells in 1.5 mL of medium were seeded into each well. The cells were then incubated for 12 h for attachment. The cells were infected with log-phase *M. smegmatis* pre-stained with Hoechst at multiplicity of infection 1:10 for 3.5 h. Then, the infected cells were incubated with Hoechst (10 μg/mL) for 15 min for nuclear staining. Acidic compartment was (pH~6.2) stained by Lyso-tracker red (50 nM), cells were fully immersed in 1 mL of staining solution and incubated at 37 °C in an incubator with a 5% $CO_2$ atmosphere for 30 min. The medium was removed and the cells were washed again with PBS for thrice. The cells were imaged using a Nikon Laser Confocal Microscope with a 60× objective lens.

**Localization of RuR3 in mammalian cells**

RAW 264.7 cells were seeded at a density of $3\times10^5$ cells/well in cell culture dish with a glass bottom. **RuR3** (20 μM) was added for cellular uptake. At indicated time points, samples were washed three times with PBS for imaging. Finally, the staining solution was removed and the cells were washed with PBS twice. Samples were imaged using confocal microscope.

A549 cells were seeded at a density of $5\times10^4$ cells/mL into 35 mm confocal dishes (JET BIOFIL, Canada) for confocal microscopy. After cultured overnight, the cells were pre-incubated with **RuR3** (10 μM) for 4 h at 37 °C. Subsequently, the medium was replaced with the staining medium containing Lyso-Green (2 μM) or Mito-green (200 nM) and stained in the dark for 30 min at 37 °C. The staining medium was removed and the cells were washed with PBS for thrice, and then observed immediately under a confocal microscope (A1, Nikon, Japan). **RuR3** was excited at 405 nm and an emission was collected at 605 ± 20 nm. The excitation wavelength of Mito-Green, Lyso-Green dyes was 488 nm and the emission were collected at 516 ± 20 nm and 505 ± 20 nm, respectively.

**ICP-MS measurement**

The cellular uptake and distribution of **RuR3** in RAW 264.7 was determined by measuring the Ruthenium contents. Briefly, cells were seeded and incubated overnight under standard growth condition. The culture medium was removed and replaced with fresh medium/DMSO (v/v, 99:1) containing **RuR3** (10 μM). After incubation for 24 h, the cells were harvested with trypsin, washed with PBS and counted. Cell pellets were lysed in RIPA Lysis Buffer (Beyotime Biotechnology, China) and the nucleus and mitochondria fractions were extracted using the Mitochondria/Nuclei Isolation Kit (KeyGEN BioTECH, China) according to the manufacturer's instructions. These fractions in the stock buffer were then digested with concentrated nitric acid (100 μL) at 368 K for 2 h, hydrogen peroxide (30 %, 50 μL) at 368 K for 1.5 h, and concentrated hydrochloric acid (50 μL) at 368 K for 1.5 h to give fully homogenized solutions. Finally, the solutions were diluted with MiliQ water to a final volume of 2 mL for the measurement and iridium contents in the samples were determined by ICP-MS (X Series 2, Thermo Fisher, USA). The average of three parallel experimental data was reported as the final results.

**Mechanism of mammalian cell internalization of RuR3**

RAW 264.7 cells were treated with **RuR3** under different conditions for to observe possible inhibitory effects. For the low temperature condition, the cells were precooled at 4 °C for 30 min, and **RuR3** (20 μM) was added for cellular uptake for 2 h at 4 °C. For conditions with endocytic inhibitors, the cells were pre-incubated with the corresponding medium with genistein (100 μg/mL), chlorpromazine (10 μg/mL), wortmannin (10 μg/mL), or mβCD (5 mM) for 30 min at 37 °C. **RuR3** (20 μM) was added for cellular uptake for 2 h at 37 °C. The uptake level of **RuR3** under different conditions was quantified by Nikon Laser Confocal Microscope.

**RuR3 co-localization with intracellular *M. smegmatis***

*M. smegmatis* infected RAW 264.7 (Ms⊂RAW 264.7) cells were treated with 20 μM **RuR3** for cellular uptake for 4 h at 37 °C. Then, the cells were washed three times with PBS and imaged using a Nikon Laser Confocal Microscope.

**Generation of co-localization histograms**

The indicated channels were compared using the Colocalization module (in NIS-Elements Viewer), to obtain a histogram and a corresponding Pearson's R value for each confocal image.

The indicated channels were compared using the Colocalization module (in NIS-Elements Viewer), to obtain a histogram and a corresponding Pearson's R value for each confocal image. The generated histograms were shown in Supplementary Figure 12b,15b. For each group of samples, the experiments were repeated independently for at least three times and obtained several sets of confocal images for each group. These images were analyzed as described above to give multiple histograms and Pearson's R values for each group.

**Localization of ROS in RAW 264.7 and Ms⊂RAW 264.7**

Ms⊂RAW 264.7 ($3\times10^5$) were treated with 16 μM **RuR3** or 500 μg/mL ROSup as described above. The cells were then washed with PBS, and stained with 10 μM DCFH-DA in fresh DMEM in dark. After 30 min, DCFH-DA was removed and the cells were imaged by Nikon Laser Confocal Microscope.

### *Ex vivo and in vivo antimicrobial efficacy of RuR3*

**Evaluation of intracellular antimicrobial activity**

An overnight *M. smegmatis* culture was harvested by centrifugation. The 7H9 medium used in the culture was discarded. The deposited bacteria were washed by PBS and resuspended in a fresh 7H9 medium. RAW 264.7 cells ($2\times10^5$ for imaging, $1\times10^5$ for quantification) were incubated in 90% DMEM containing 10% FBS at 37 °C with 5% $CO_2$ supplement overnight. Then, the cells were infected with *M. smegmatis* using a respective bacteria-containing medium, at a MOI of 10 for 3.5 h. Then, the extracellular bacteria were killed using gentamicin (50 μg/mL in 90% DMEM with 10% FBS) for 30 min. The infected cells were washed by PBS for three time. Then, 1.5 mL of fresh DMEM plus 10% FBS medium were added into each well, followed by the addition of compounds of interest. After incubation for 24 h, the medium was removed and the cells were washed with PBS.

For imaging, the samples were observed using a Nikon Laser Confocal Microscope. For quantification of bacterial load and mammalian cell viability, the cells were harvested and 10-fold diluted with PBS containing 0.04% trypan blue. After incubating at 37 °C for 5 min, the live cells without blue stain were counted with optical microscope. To quantify bacterial load, 10 μL medium containing *M. smegmatis* were diluted with PBS, in a 10-fold serial dilution, and dropped on an agar plate. The plates were then incubated at 37 °C for 48 h before CFU counting.

***M. fortuitum*-infected zebra fish model study**

Overnight cultured *M. fortuitum* in 7H9 medium ($OD_{600}$ = 0.6) was harvested by centrifugation and was washed by PBS. Zebra fish was divided into three groups ($N$ = 12). Then, *M. fortuitum* ($1.5\times10^8$ CFU) were intraperitoneally injected into the fish. After 30 min, **RuR3** (30 mg/kg), ciprofloxacin (30 mg/kg), ethambutol (30 mg/kg), or 50% PBS+50% PEG (co-solvent) was injected intraperitoneally into the fish in the corresponding groups. The appearance and vitality of the fish in each group were recorded at indicated time points.

**Effectiveness of RuR3 in mouse model with *M. marinum* infection**

ICR male mice used in this study with an average weight of 28 g were purchased from hunan SJA laboratory animal CO., LTD (changsha, China). To establish the mouse infection model, *M. marinum* was cultured on a 7H10 agar plate for one week. After being transferred into a 7H9 medium with static culture for 5-7 days at 32 °C, log phase *M. marinum* ($OD_{600}$ = 0.6) were collected by centrifugation, and then washed trice and resuspended in PBS to a final bacterial solution containing $2\times10^8$ CFU/mL *M. marinum*. 100 μL bacterial solution was then intravenous (i.v.) injected into the mice through a 29 Gauge needle. After 24 h, the successfully infected mice were divided into three groups: **RuR3** group ($N$ = 5), kanamycin group ($N$ = 5) and control group ($N$ = 5). For the **RuR3** and kanamycin group, a dose of 2.5 mg/kg in 100 μL of solution was applied to each mouse by i.v. injection. The same injections were performed at day 4 and day 7 drug post infection, respectively. For the control group, 100 μL of 50% PBS+50% PEG (co-solvent) were injected to the mice at the same time as in the treatment group. The tail condition of each mouse was monitored and photographed every day for 14 days.

**Immunohistology**

**RuR3** (2.5 mg/kg at day1, day 4 and day 7) and 50% PBS+50% PEG (co-solvent) were i.v. injected into the mice. The body weight of mice was monitored per day and the mice were euthanized after 17 days. Liver, lung, spleen and kidney were collected and fixed in formalin for 24 h at 4 °C. Then, paraffin embedding and stained with Hematoxylin and Eosin (H&E) by Wuhan Servicebio Technology Co. Ltd. and observed by fluorescence microscopy (Pannoramic MIDI microscope).

### *Molecular Dynamic Simulation*

#### DNA-ligand complex/adduct modeling

Molecular dynamic simulation was performed for non-covalent **RuR3**-DNA complex, non-covalent $Ru^{II}$ arene-DNA complex, and covalent **RuR3**-DNA adduct, and the Molecular Mechanics/Generalized Born Surface Area (MM/GBSA) energy was calculated between the ligand and DNA.

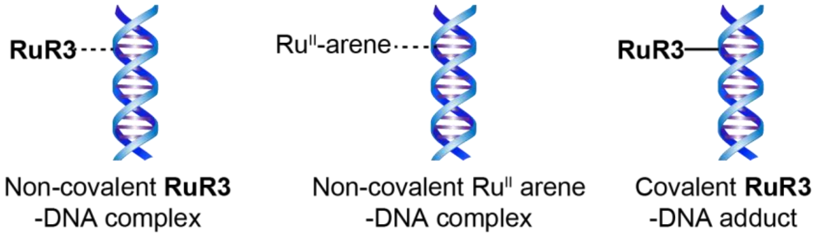


Non-covalent DNA-ligand complex

First, the molecular configurations of **RuR3** and $Ru^{II}$-arene were modeled manually according to the reported crystal structure of relevant compounds[16]. Next, the structure of a platinum-acridine-liganded octamer in the Protein Data Bank (PDB ID code 1XRW) was used as the template to model the initial the complex formed between DNA and ligand. The modeled complexes were refined by molecular dynamics (MD) simulations using the AMBER16 package. As there are no standard force field parameters available for Ru-based drugs, their force field parameters were thus prepared manually. The metal center parameter builder (MCPB.py) was used to parameterize the metal site of the Ru complex.[17] The Gaussian09[18] was used in MCPB.py to obtain the angle, bond and charge parameters according to the B3LYP density function method and the relativistic effective core pseudopotential (LANL2DZ). The OL15 force field[19] was used to parameterize the DNA molecule. Each complex was solvated in a periodic TIP3P water box and neutralized with $Na^+$ counterions. Electrostatic interactions were considered by the Particle Mesh Ewald algorithm with a cutoff of 10 Å. The SHAKE algorithm was used to constrain all the bonds involving hydrogen atoms. The systems were heated up to 300 K after an initial minimization while imposing positional restraints on the heavy atoms. Subsequently, 300 ps of simulations were conducted in the NPT ensemble without restraints, followed by annealing to 0 K in 100ps. During these simulations, the pressure and temperature were maintained using the Berendsen barostat and Langevin thermostat. The last frame of the simulation was the final structure model that to be analyzed for each complex.

Covalent DNA-**RuR3** adduct

For coordinated binding of **RuR3** to guanine, N7 of guanine base is bonded to the metal site, and the relevant parameters for Ru-N bond were obtained by MCPB.py, while the other parameters of guanine base being the same as for the OL15 force field.

#### Interaction between DNA polymerase and liganded DNA

Interactions between DNA polymerase and liganded DNA were modeled for the following samples: polymerase-DNA, polymerase-(**RuR3**-DNA adduct) and polymerase-($Ru^{II}$ arene-DNA complex), and the MM/GBSA energy was calculated between the polymerase and DNA.

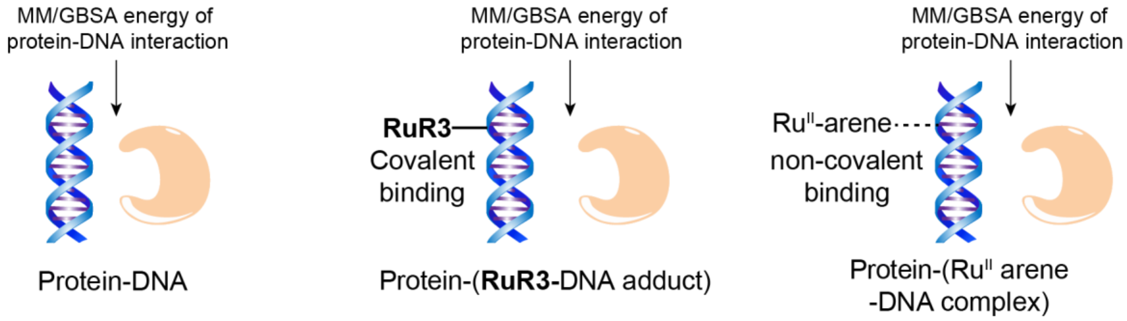


The crystal structure of a DNA polymerase I with 11 base pairs of duplex DNA (PDB ID code 1L3U), was selected to investigate the impact of $Ru^{II}$ arene or **RuR3** in the interaction between polymerase and DNA. First, DNA was separated from the DNA-polymerase structure, and $Ru^{II}$- arene or **RuR3** was docked into the DNA structure nonselectively with the previous DNA-ligand complex/adduct as reference. Next, the liganded DNA structures (covalent binding for **RuR3**-DNA adduct and non-covalent binding for $Ru^{II}$ arene-DNA complex) were refined using the same protocol as described above in “DNA-ligand complex/adduct modeling”. Third, the liganded DNA structure was embedded back into the polymerase by aligning with the original DNA. Finally, atomistic molecular simulations were performed (15 ns) for the three samples using the AMBER 16 package. The MM/GBSA binding free energies between polymerase and DNA were obtained. Specifically, each system was solvated in a periodic box of $\sim 92 \times 86 \times 104$ Å$^3$ and neutralized with $Na^+$ counterions. The system was subjected to energy minimization of 500 steps of steepest descent and 500 steps of conjugate gradient with no restraints. Then, the system was heated to 300 K at a constant volume during 100 ps (50000 steps) with harmonic restraints of 3.0 kcal mol$^{-1}$ Å$^{-2}$ on the heavy atoms. Subsequently, restraints were removed and 15 ns of unrestrained dynamics in the NVT ensemble were performed for each system. The time step was set to 2 fs and the coordinates of the system were recorded for every 5000 steps (i.e. 10 ps). Finally, the MM/GBSA binding free energy[20] between DNA and protein was calculated with 1 frame interval using the python script MMPBSA.py[21]. The results show that **RuR3** weaken the interaction between Protein and DNA.

16. Caruso, F.; Monti, E.; Matthews, J.; Rossi, M.; Gariboldi, M. B.; Pettinari, C.; Pettinari, R.; Marchetti, F., Synthesis, Characterization, and Antitumor Activity of Water-Soluble (Arene)ruthenium(II) Derivatives of 1,3-

Dimethyl-4-acylpyrazolon-5-ato Ligands. First Example of Ru(arene)(ligand) Antitumor Species Involving Simultaneous Ru-N7(guanine) Bonding and Ligand Intercalation to DNA. *Inorganic Chemistry* **2014,** *53* (7), 3668-3677.

17. Li, P.; Merz, K. M., Jr., MCPB.py: A Python Based Metal Center Parameter Builder. *Journal of Chemical Information and Modeling* **2016,** *56* (4), 599-604.

18. Frisch, M. J.; Trucks, G.; Schlegel, H. B.; Scuseria, G. E.; Robb; Cheeseman, J. R.; Scalmani, G.; Barone, V.; Mennucci, B.; Petersson, G. A. In *Gaussian 09W, revision A. 02*, 2016.

19. Galindo-Murillo, R.; Robertson, J. C.; Zgarbova, M.; Sponer, J.; Otyepka, M.; Jurecka, P.; Cheatham, T. E., III, Assessing the Current State of Amber Force Field Modifications for DNA. *Journal of Chemical Theory and Computation* **2016,** *12* (8), 4114-4127.

20. Genheden, S.; Ryde, U., The MM/PBSA and MM/GBSA methods to estimate ligand-binding affinities. *Expert Opinion on Drug Discovery* **2015,** *10* (5), 449-461.

21. Miller, B. R., III; McGee, T. D., Jr.; Swails, J. M.; Homeyer, N.; Gohlke, H.; Roitberg, A. E., MMPBSA.py: An Efficient Program for End-State Free Energy Calculations. *Journal of Chemical Theory and Computation* **2012,** *8* (9), 3314-3321.

**Supplementary Figures 1-51 and Tables 1-2 for**

**Manipulation of Bacterial ROS Production Leads to Self-escalating DNA Damage and Resistance-resistant Lethality for Intracellular Mycobacteria**

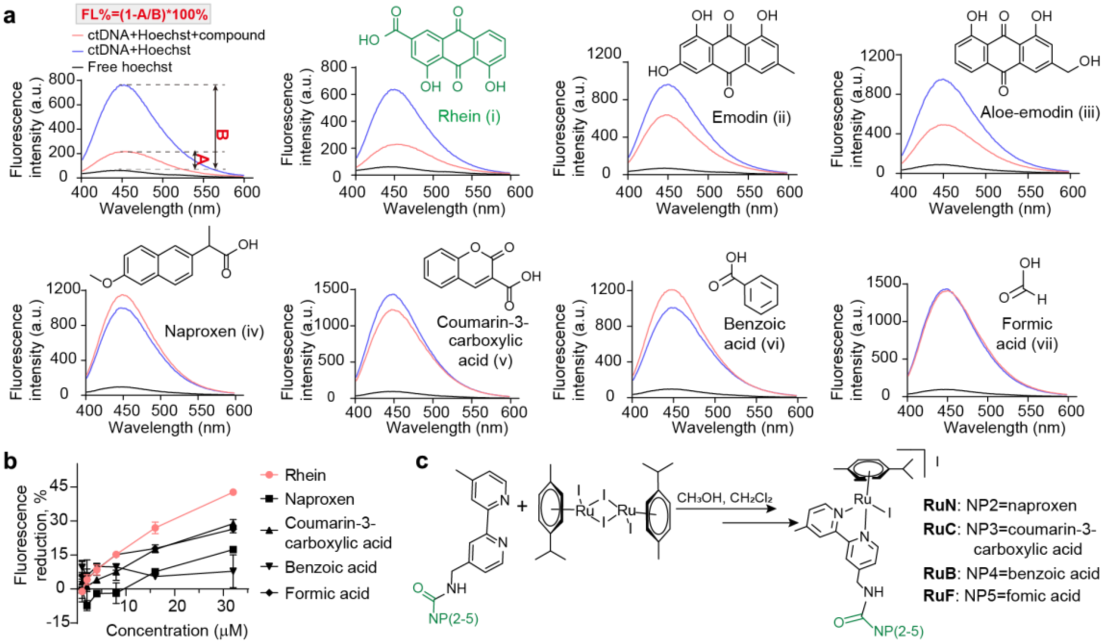


Supplementary Figure 1. (a) Schematic representation of fluorophotometric competition assay to identify minor-groove binding natural products. The percentage of Hoechst displacement= $(1-A_{\text{Fluorescence intensity}}/B_{\text{Fluorescence intensity}})\times100\%$. (b) Experimental data for fluorophotometric competition assay of Rhein, Emodin, Aloe-emodin, Naproxen, Coumarin-3-carboxylic acid, Benzoic acid and Formic acid. (c) Synthetic scheme of the **Ru$^{II}$-NP**s studied in this work.

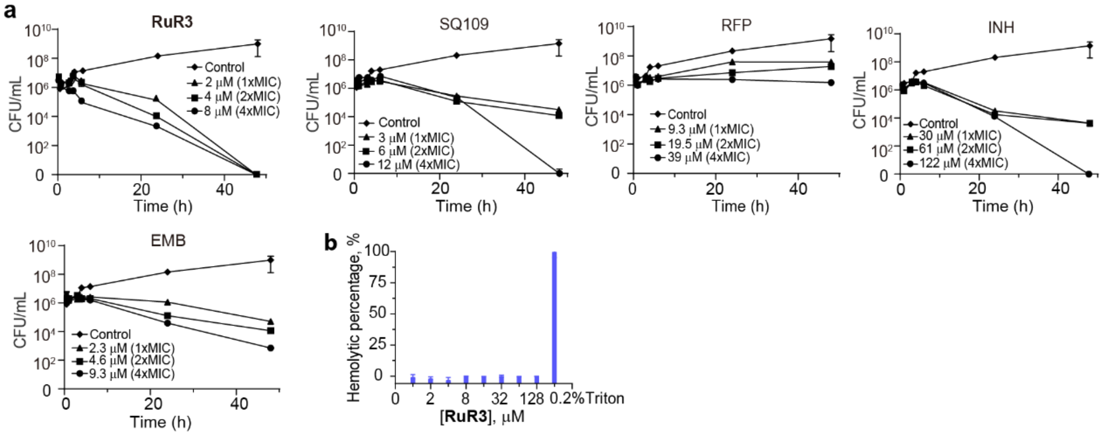


Supplementary Figure 2. (a) Killing of *M. smegmatis* in 7H9-tween 80 (plus 10% ADC) in the presence of varying concentrations of **RuR3**, SQ109, RFP, INH or EMB. The initial cell density is ~1 x $10^6$ CFU/mL. (b) Hemolytic toxicity assay of **RuR3**.

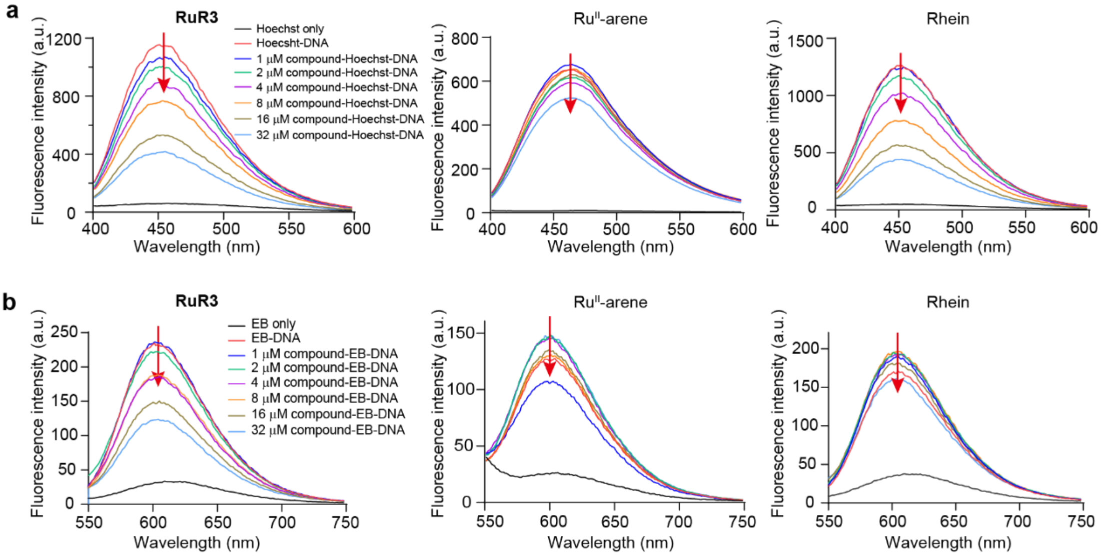


Supplementary Figure 3. Experimental data for dye displacement assay with the minor groove binding (a) Hoechst and the intercalating DNA dye (b) EB.

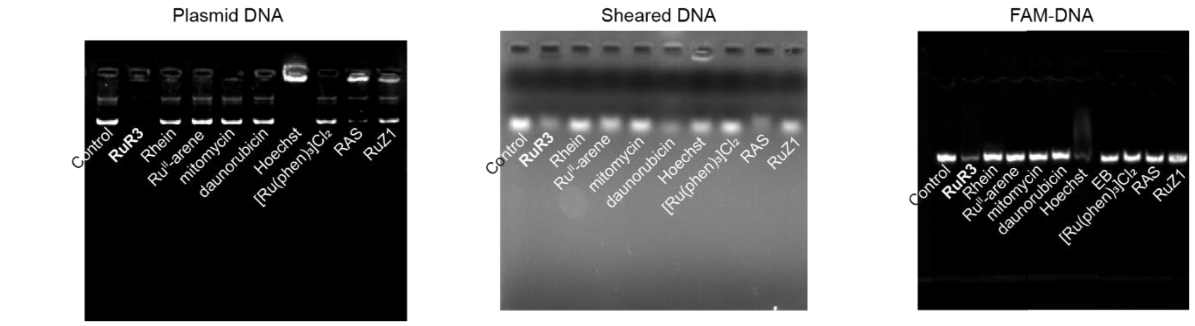


Supplementary Figure 4. Electrophoretic mobility shift assay to investigate DNA-compound interaction. Effects of 128 μM **RuR3** or control compounds were examined with three forms of DNA: Plasmid DNA (pCold-ctxm-15), Sheared Herring Sperm DNA (sheared DNA), carboxyfluorescein (FAM) labeled DNA (FAM-DNA).

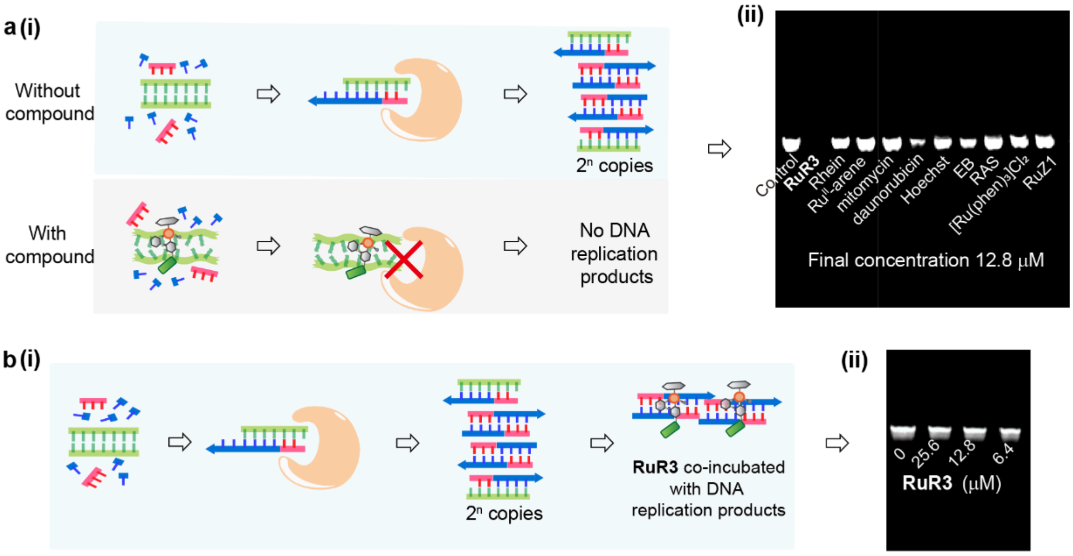


Supplementary Figure 5. (a) (i) Cartoon illustration of the PCR-based *in vitro* DNA replication assay. (ii) Inhibition of PCR product formation upon compound treatment. Template DNA was pre-treated with compound before the PCR reaction with final concentration of 12.8 μM. (b) (i) Cartoon illustration of the control experiment of the PCR inhibition assay, in which **RuR3** was added after the PCR reaction. (ii) The lack of PCR product inhibition in this assay suggests the observed PCR product inhibition in (a) originates from interruption of DNA replication, but not direct PCR product binding.

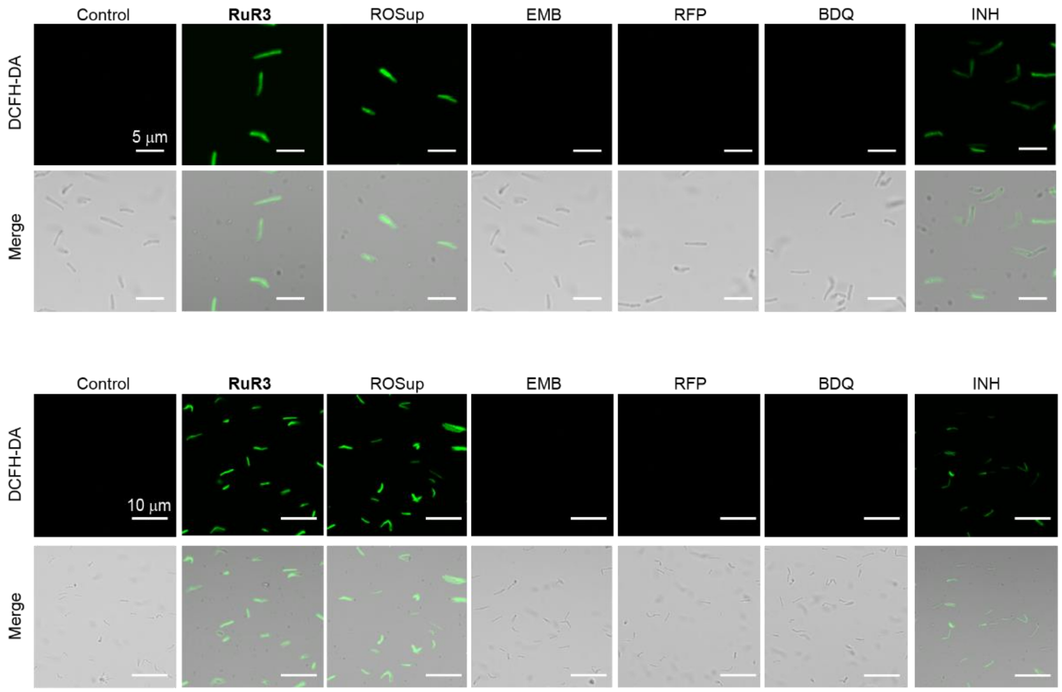


Supplementary Figure 6. Two additional data sets of confocal visualization of ROS accumulation of **RuR3**, ROSup and conventional stressors (EMB, RFP, BDQ and INH). ROS is probed by a green fluorescent dye DCFH-DA.

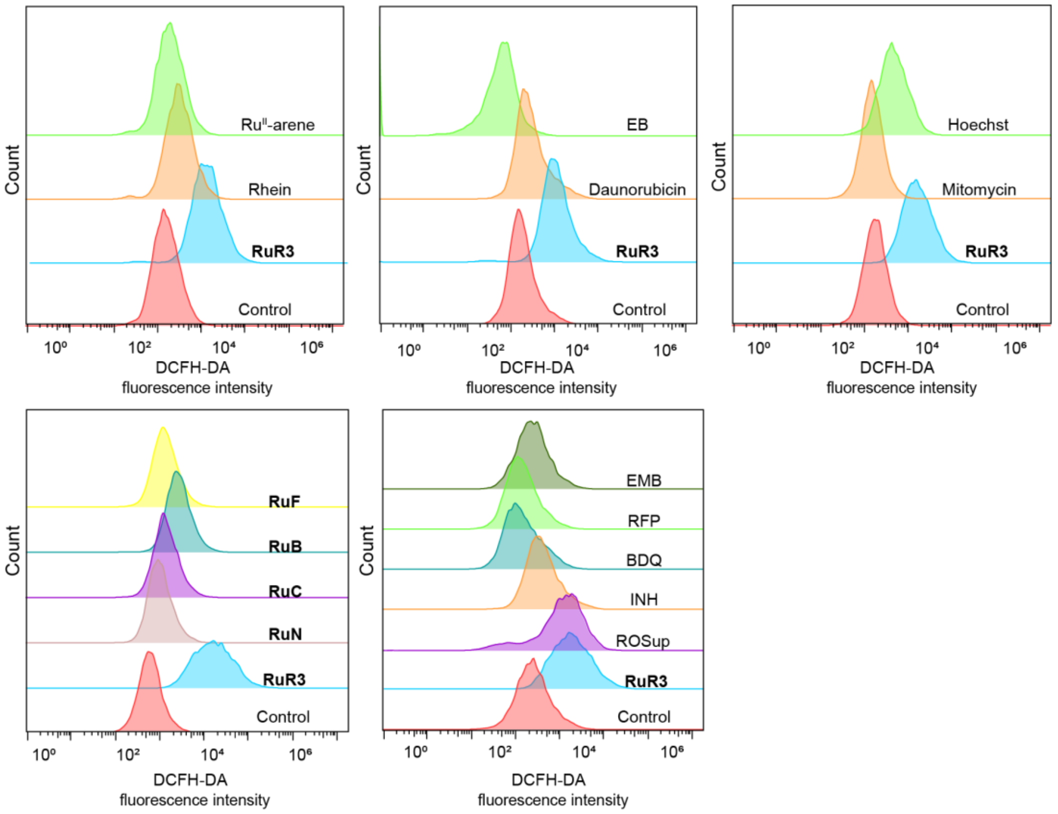


Supplementary Figure 7. Flow cytometric analysis of ROS accumulation in *M. smegmatis* upon treatment with 4×MIC compounds of interest. ROS is probed by a green fluorescent dye DCFH-DA. Relative DCFA-DA fluorescence presented in the main text figure is calculated as following (GEO=The fluorescence geometric mean of DCFH-DA):

$$\text{Relative ROS level upon compounds treatment} = \frac{\text{GEO of compound} - \text{GEO of no treatment group}}{\text{GEO of } \mathbf{RuR3} - \text{GEO of no treatment group}} \times 100\%$$

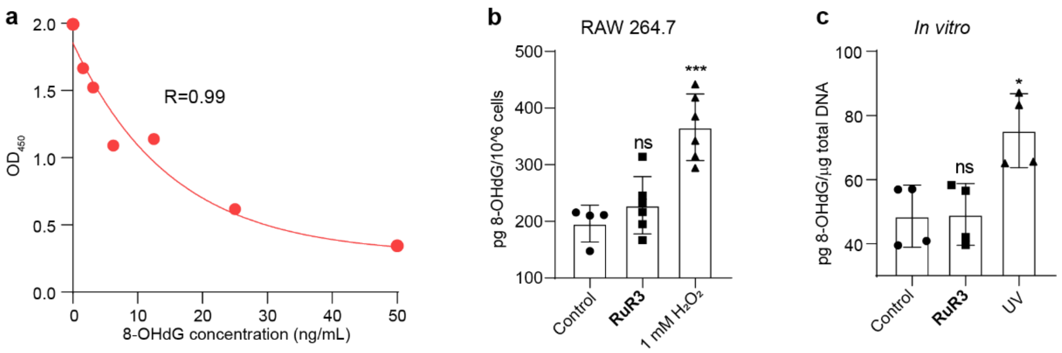


Supplementary Figure 8. (a) Standard curve of ELISA for 8-OHdG detection. (b) ELISA-based quantification of 8-OHdG level on genomic DNA of RAW 264.7 upon treatment with16 μM **RuR3** or $H_2O_2$. (c) ELISA-based quantification of 8-OHdG level on free DNA upon *in vitro* treatment with 16 μM **RuR3** or UV.

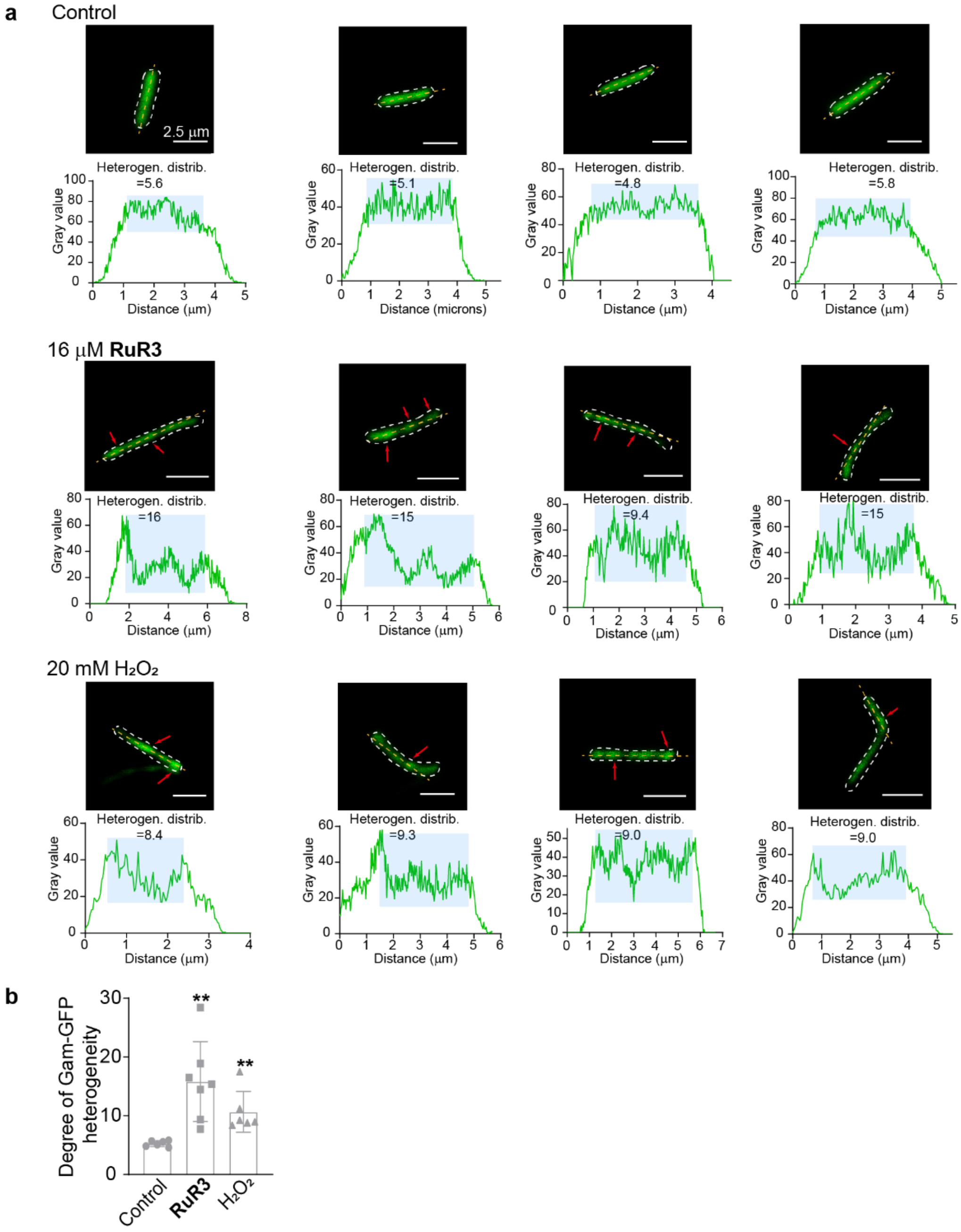


Supplementary Figure 9. (a) The additional confocal images with arrows indicating Gam-GFP foci, which occur at double-strand breaks. Heterogeneity of Gam-GFP distribution is calculated as standard deviation of Gam-GFP distribution within the bacterial cell. (b) Statistical analysis of Gam-GFP fluorescence heterogeneity in the bacterial population. (Heterogen. distrib.: heterogeneity distribution).

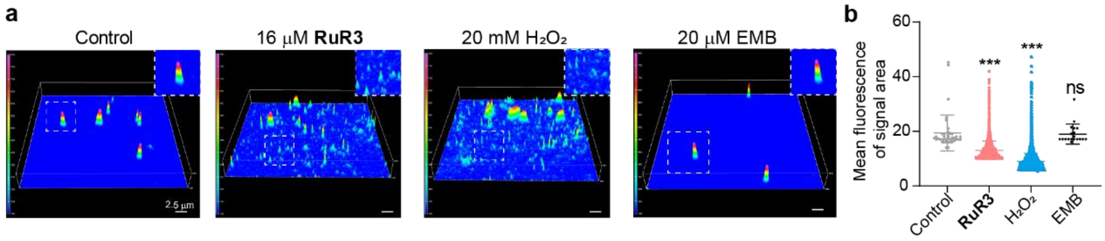


Supplementary Figure 10. The additional results of single-cell gel electrophoresis assay to examine integrity of genomic DNA. (a) Additional confocal images showing the intact genomic DNA in control and ethambutol (EMB) treated samples, and DNA fragments in **RuR3** and $H_2O_2$ treated samples. (b) Statistical analysis of DNA fragmentation upon different treatments.

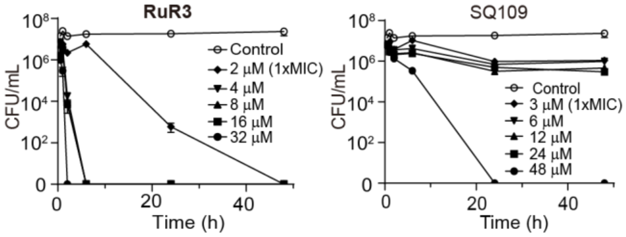


Supplementary Figure 11. Killing kinetics of dormant *M. smegmatis* in PBS by **RuR3** or SQ109. The initial cell density is ~2×$10^7$ CFU/mL.

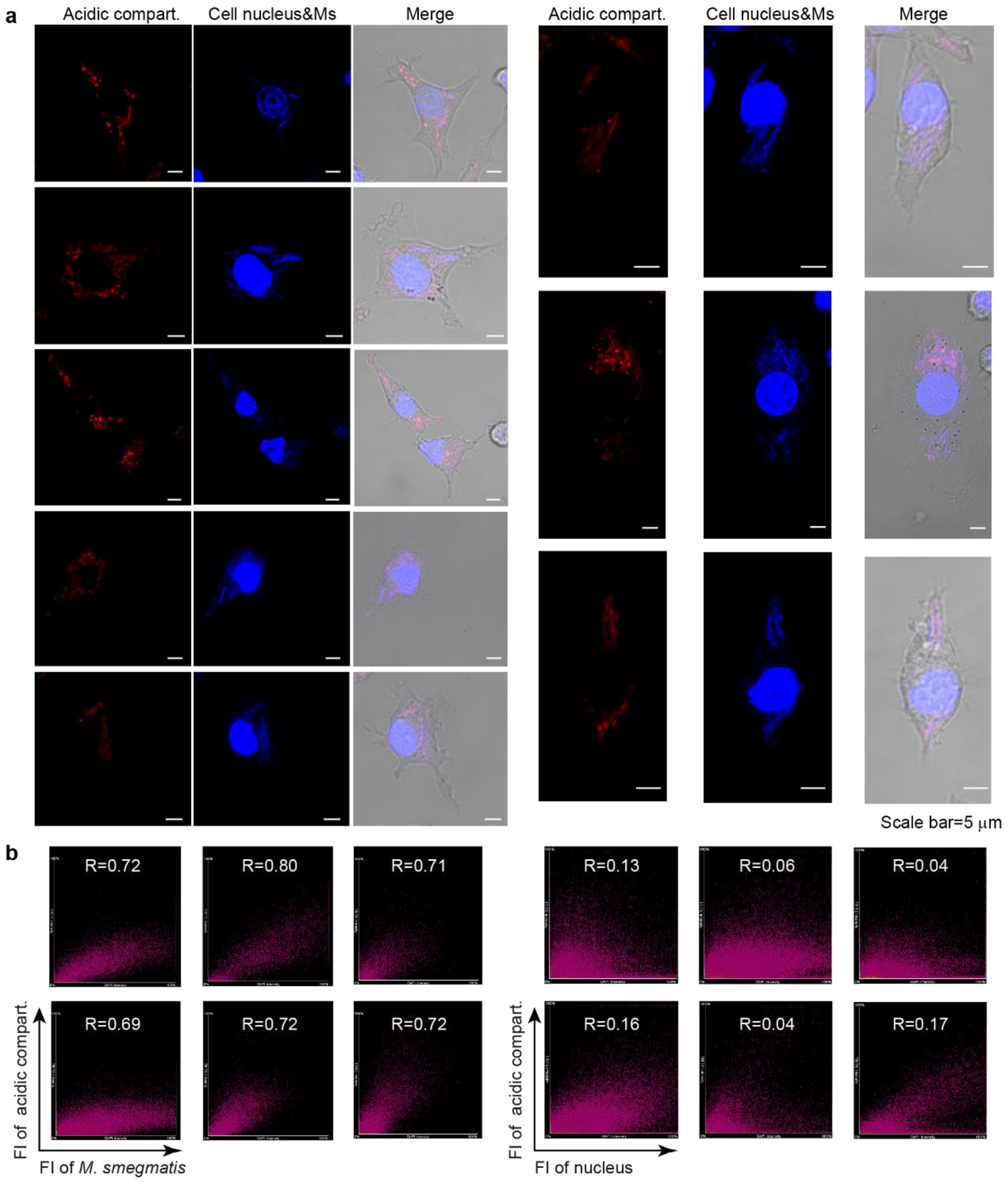


Supplementary Figure 12. (a) Additional confocal images of in *M. smegmatis* infected macrophages. The subcellular localization of *M. smegmatis* suggesting the residence of *M. smegmatis* in the acidic mycobacteria pathogen vacuoles (MCV, pH~6.2) (blue: Hoechst-stained cell nucleus and *M. smegmatis*; red: lysotracker-red to stain acidic compartment). (b) The 2D intensity histogram was plotted using acidic compartment fluorescence against *M. smegmatis* fluorescence, and acidic compartment fluorescence against nucleus fluorescence (as control), respectively.

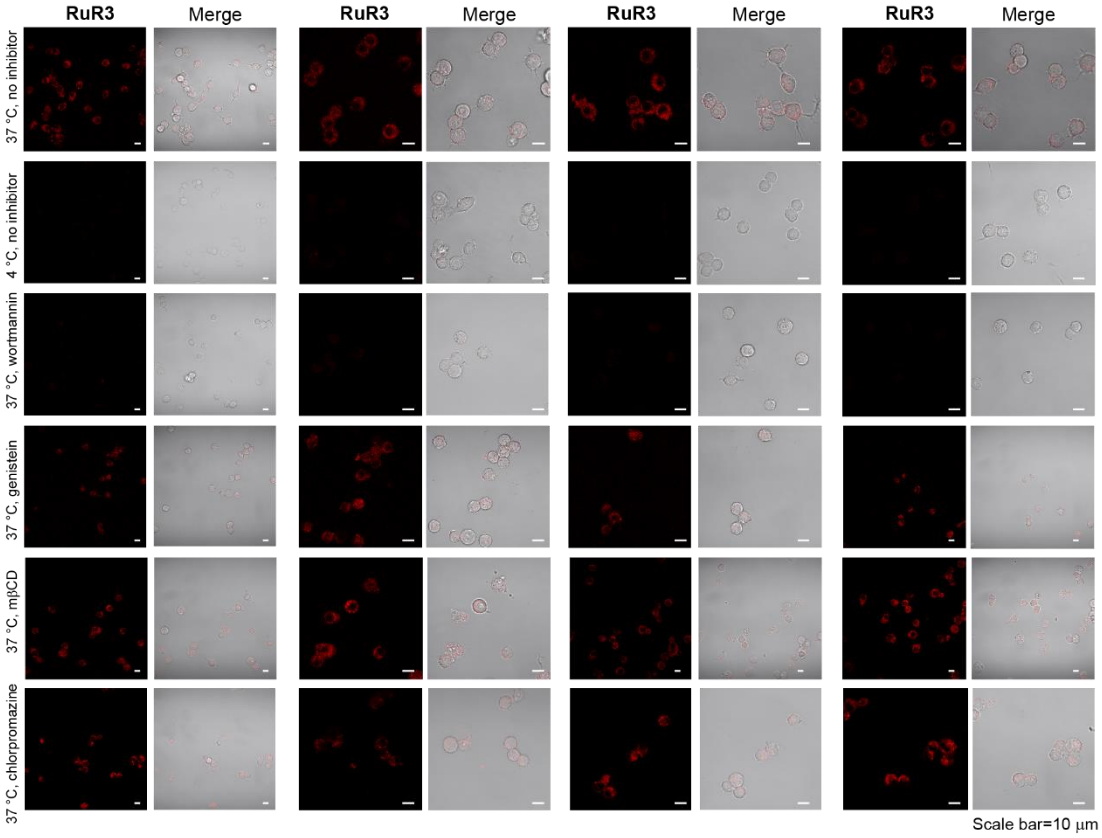


Supplementary Figure 13. Four sets of confocal images of cellular uptake levels of **RuR3** under different inhibitory conditions. The results suggest a macropinocytosis-mediated endocytic cell uptake mechanism (red: **RuR3** fluorescence). To quantify **RuR3** accumulation, the **RuR3** fluorescence for each cell in the image was calculated by ImageJ, and averaged for each treatment group. At least 30 cells were included for each group.

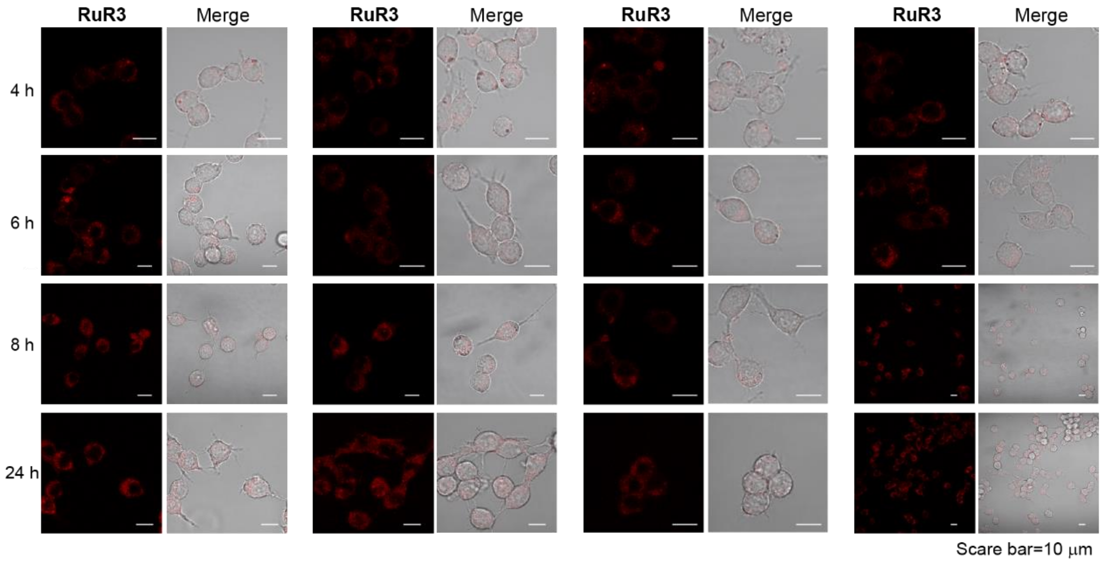


Supplementary Figure 14. Four sets of confocal images of RAW 264.7 after incubation with **RuR3** for different time point (red: **RuR3** fluorescence).

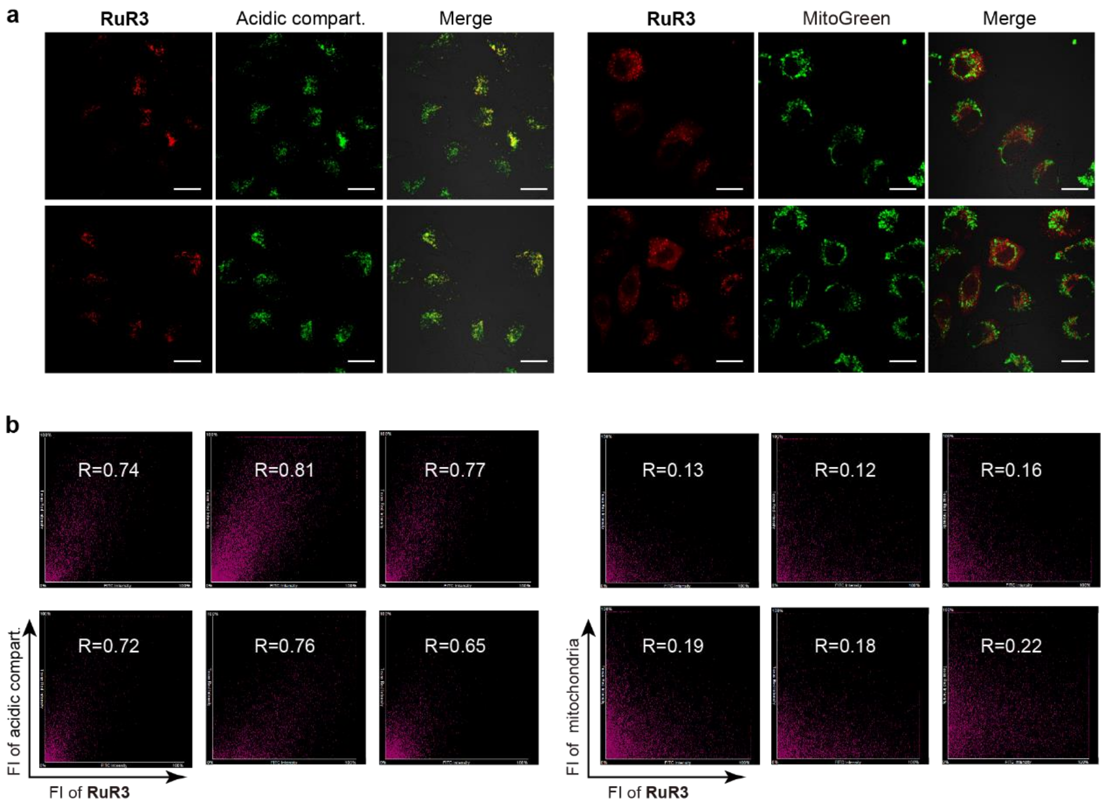


Supplementary Figure 15. (a) The additional confocal images of subcellular localization of **RuR3** suggesting the residence of **RuR3** in the acidic endosomes or lysosomes (red: **RuR3** fluorescence; green: lysotracker-green to stain acidic compartment). (b) The 2D intensity histogram was plotted using acidic endosomes or lysosomes fluorescence against **RuR3** fluorescence, and mitochondria fluorescence against **RuR3** fluorescence, respectively.

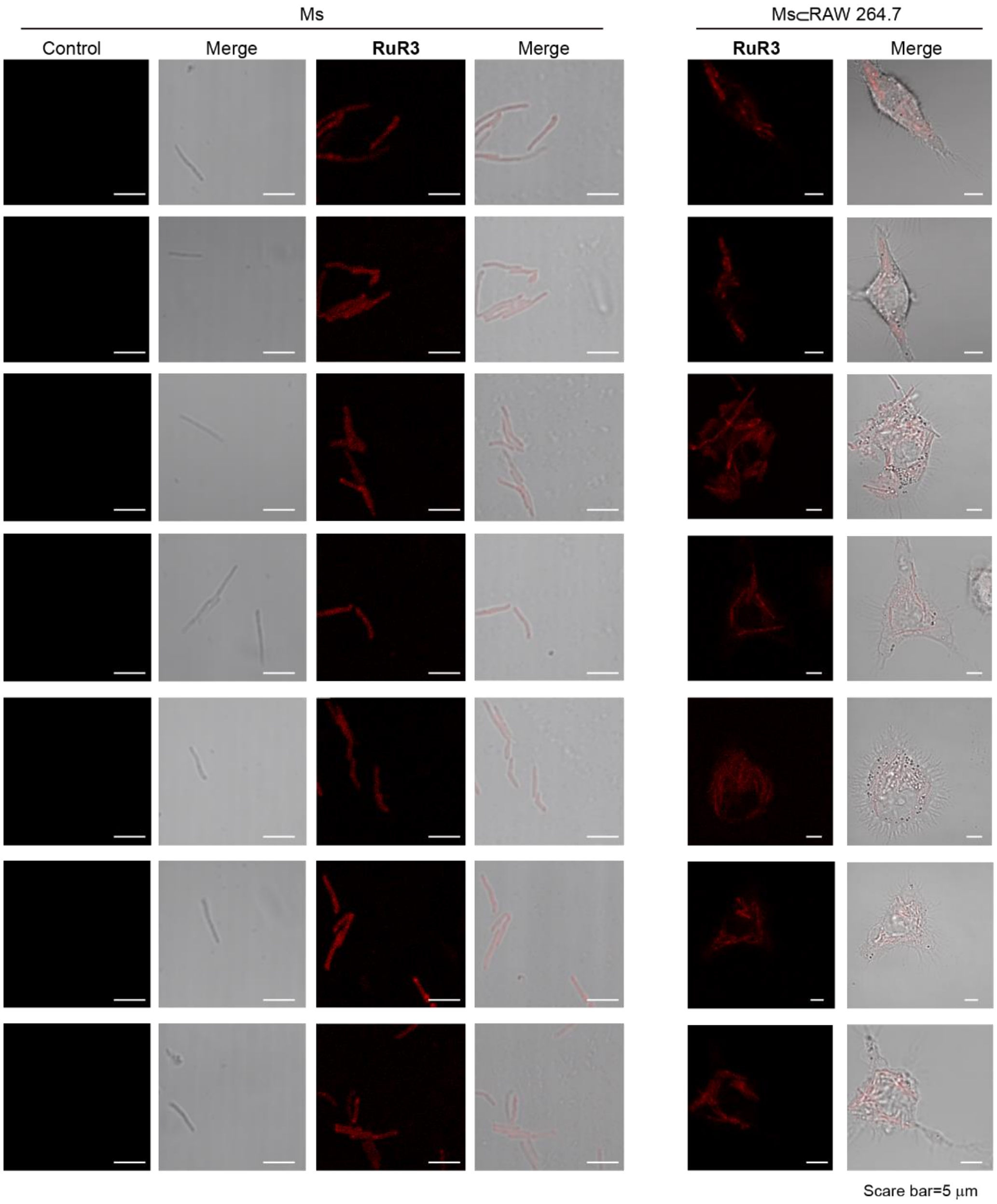


Supplementary Figure 16. The additional confocal images of subcellular colocalization of **RuR3** and *M. smegmatis* (red: **RuR3** fluorescence).

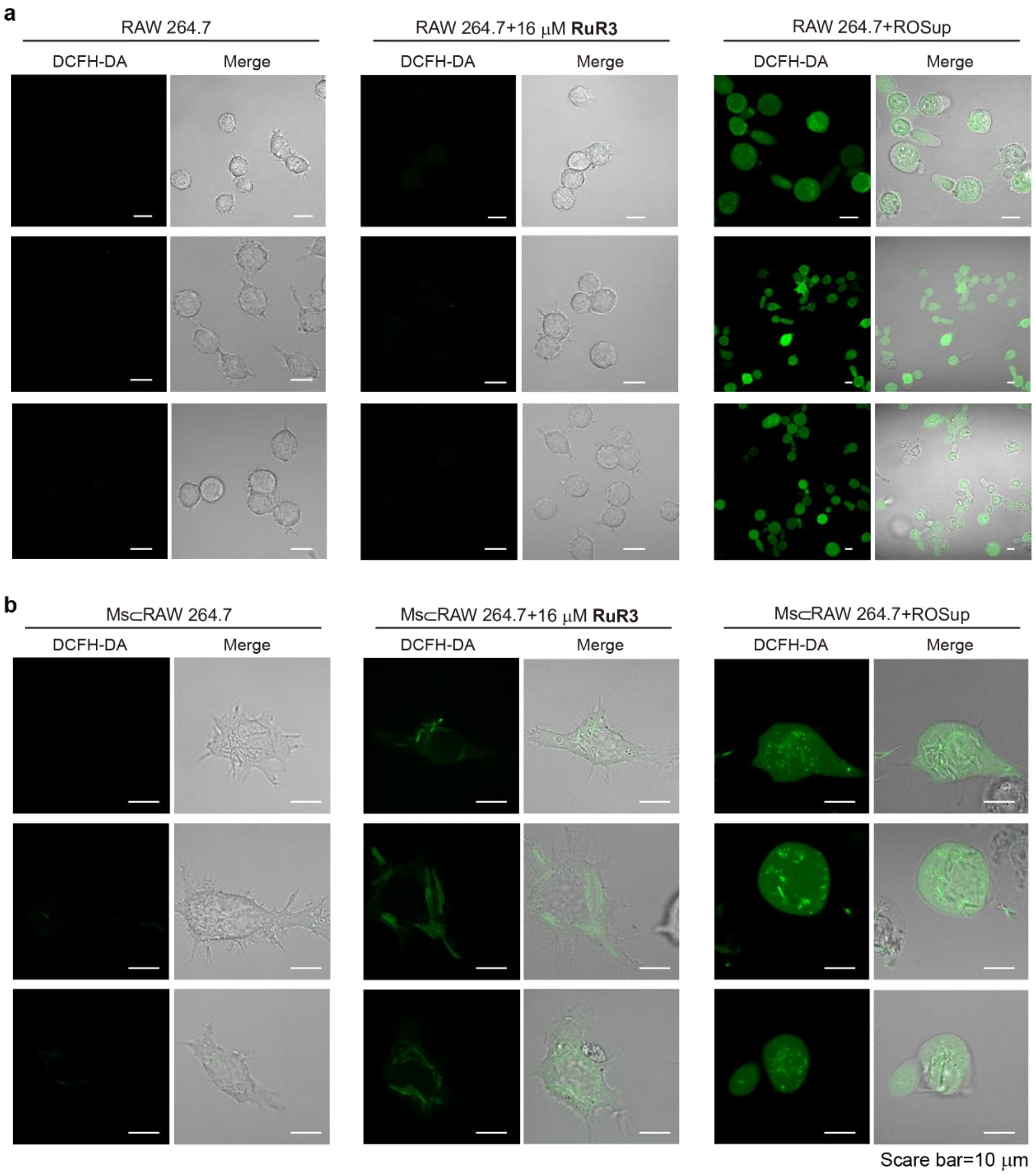


Supplementary Figure 17. (a) The additional confocal images of **RuR3**-induced ROS in uninfected RAW 264.7. (b) The additional confocal images of **RuR3**-induced ROS in *M. smegmatis* infected cells (Ms⊂RAW 264.7, green: DCFH-DA probed ROS).

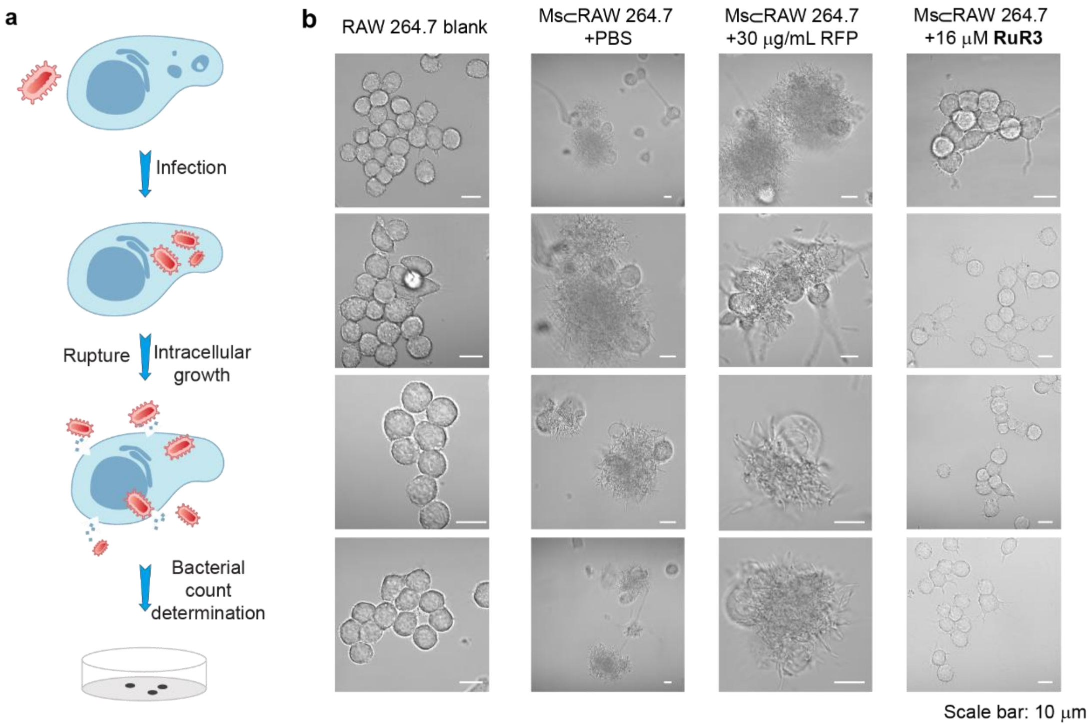


Supplementary Figure 18. (a) Cartoon illustration of the intracellular bacteria eradication experiment. (b) The additional brightfield images of Ms⊂RAW 264.7 cells after a 24 h treatment with PBS, RFP or **RuR3**.

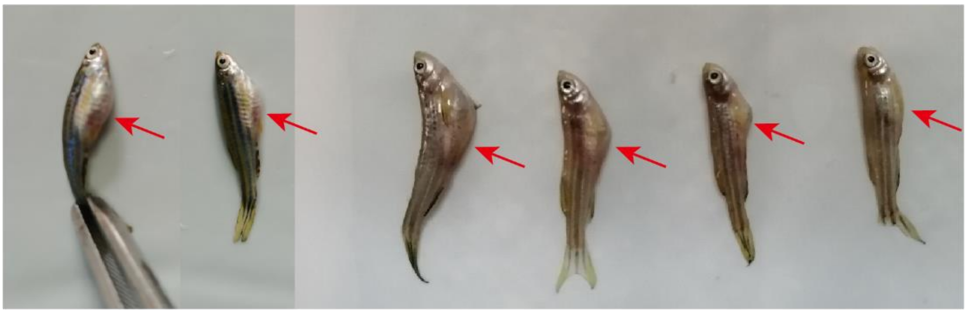

Supplementary Figure 19. The additional images of dead zebrafish with characteristic distended abdomen from *M. fortuitum* infection.

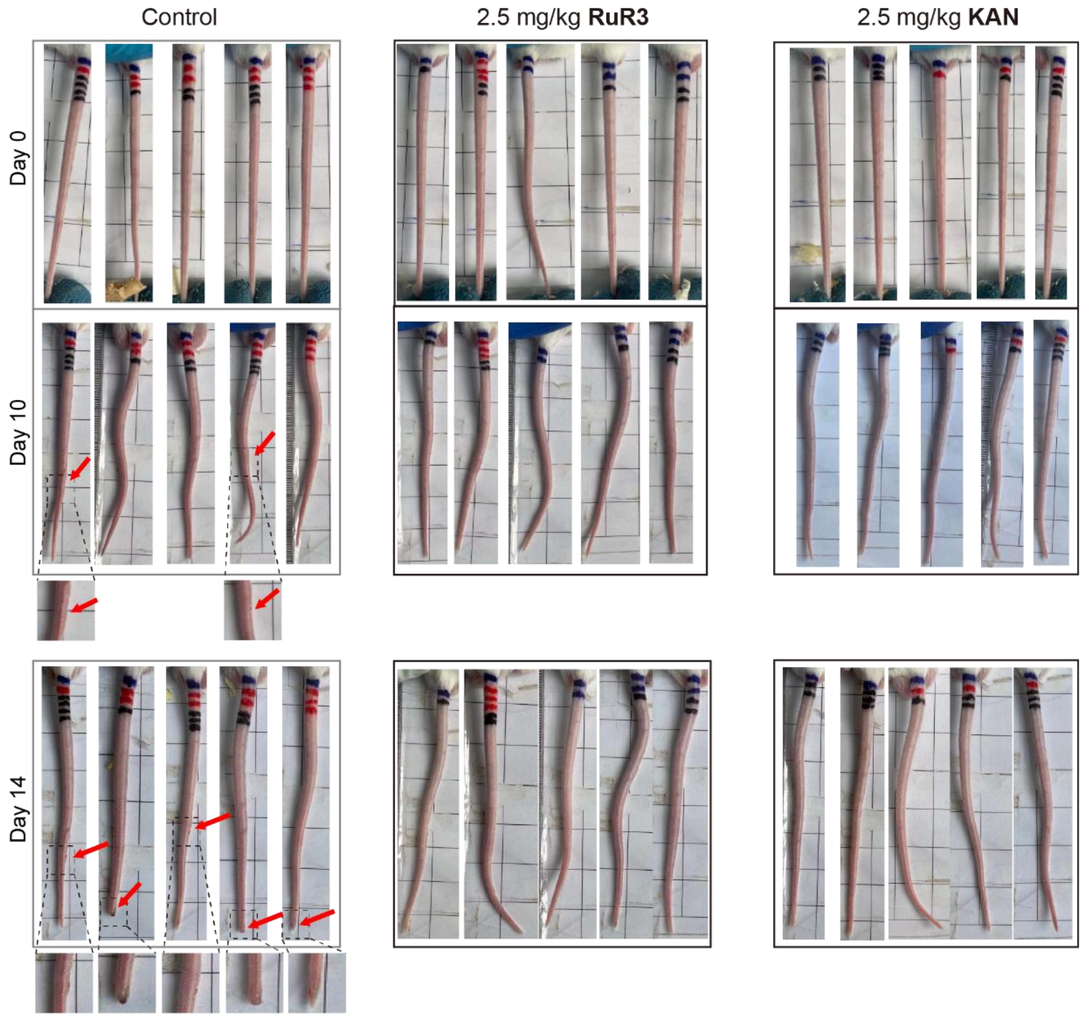


Supplementary Figure 20. The additional images of mice tails taken at 1, 10- and 14-days post infection, respectively.

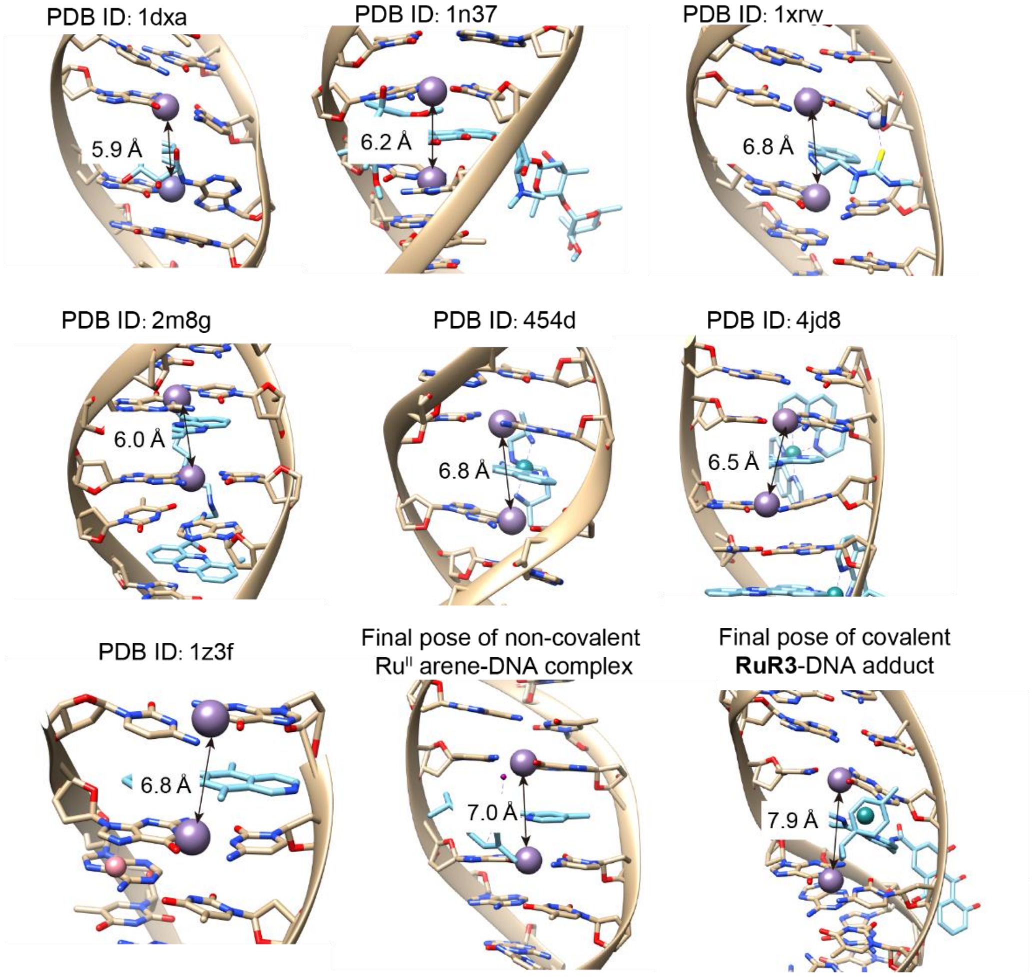


Supplementary Figure 21. Average helix rises per base pair for reported structure with intercalators, non-covalent $Ru^{II}$ arene-DNA complex and covalent **RuR3**-DNA adduct.

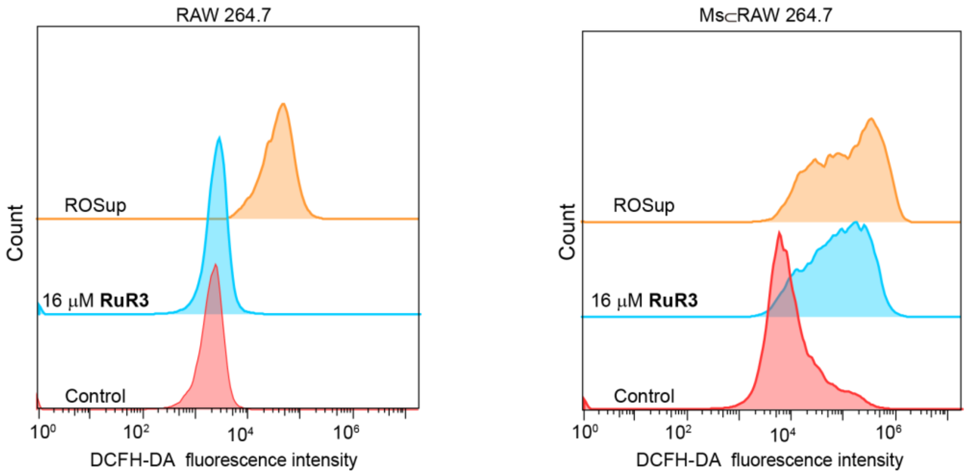


Supplementary Figure 22. Flow cytometric analysis of ROS accumulation of **RuR3** in RAW 264.7 or Ms⊂RAW 264.7.

ROS generation level is calculated as following (GEO=The FL geometric mean of ROS generation):

$$\text{Relative ROS level upon compound treatment} = \frac{\text{GEO of compound of interest}}{\text{GEO of no treatment group}}$$

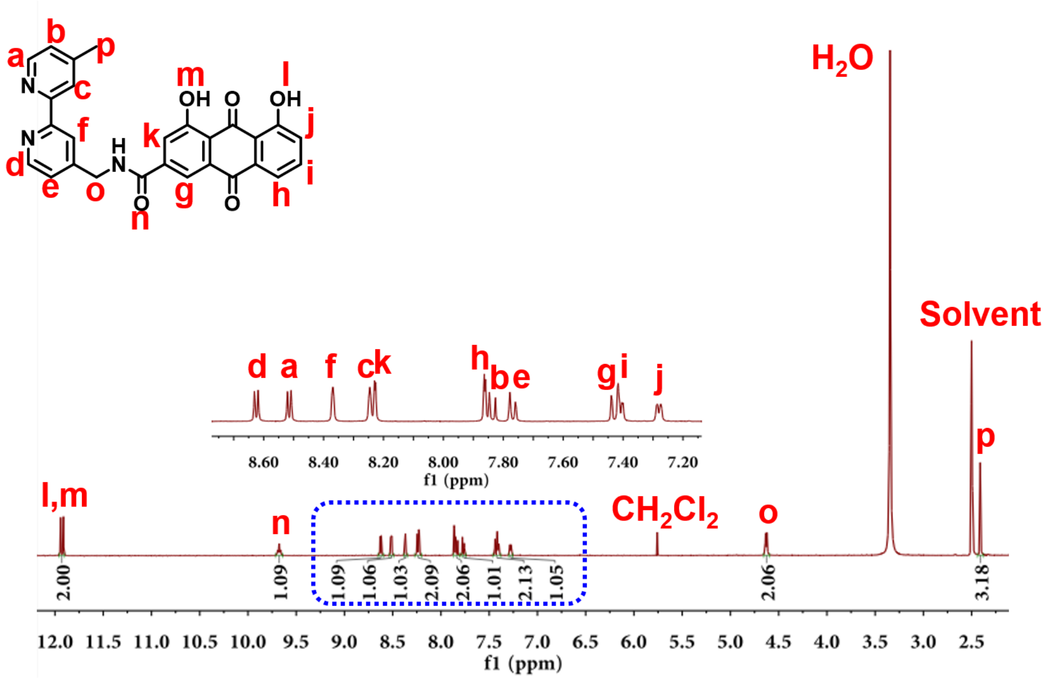


Supplementary Figure 23. [1]H NMR spectrum (400 MHz, $d_6$-DMSO) of L1.

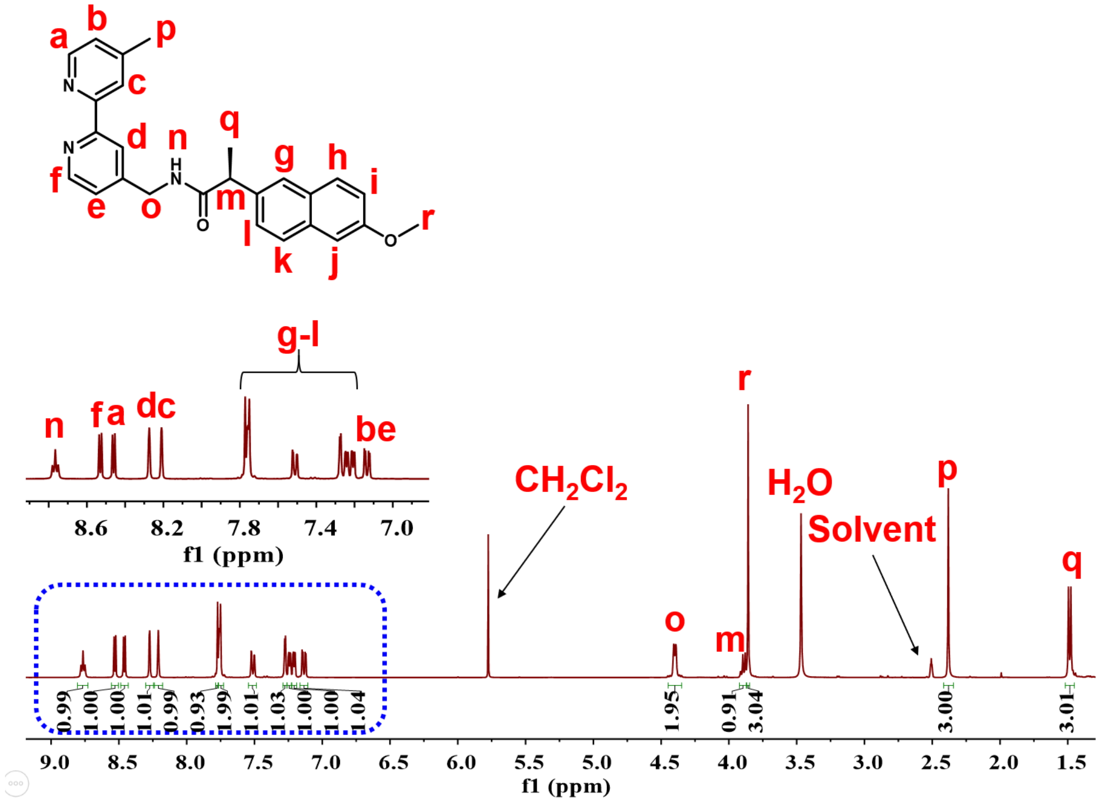


Supplementary Figure 24. [1]H NMR spectrum (400 MHz, $d_6$-DMSO) of L2.

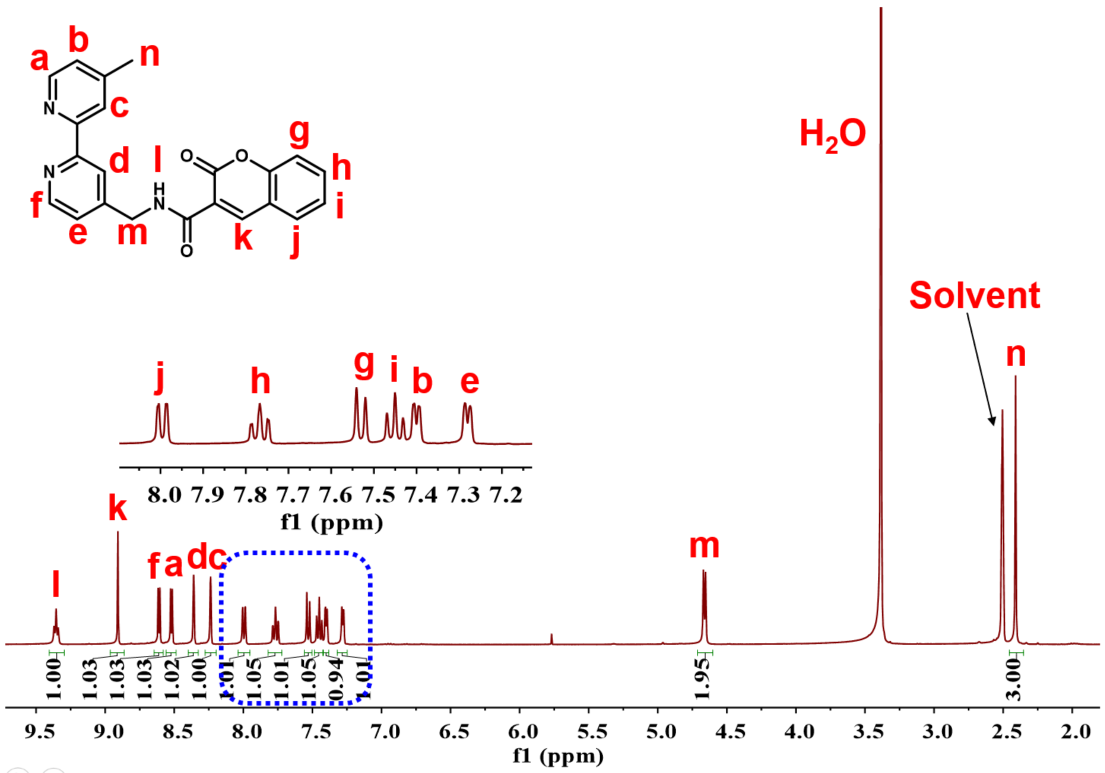


Supplementary Figure 25. $^1$H NMR spectrum (400 MHz, $d_6$-DMSO) of L3.

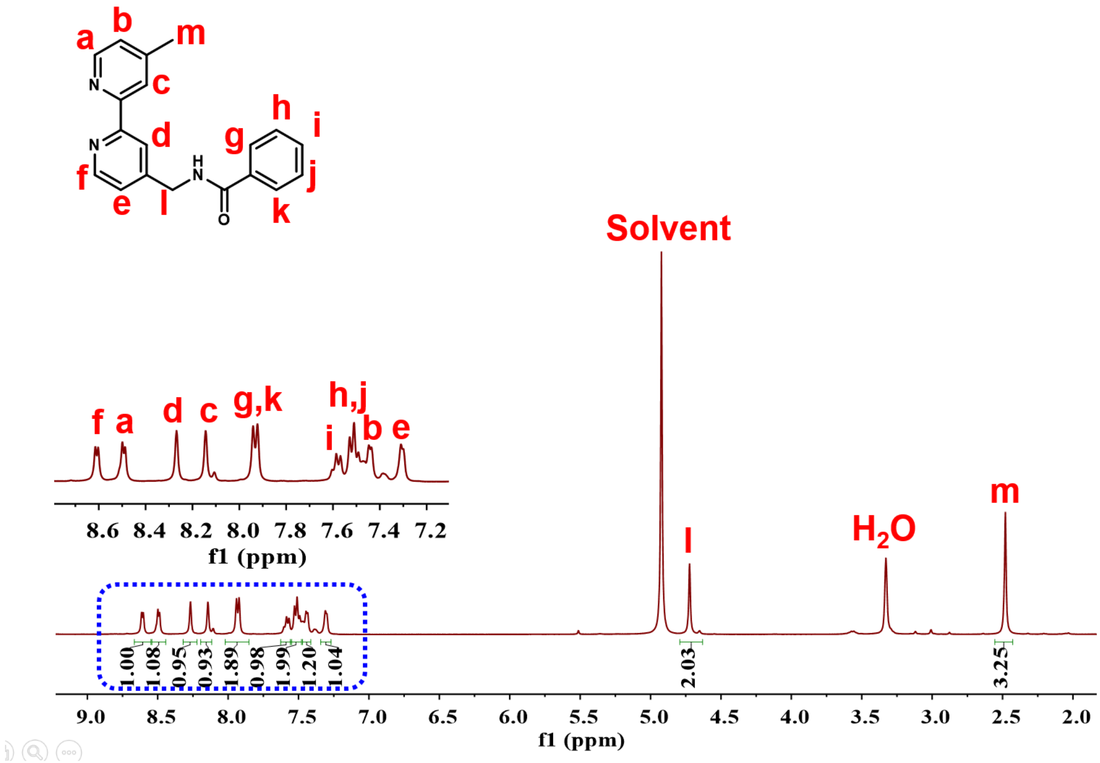


Supplementary Figure 26. $^1$H NMR spectrum (400 MHz, $d_4$-Methanol) of L4.

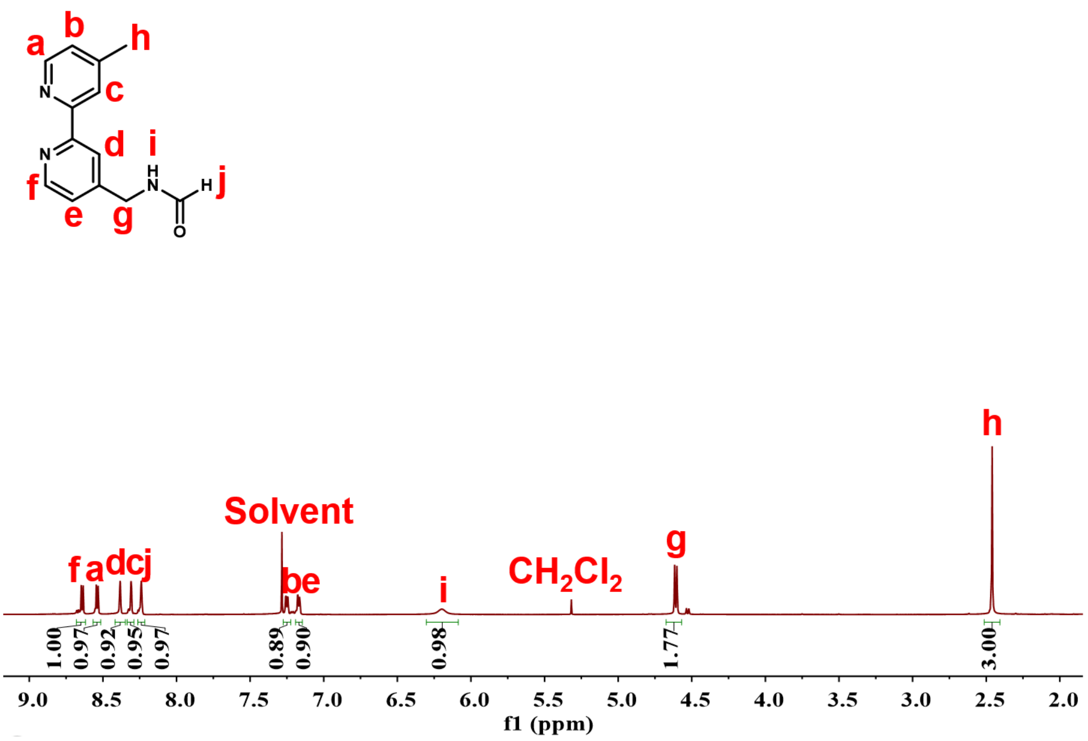


Supplementary Figure 27. ${}^{1}H$ NMR spectrum (400 MHz, *d*-Chloroform) of L5.

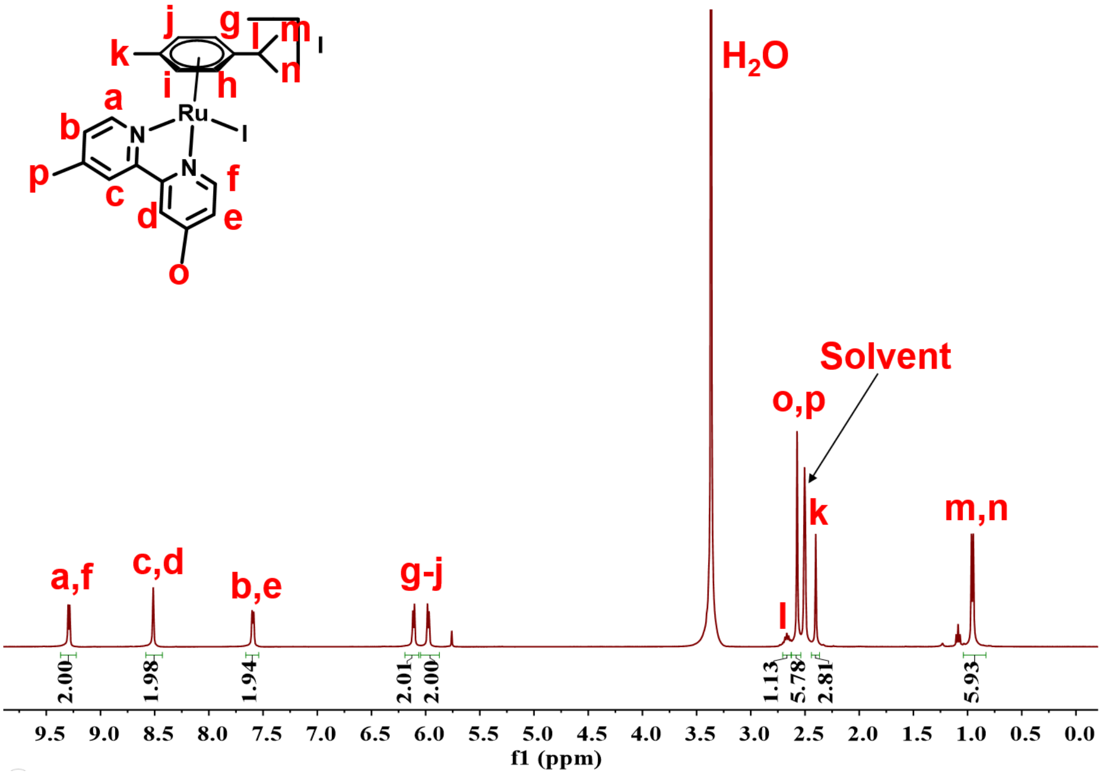


Supplementary Figure 28. ${}^{1}H$ NMR spectrum (400 MHz, *d₆*-DMSO) of $Ru^{II}$-arene.

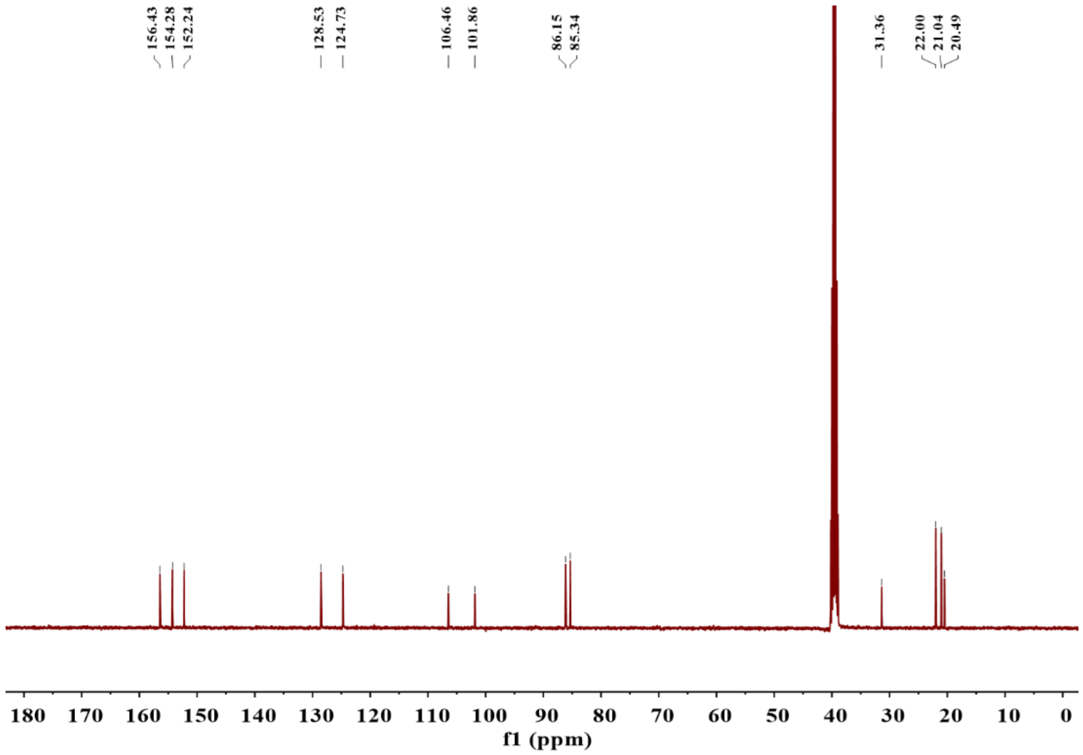


Supplementary Figure 29. [13]C NMR spectrum (400 MHz, $d_6$-DMSO) of Ru$^{II}$-arene.

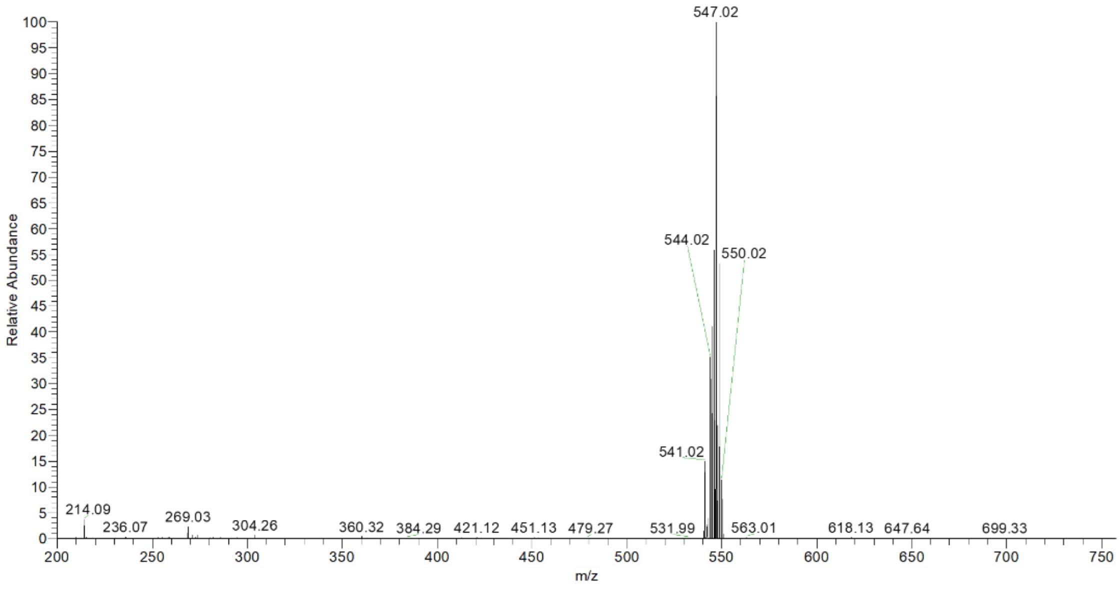


Supplementary Figure 30. ESI-MS spectrum of Ru$^{II}$-arene in $CH_3OH$. (calcd for [Ru$^{II}$-arene-I$^-$]$^+$ m/z = 546.43, found m/z = 547.02.)

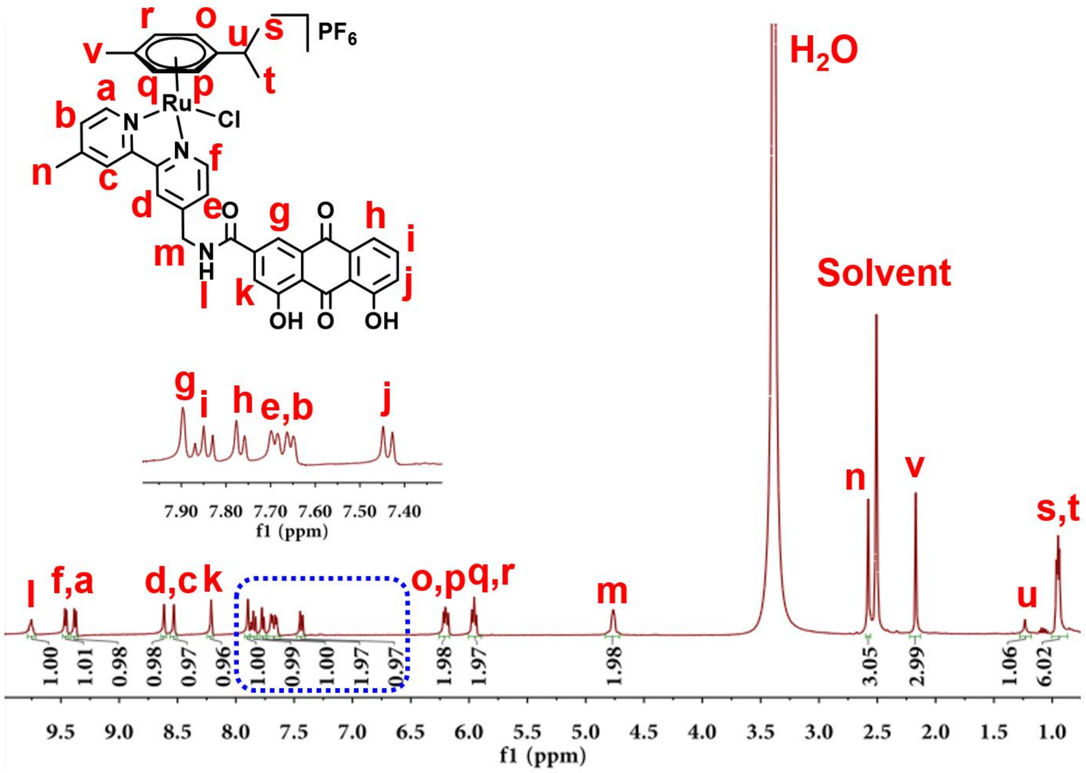

Supplementary Figure 31. $^{1}$H NMR spectrum (400 MHz, $d_6$-DMSO) of **RuR1**.

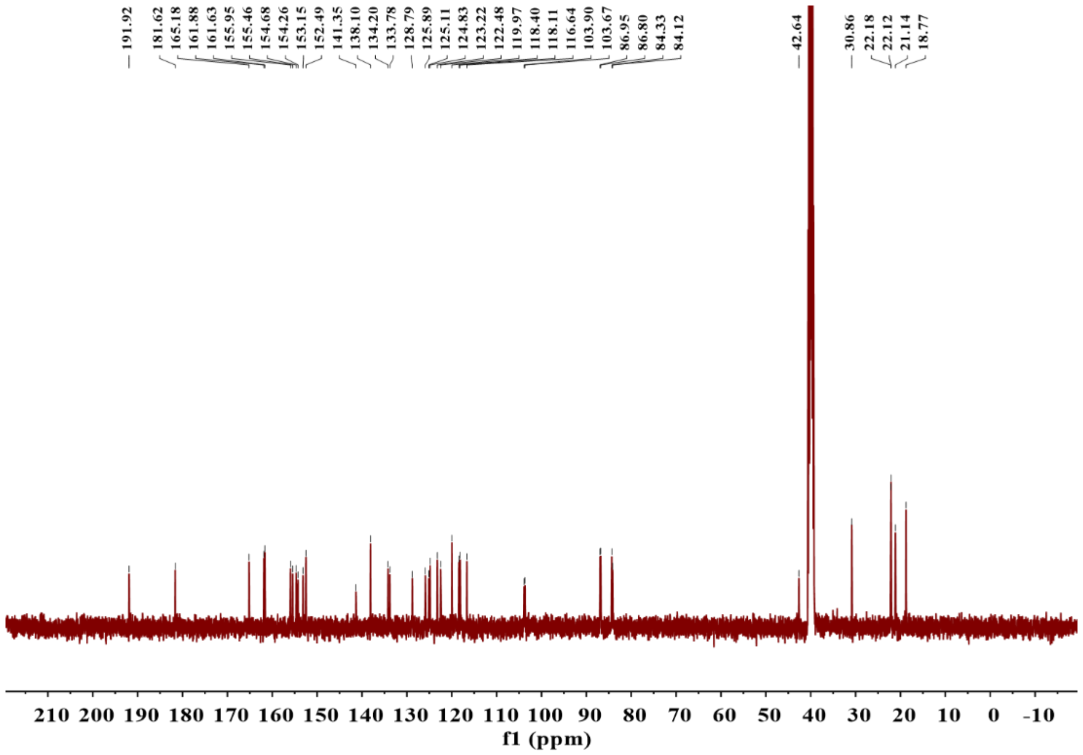

Supplementary Figure 32. $^{13}$C NMR spectrum (400 MHz, $d_6$-DMSO) of **RuR1**.

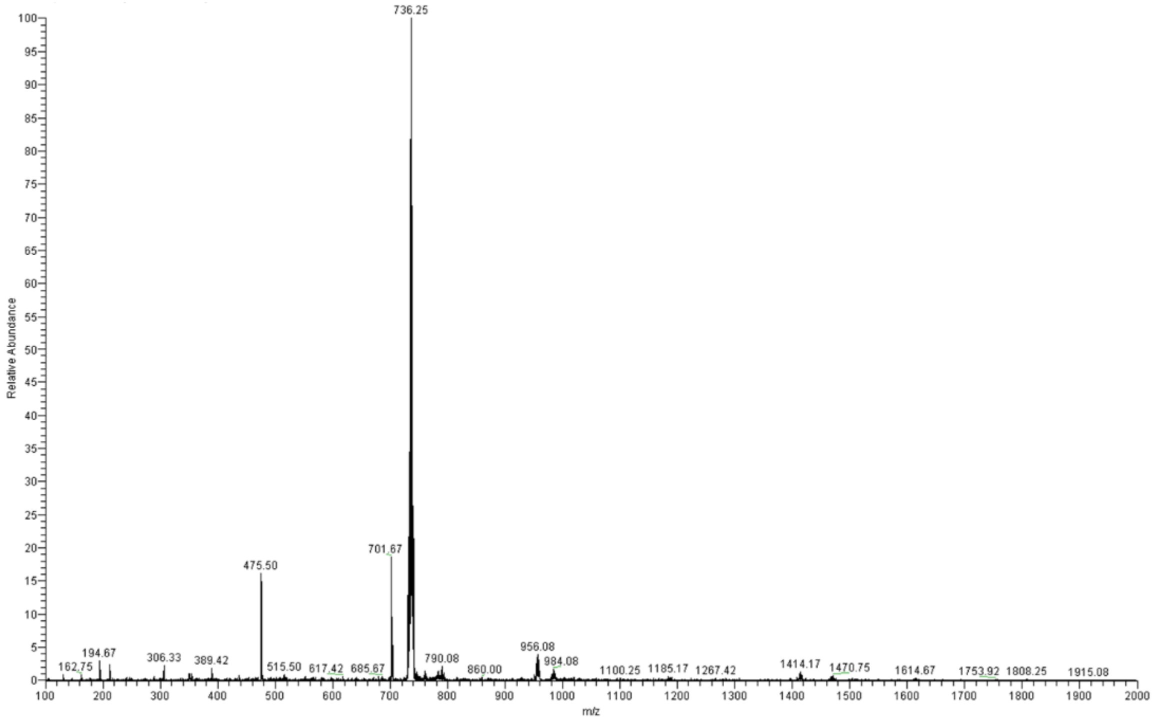


Supplementary Figure 33. ESI-MS spectrum of **RuR1** in $CH_3OH$. (calcd for [**RuR1**-$PF_6^-$]$^+$ m/z = 736.21, found m/z = 736.25.)

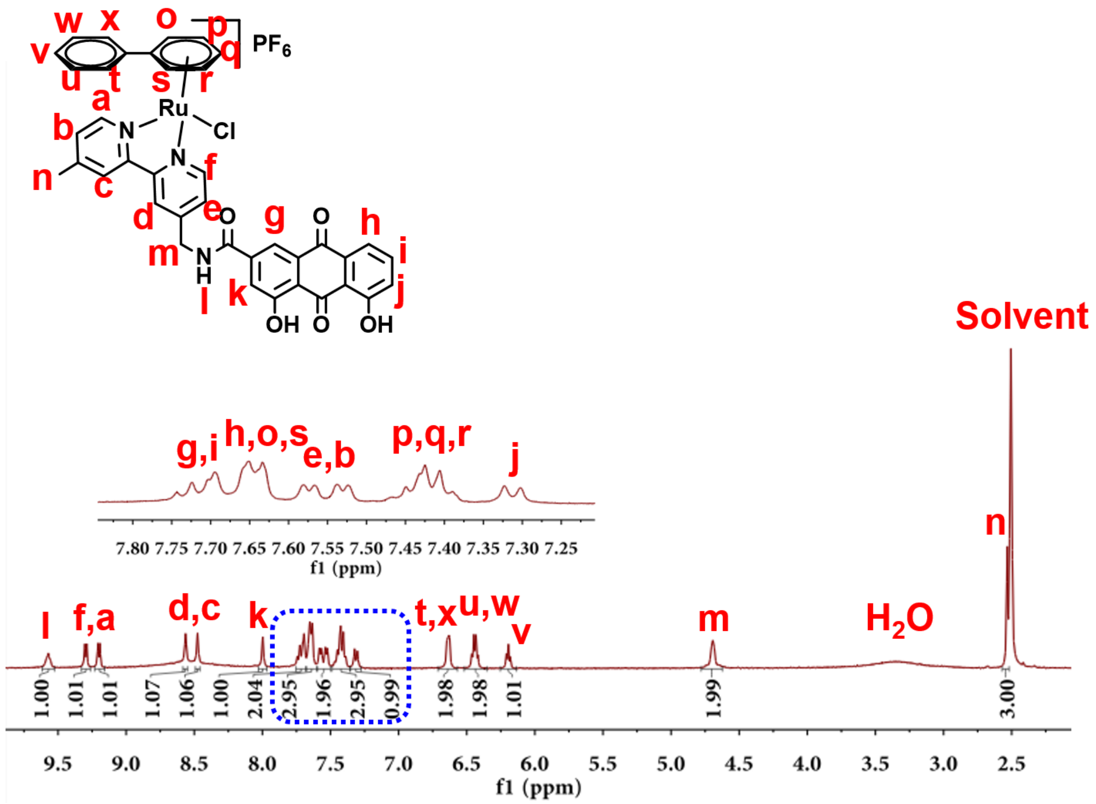


Supplementary Figure 34. $^1$H NMR spectrum (400 MHz, $d_6$-DMSO) of **RuR2**.

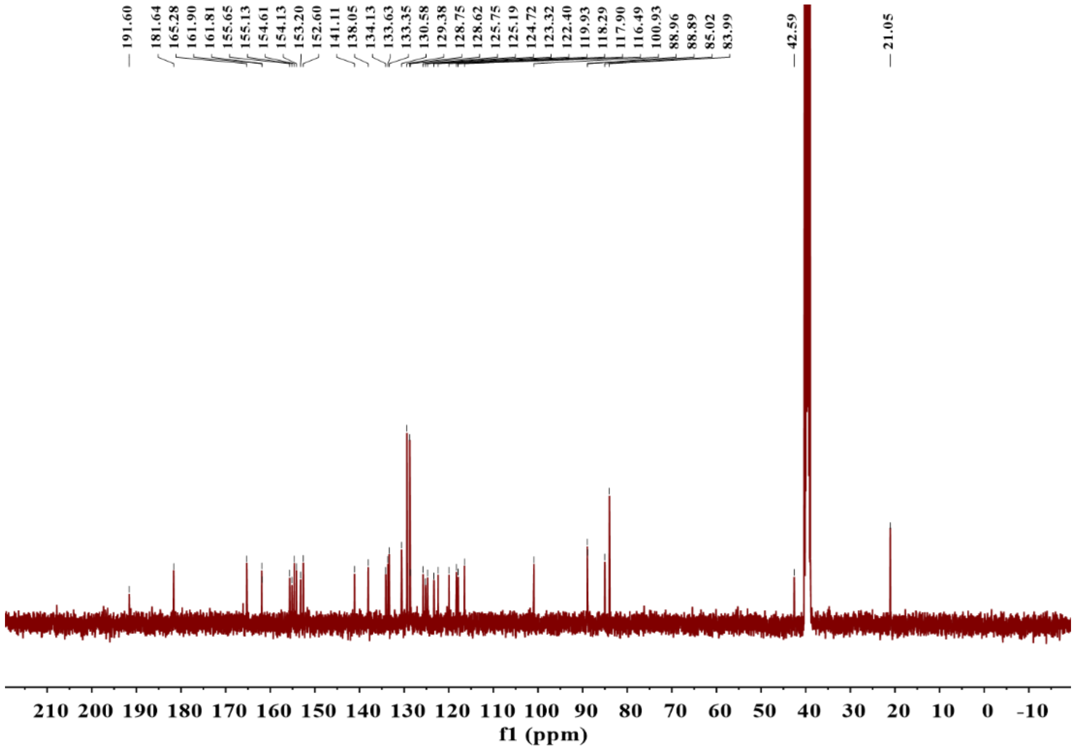


Supplementary Figure 35. $^{13}C$ NMR spectrum (400 MHz, $d_6$-DMSO) of **RuR2**.

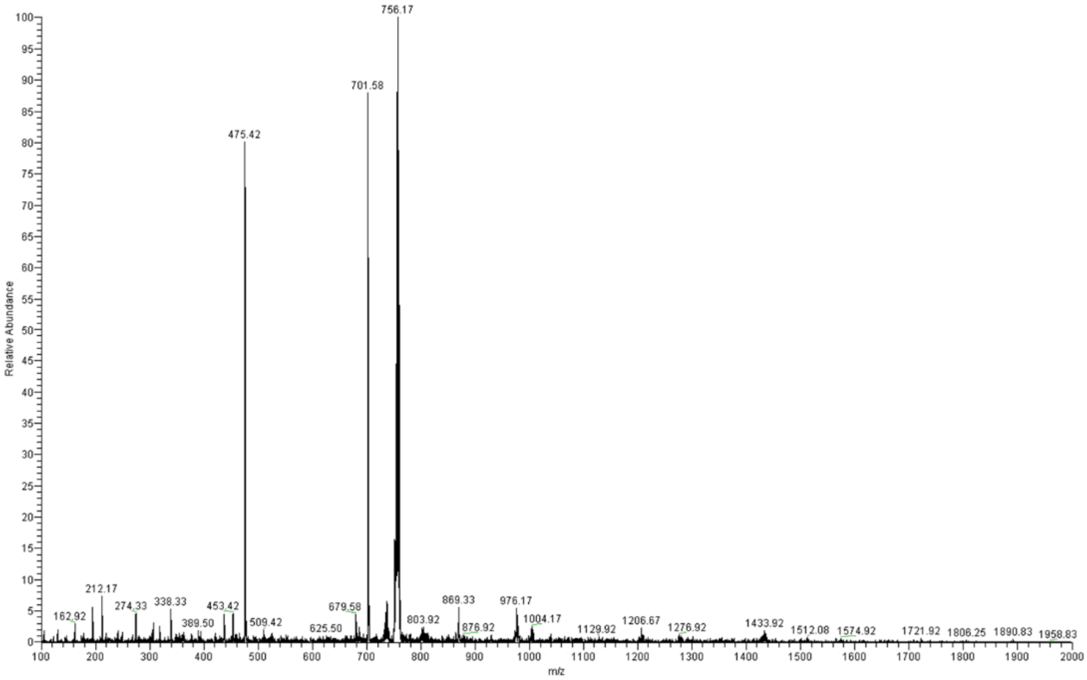


Supplementary Figure 36. ESI-MS spectrum of **RuR2** in $CH_3OH$. (calcd for [**RuR2**-$PF_6^-$]$^+$ m/z = 756.20, found m/z = 756.17.)

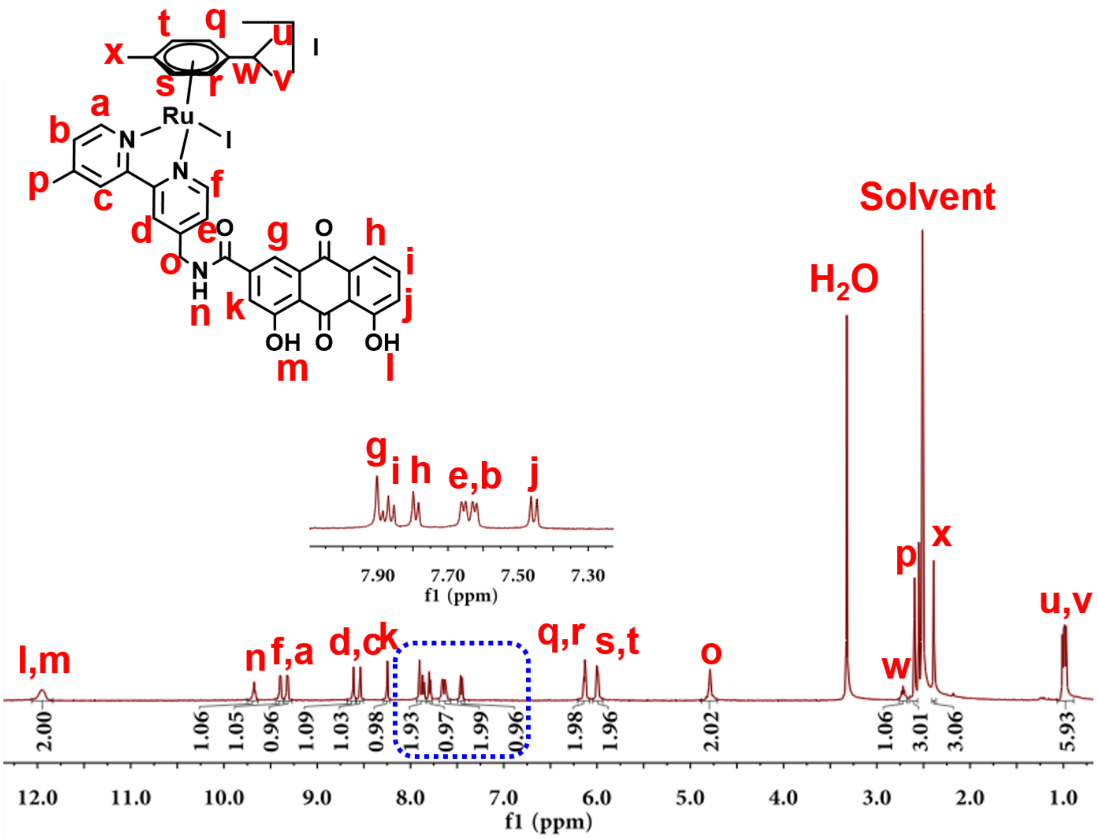


Supplementary Figure 37. $^{1}H$ NMR spectrum (400 MHz, $d_6$-DMSO) of **RuR3**.

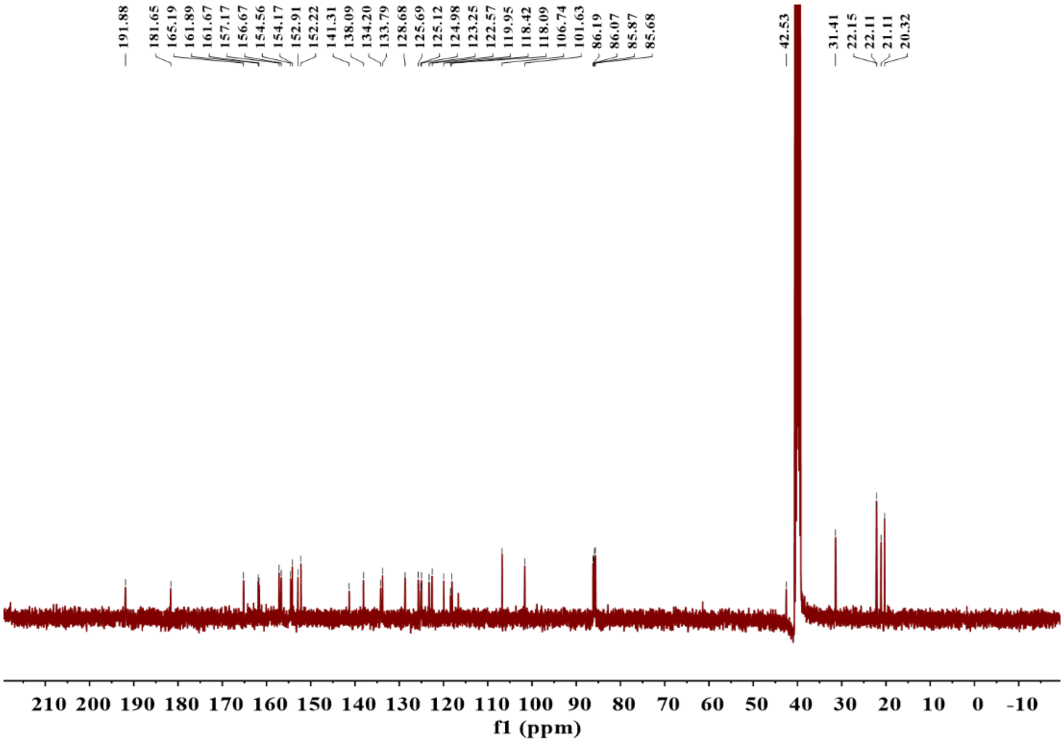


Supplementary Figure 38. $^{13}C$ NMR spectrum (400 MHz, $d_6$-DMSO) of **RuR3**.

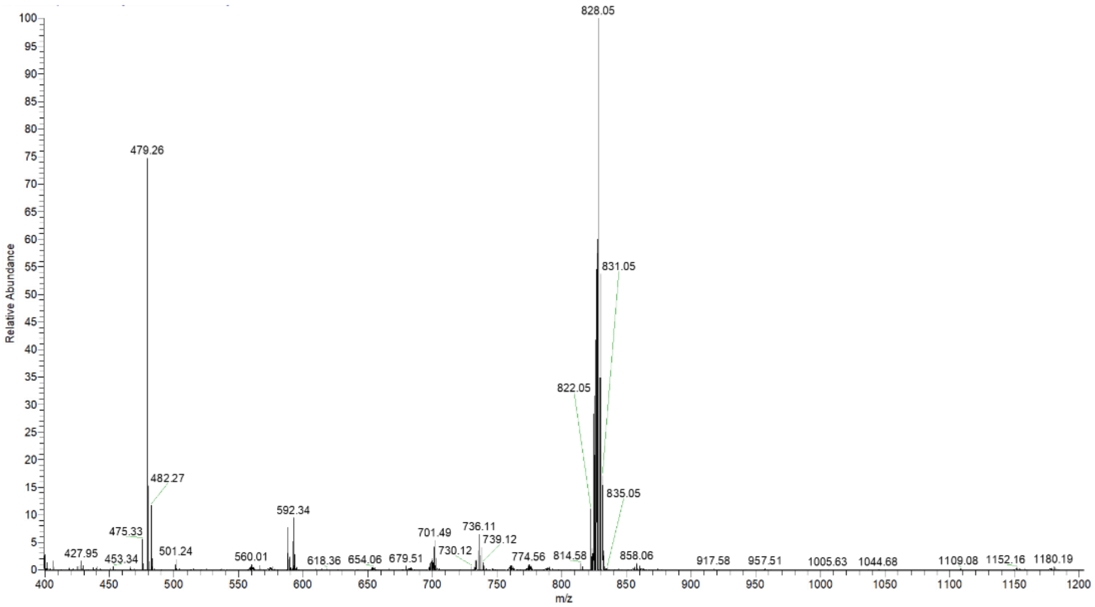

Supplementary Figure 39. ESI-MS spectrum of **RuR3** in $CH_3OH$. (calcd for [**RuR3**-$I^-$]$^+$ m/z = 827.65, found m/z = 828.05.)

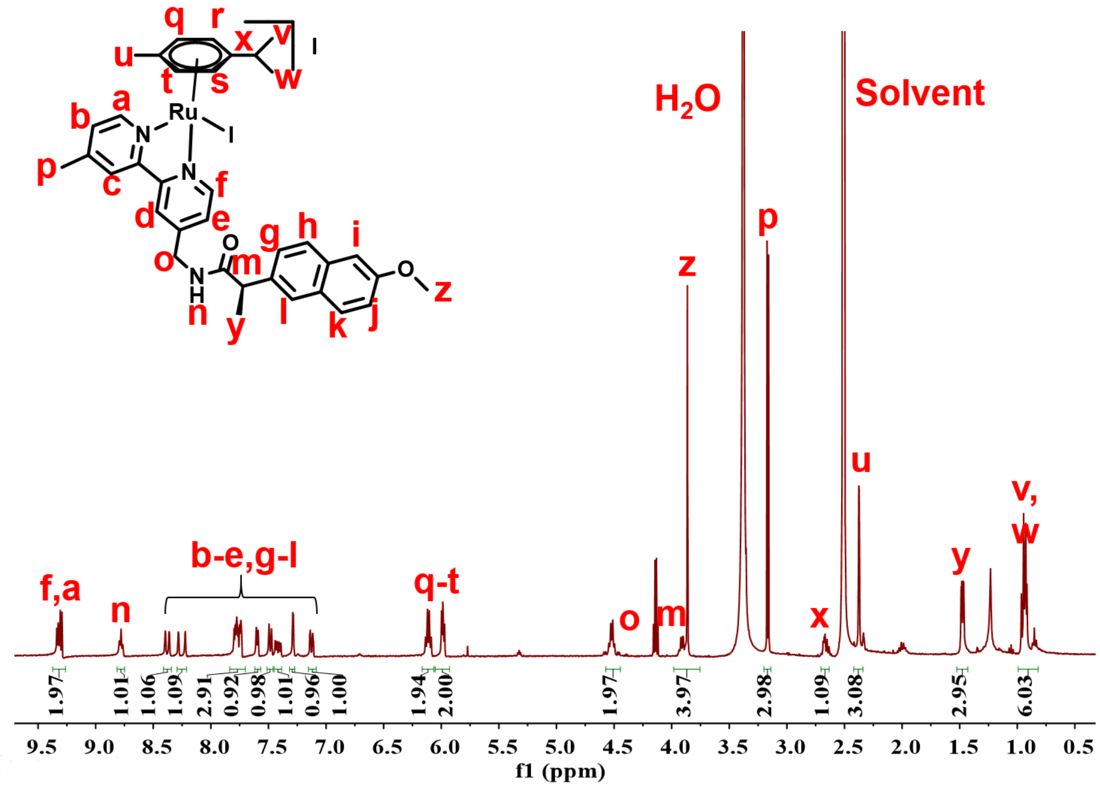

Supplementary Figure 40. $^1H$ NMR spectrum (400 MHz, $d_6$-DMSO) of **RuN**.

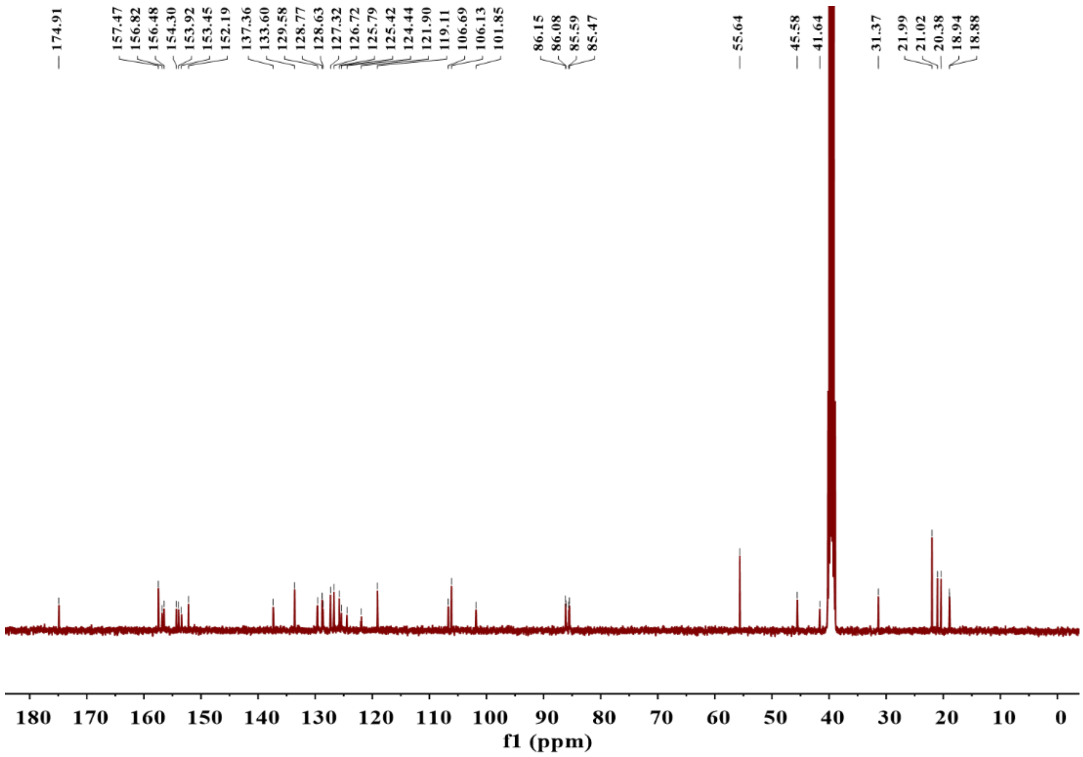


Supplementary Figure 41. $^{13}$C NMR spectrum (400 MHz, $d_6$-DMSO) of **RuN**.

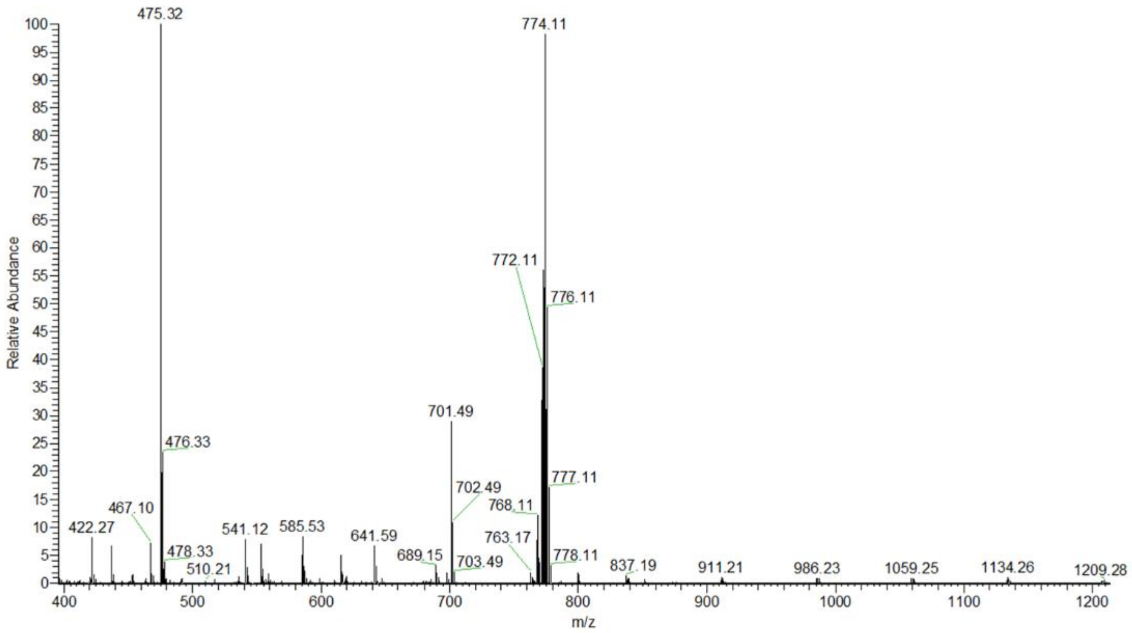


Supplementary Figure 42. ESI-MS spectrum of **RuN** in $CH_3OH$. (calcd for [**RuN**-I$^-$]$^+$ m/z = 773.61, found m/z = 774.11.)

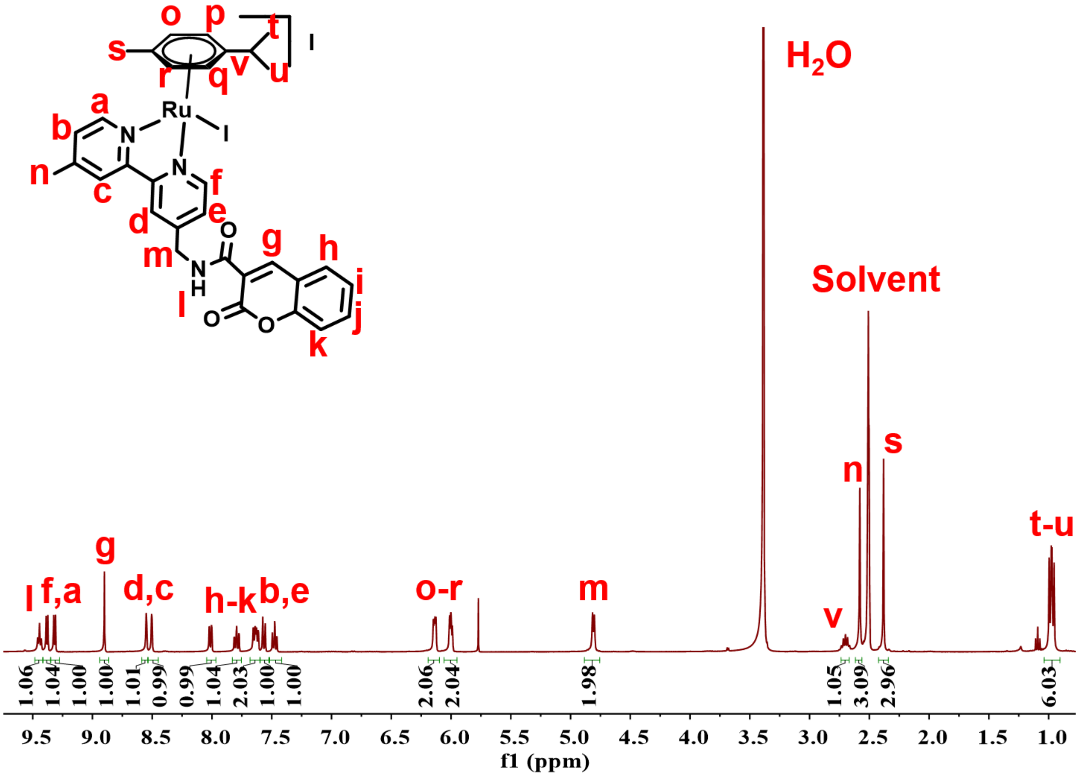


Supplementary Figure 43. $^{1}H$ NMR spectrum (400 MHz, $d_6$-DMSO) of **RuC**.

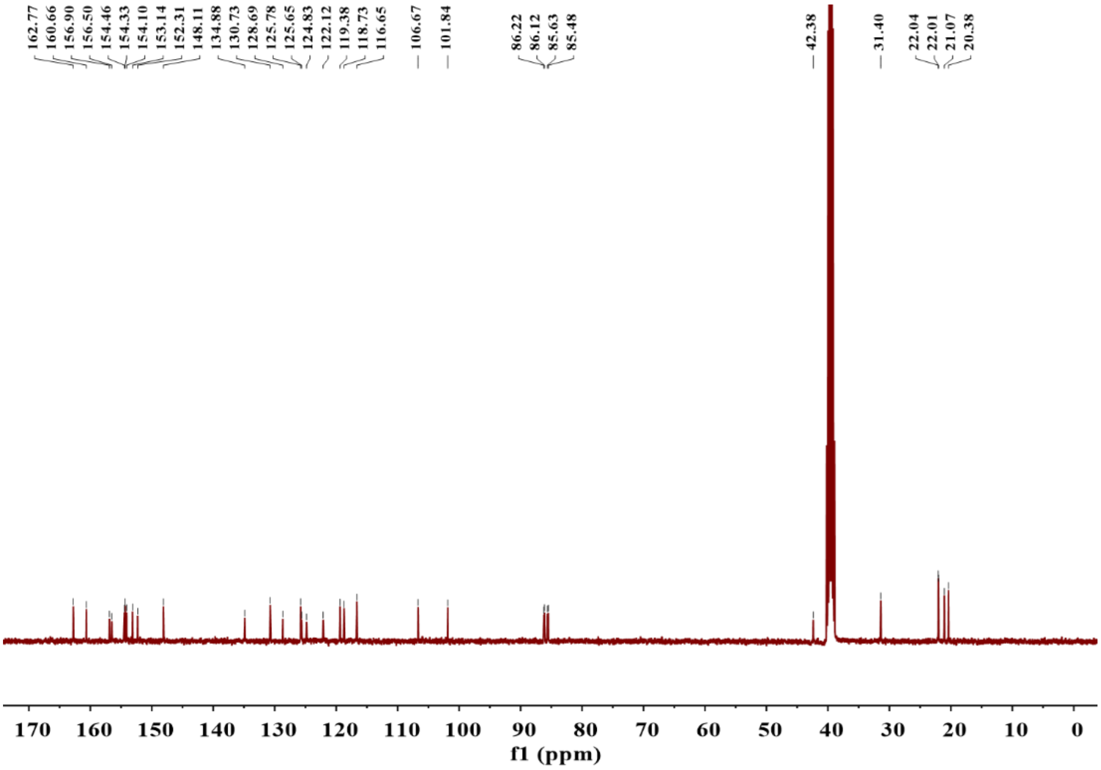


Supplementary Figure 44. $^{13}C$ NMR spectrum (400 MHz, $d_6$-DMSO) of **RuC**.

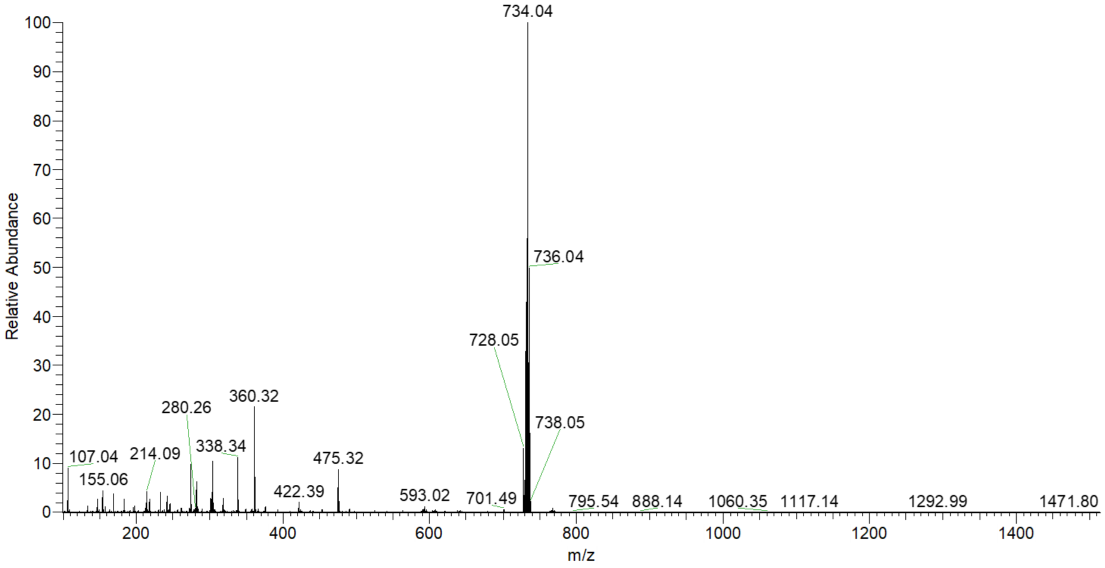


Supplementary Figure 45. ESI-MS spectrum of **RuC** in $CH_3OH$. (calcd for [**RuC**-$I^-$]$^+$ m/z = 733.5, found m/z = 734.04.)

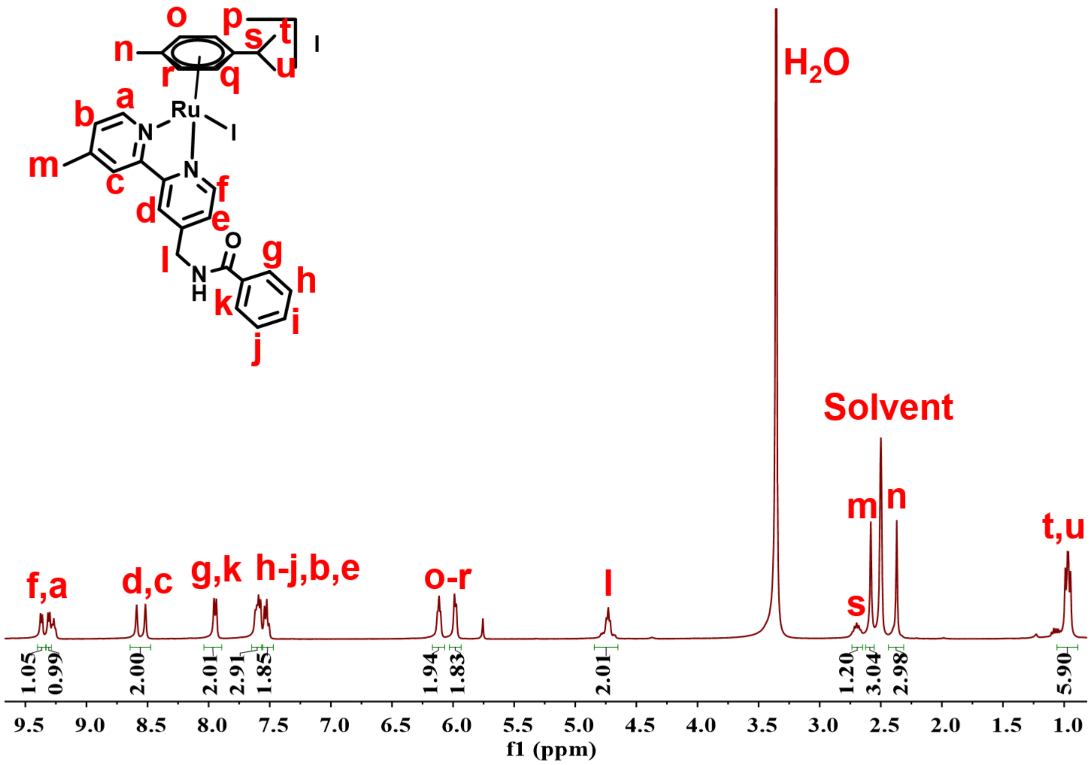


Supplementary Figure 46. $^1$H NMR spectrum (400 MHz, $d_6$-DMSO) of **RuB**.

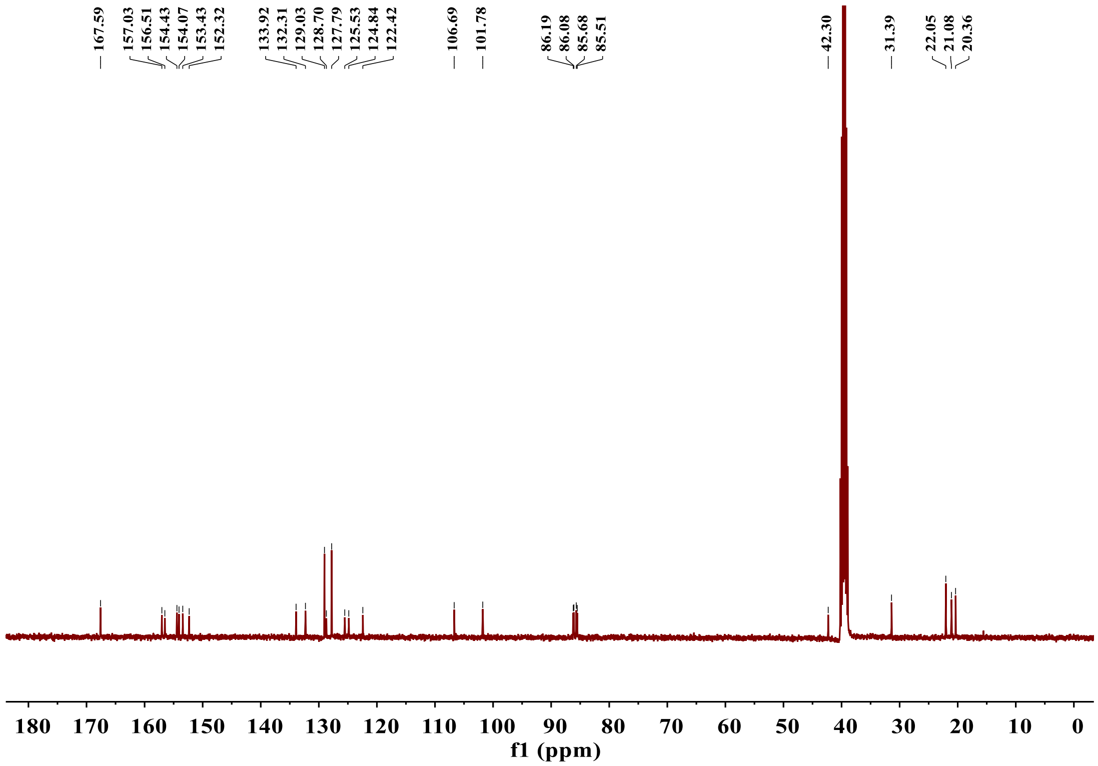


Supplementary Figure 47. $^{13}$C NMR spectrum (400 MHz, $d_6$-DMSO) of **RuB**.

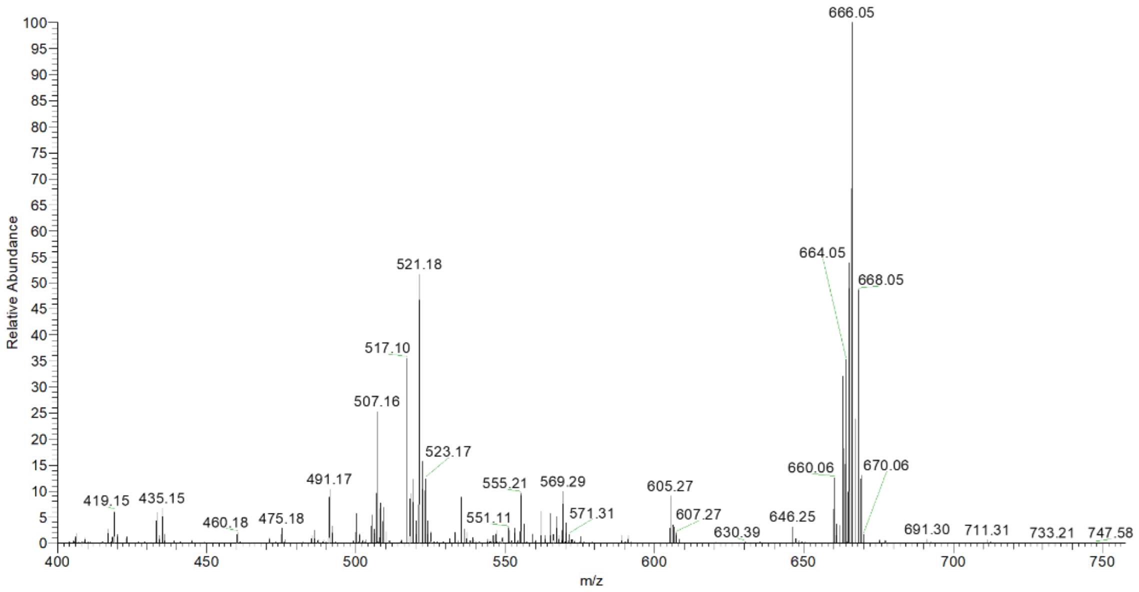


Supplementary Figure 48. ESI-MS spectrum of **RuB** in $CH_3OH$. (calcd for [**RuB**-$I^-$]$^+$ m/z = 665.47, found m/z = 666.05.)

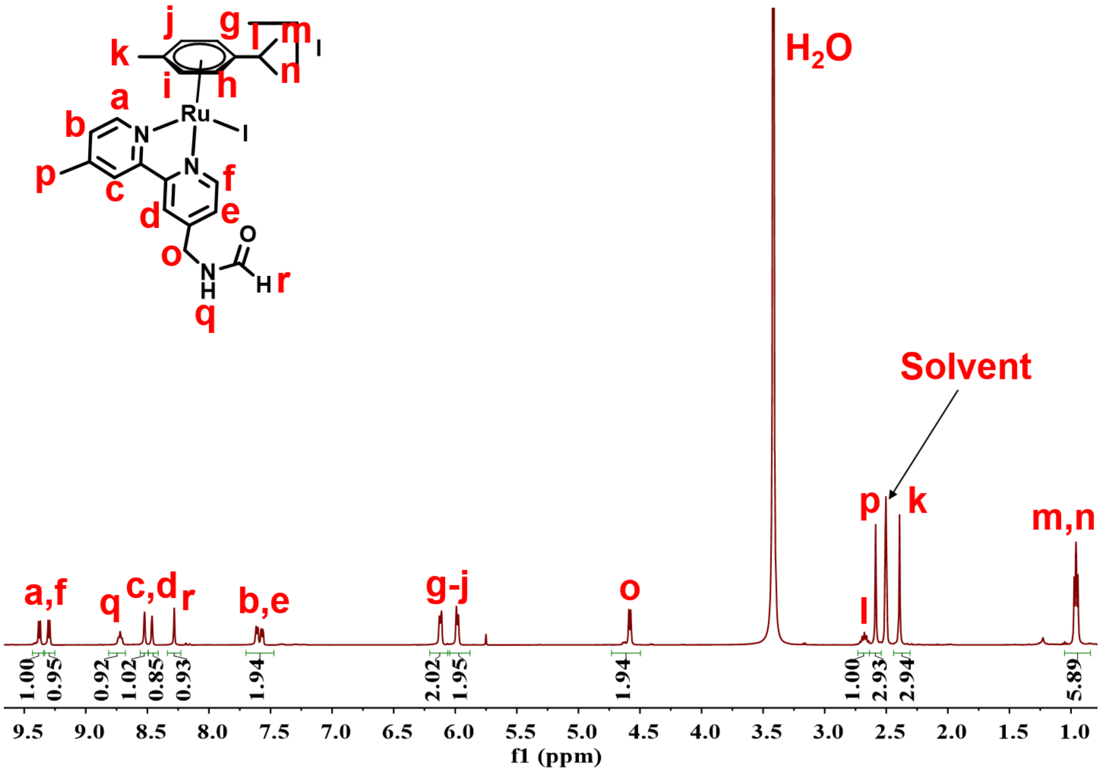


Supplementary Figure 49. $^{1}H$ NMR spectrum (400 MHz, $d_6$-DMSO) of **RuF**.

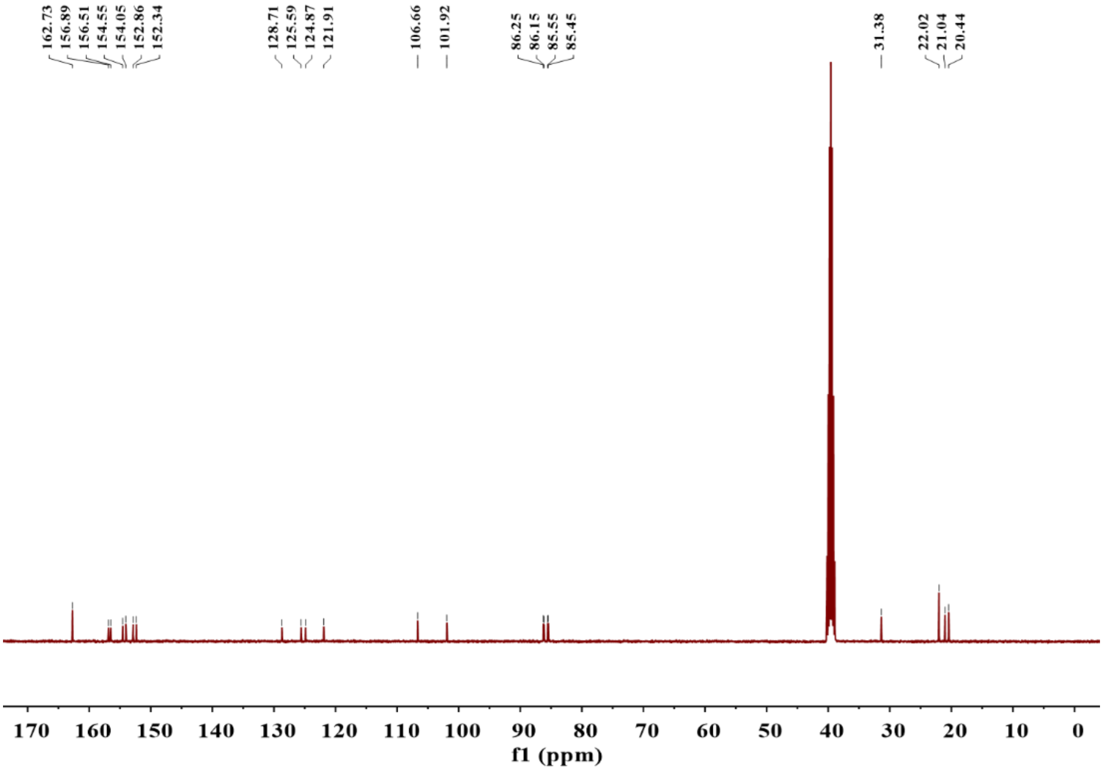


Supplementary Figure 50. $^{13}C$ NMR spectrum (400 MHz, $d_6$-DMSO) of **RuF**.

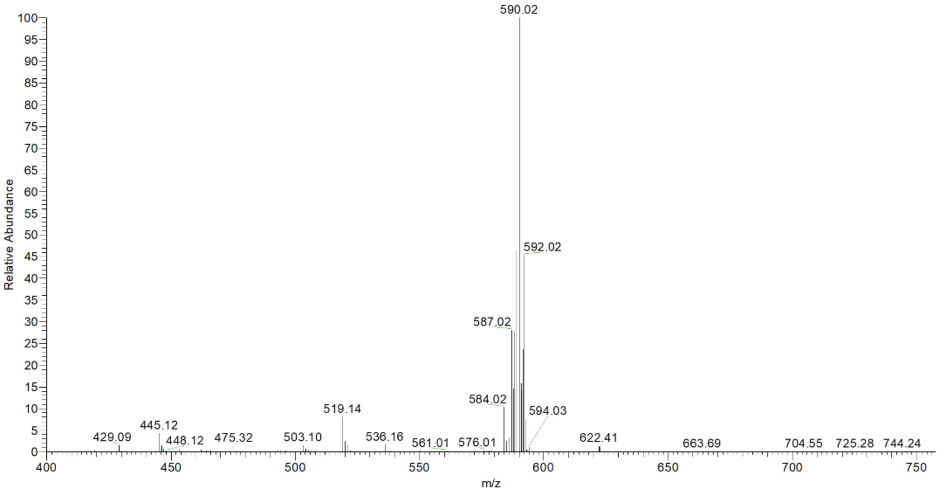


Supplementary Figure 51. ESI-MS spectrum of **RuF** in $CH_3OH$. (calcd for [**RuF**-$I^-$]$^+$ m/z = 589.46, found m/z = 590.02.)

Supplementary Table 1. MIC values of compounds of interest.

| Strains | **RuR1** | **RuR2** | **RuR3** | **RuN** | **RuC** | **RuB** | **RuF** | $Ru^{II}$-arene | Rhein | RFP | INH | EMB | KAN |
|---|---|---|---|---|---|---|---|---|---|---|---|---|---|
| Ms | 2 | 2 | 2 | 8 | 16 | 24 | >256 | 128 | >256 | 8 | 4 | 0.3 | 0.5 |
| Mf | 8 | 32 | 8 | 8 | 32 | 64 | >64 | >64 | >256 | 4 | 1 | 6 | 4 |
| Mm | 8 | 32 | 2 | 24 | 32 | 64 | 64 | >64 | >256 | 1 | >128 | >128 | 1 |
| H37Rv | 4 | 8 | 1 | 16 | 32 | >32 | >32 | >64 | 64 | 0.002 | <0.03 | 2 | 2 |

Ms: *Mycobacterium smegmatis*, Mf: *Mycobacterium fortuitous*, Mm: *Mycobacterium marinum*.

Supplementary Table 2. Raw data for heatmap of DNA binding affinity for compounds of interest.

| | Compounds | **RuR3** | Daunorubicin | EB | Hoechst | Mitomycin | RAS | RuZ1 | $[Ru(phen)_3]Cl_2$ | $Ru^{II}$-arene | Rhein |
|---|---|---|---|---|---|---|---|---|---|---|---|
| | $K_i$ (MGB), nM | 6.4 | 0.62 | 18 | 2.5 | 80 | 15 | 17 | 14 | >160 | 20 |
| | $K_i$ (Intercalation), nM | 540 | NT | 100 | >3200 | >3200 | 696 | 1262 | NT | >6400 | >6400 |
| DNA retardation | Plasmid DNA | 0 | 76 | NT | 3.8 | 96 | 67 | 69 | 79 | 92 | 89 |
| | Sheared DNA | 4.5 | 41 | NT | 33 | 92 | 47 | 60 | 99 | 46 | 93 |
| | FAM-DNA | 35 | 83 | 94 | 38 | 96 | 87 | 95 | 96 | 93 | 98 |

NT: not tested. Data not tested were indicated with a black dot on the heatmap in Extended Data Fig. 2d.
